# Phylogenetics of a Rapid, Continental Radiation: Diversification, Biogeography, and Circumscription of the Beardtongues (*Penstemon*; Plantaginaceae)

**DOI:** 10.1101/2021.04.20.440652

**Authors:** Andrea D. Wolfe, Paul D. Blischak, Laura S. Kubatko

## Abstract

*Penstemon* (Plantaginaceae), the largest genus of plants native to North America, represents a recent continental evolutionary radiation. We investigated patterns of diversification, phylogenetic relationships, and biogeography, and determined the age of the lineage using 43 nuclear gene loci. We also assessed the current taxonomic circumscription of the ca. 285 species by developing a phylogenetic taxonomic bootstrap method. *Penstemon* originated during the Pliocene/Pleistocene transition. Patterns of diversification and biogeography are associated with glaciation cycles during the Pleistocene, with the bulk of diversification occurring from 1.0–0.5 mya. The radiation across the North American continent tracks the advance and retreat of major and minor glaciation cycles during the past 2.5 million years with founder-event speciation contributing the most to diversification of *Penstemon*. Our taxonomic bootstrap analyses suggest the current circumscription of the genus is in need of revision. We propose rearrangement of subgenera, sections, and subsections based on our phylogenetic results. Given the young age and broad distribution of *Penstemon* across North America, it offers an excellent system for studying a rapid evolutionary radiation in a continental setting.

---

*Penstemon* Schmidel (Plantaginaceae), commonly known as the beardtongues, is the largest plant genus endemic to the North American continent, containing ca. 285 species, with new taxa being described every few years (Turner 2010; Estes 2012; O’Kane and Heil 2014; Zacarías-Correa et al. 2019). The recent publication of the Flora of North America treatment for the genus (Freeman 2019) included 239 species found north of the US/Mexico border. Mexico has ca. 60 species (Zacarías-Correa 2020). This large and diverse genus is an example of a rapid evolutionary radiation, with much of the diversification hypothesized to have taken place during the Pleistocene (Wolfe et al. 2006). Most of the species are found in the Intermountain West (Holmgren 1984; Holmgren and Holmgren 2016), with the number of species decreasing dramatically east of the Rocky Mountains. For example, the Great Plains region to the Mississippi River contain ca. 39 species, with an additional 10 species found only east of the Mississippi (Lindgren and Wilde 2003). In contrast, Utah has 76 species, with ca. 29 percent endemic to the state (Holmgren 1984; Stevens et al. 2020).

Species of *Penstemon* occur in a wide variety of habitats, including edaphic specialization (e.g., deep sand, limestone derived soils, igneous soils, oil shales), with most species adapted to xeric landscapes (Lindgren and Wilde 2003; Dockter et al. 2013). The diversity of floral shapes, sizes, and colors suggests a significant role of pollinator selective pressure in the evolutionary history of the genus (Pennell 1935; Straw 1956a, 1956b, 1963). Other morphological traits with high levels of variation include size, habit, inflorescence, leaves, anthers, and staminodes. Most of the species are restricted to narrow ranges, with about 1/3 of the species restricted to a single state in the United States or Mexico. Habitat specialization, combined with narrow geographic distribution, has resulted in conservation concerns for many species (Wolfe et al. 2014, 2016; Rodriguez-Peña et al. 2018; Stone et al. 2019; Zacarías-Correa et al. 2020). *Penstemon penlandii*, *P. haydenii,* and *P. debilis* have been Federally listed as endangered or threatened under the US Endangered Species Act (USFWS 2011), with more than 90 species under consideration for protection (USFWS 1993).

The taxonomy of the genus has been developed over the past 250 years, since its recognized designation by Schmidel (1763). Earlier work included a description by John Mitchell, which was published in 1748 (Straw 1966), but his work is not currently recognized as establishing the genus (Freeman 2019). The most comprehensive listings of *Penstemon* taxonomy have been compiled by the American Penstemon Society in a series of publications that were most recently updated nearly 20 years ago (Lodewick and Lodewick 1987; Lindgren and Wilde 2003). Because the Flora of North America treatment by Freeman (2019) did not address the taxonomy of the genus, we will refer to the taxonomic designations from Lindgren and Wilde (2003), amended with information from the updated *Penstemon* treatments in Volumes Four and Seven of *The Intermountain Flora* (Holmgren 1984; Holmgren and Holmgren 2016).

The most comprehensive phylogenetic study for *Penstemon* was published by Wolfe et al. (2006) and included 163 species representing all subgenera and sections of the genus. This study used ITS and two cpDNA loci and was able to establish some major trends in the evolutionary patterns for the genus such as 1) the rapid evolutionary radiation of a large genus in a continental setting, 2) biogeographic trends across North America, 3) a probable role in diversification due to hybridization, and 4) the independent derivation of a hummingbird pollination syndrome in at least 10 lineages. Taxonomic trends observed in the 2006 study included strong support for the circumscription of subgenus *Dasanthera* as the earliest diverging lineage, and close affinities among some sections of subgenera *Saccanthera* and *Penstemon*, and *Penstemon* and *Habroanthus*. The resolution of the backbone for the tree was insufficient to fully resolve taxonomic relationships, but indications for non-monophyly for three of the four multi-taxa subgenera were present.

Recent phylogenetic studies for *Penstemon* have focused on the core group of species encompassing sections *Gentianoides*, *Coerulei*, *Spectabiles,* and members of what has traditionally been known as subgenus *Habroanthus* (Crosswhite 1967; Wessinger et al. 2016, 2019). The Wessinger et al. (2016, 2019) studies employed multiplexed shotgun genotyping (MSG) SNP data for 75 and 120 species, respectively. The phylogenies based on MSG data had more resolution than did the trees presented in the Wolfe et al. (2006) study but were mostly in agreement with clade topologies of the earlier tree, and the trees were specifically used as a framework for understanding the evolutionary trends in the shift between bee- to bird-pollination syndromes.

In the Flora of North America, including most of the species of *Penstemon*, Freeman (2019) revised the taxonomy of genus to include only two subgenera (*Dasanthera* and *Penstemon*), based on the phylogenetic results from Wolfe et al. (2006). In addition to subsuming four subgenera, the number of sections was reduced, and there were no divisions below the sectional level. This resulted in the small subgenus *Dasanthera* circumscribing only nine species including *P. personatus*, formerly placed in a monotypic subgenus (*Cryptostemon*), and the remainder of the genus circumscribing subgenus *Penstemon*.

In this study we expand the phylogenetic survey to include 239 species of *Penstemon* with the intent to examine the correspondence of historical and contemporary taxonomic boundaries. With the expanded phylogenetic survey, we also address the age of the genus, patterns of diversification, and biogeographic trends, and we compare our results to previous phylogenetic surveys for *Penstemon*.

## Materials and Methods

### Sample Collection, DNA Extraction, and Amplicon Sequencing

DNA was extracted from a combination of either field-collected, silica-dried leaf tissue or leaves sampled from herbarium specimens using a modified CTAB protocol for DNA isolation (Wolfe 2005). After extraction, all samples were quantified using a Qubit fluorometer (Invitrogen, Carlsbad, CA, USA) and normalized to a concentration of 20 ng/μL. Normalized DNA samples for the 282 accessions of *Penstemon* representing 239 species plus two hybrids (ca. 84% of the genus) used in this study, plus nine accessions from other members of the tribe Cheloneae (one *Pennelianthus*, five *Keckiella*, one *Nothochelone*, one *Chelone*, and one *Chionophila*), were sent to the IBEST Genomics Resource Core at the University of Idaho (Moscow, ID, USA) for sample preparation and amplicon sequencing (Appendix 1). All subgenera, sections, and subsections of *Penstemon* were represented in the sampling (Table 1). Amplification of targeted regions and the addition of sample barcodes and Illumina adapters was done using microfluidic PCR on the Fluidigm 48×48 Access Array (Fluidigm Corporation, South San Francisco, CA, USA), followed by 300 bp, paired-end sequencing on an Illumina MiSeq (Illumina, San Diego, CA, USA) (Uribe-Convers et al. 2016). Primers for the 48 loci used in this study were designed and tested as described in Blischak et al. (2014) and are given in Table S1. Raw, paired-end sequencing reads were demultiplexed with dbcAmplicons [https://github.com/msettles/dbcAmplicons] prior to being returned from IBEST (Uribe-Convers et al. 2016). We then processed the sequence data using Fluidigm2PURC v0.2.1 [https://github.com/pblischak/fluidigm2purc] (Blischak et al. 2018) to assemble and align haplotypes for phylogenetic inference (Rothfels et al. 2017). The initial, unfiltered haplotype data returned by Fluidigm2PURC was further processed in Geneious v8.1.9 (Kearse et al. 2012). Each locus was first realigned in Geneious using MUSCLE with default settings (Edgar 2004), after which we removed poorly aligned sequences, as well as a small number of duplicated accessions that were repeated across sequencing runs.

**Table 1.** Taxonomy of *Penstemon* as recognized by the American Penstemon Society, with annotations from the Freeman (2019) Flora of North America Treatment. Sampling of species for this study is indicated in the numerator, and abbreviations are those used in Figure 1.

| Subgenus | Section | Subsection | Abbreviations | Species Sampling | FNA designations |
| --- | --- | --- | --- | --- | --- |
| *Cryptostemon* |  |  | C | 1/1 | Subg. Dasanthera |
| *Dasanthera* |  |  | D | 9/9 |  |
| *Dissecti* |  |  | Di | 1/1 |  |
| *Habroanthus* | *Elmigera* |  | H, El | 6/7 |  |
|  | *Glabri* |  | H, Gl | 34/44 |  |
| *Penstemon* | *Ambigui* |  | P, A | 1/2 |  |
|  | *Baccharifolii* |  | P, B | 1/1 |  |
|  | *Chamaeleon* |  | P, Ch | 3/4 |  |
|  | *Coerulei* |  | P, Co | 20/20 |  |
|  | *Cristati* |  | P, Cr | 24/28 | 3 spp transferred from sect. *Ericopsis* |
|  | *Ericopsis* | *Caespitosi* | P, E, Ca | 12/12 | sect. *Caespitosi* (2 spp transferred to sect. *Cristati*) |
|  |  | *Ericopsis* | P, E, E | 1/1 | sect. *Cristati* |
|  |  | *Linarioides* | P, E, L | 3/3 | sect. *Caespitosi* |
|  | *Fasciculus* | *Campanulati* | P, F, Cp | 5/8 |  |
|  |  | *Fasciculi* | P, F, Fa | 10/13 |  |
|  |  | *Perfoliati* | P, F, Pf | 1/3 |  |
|  |  | *Racemosi* | P, F, R | 4/4 |  |
|  | *Peltanthera* | *Centranthifolii* | P, Pe, Ce | 10/10 | sect. *Gentianoides* |
|  |  | *Havardiani* | P, Pe, Hv | 1/3 |  |
|  |  | *Peltanthera (Spectabiles)* | P, Pe, Sp | 14/16 | sect. *Spectabiles* |
|  |  | *Petiolati* | P, Pe, Pt | 1/1 | sect. *Petiolati* |
|  | *Penstemon* | *Arenarii* | P, P, Ar | 1/2 | sect. *Penstemon* |
|  |  | *Deusti* | P, P, De | 3/3 |  |
|  |  | *Gairdneriani* | P, P, G | 2/2 |  |
|  |  | *Harbouriani* | P, P, Ha | 1/1 |  |
|  |  | *Humiles* | P, P, Hu | 15/20 |  |
|  |  | *Multiflori* | P, P, M | 1/1 |  |
|  |  | *Penstemon* | P, P, P | 10/18 |  |
|  |  | *Proceri* | P, P, Pr | 16/17 |  |
|  |  | *Tubaeflori* | P, P, T | 1/1 |  |
| *Saccanthera* | *Bridgesiani* |  | S, B | 1/1 |  |
|  | *Saccanthera* | *Heterophylli* | S, S, He | 20/22 | sect. *Saccanthera* |
|  |  | *Serrulati* | S, S, Se | 6/6 | Sect. *Saccanthera* |

Total = 239/285

**Figure 1.**
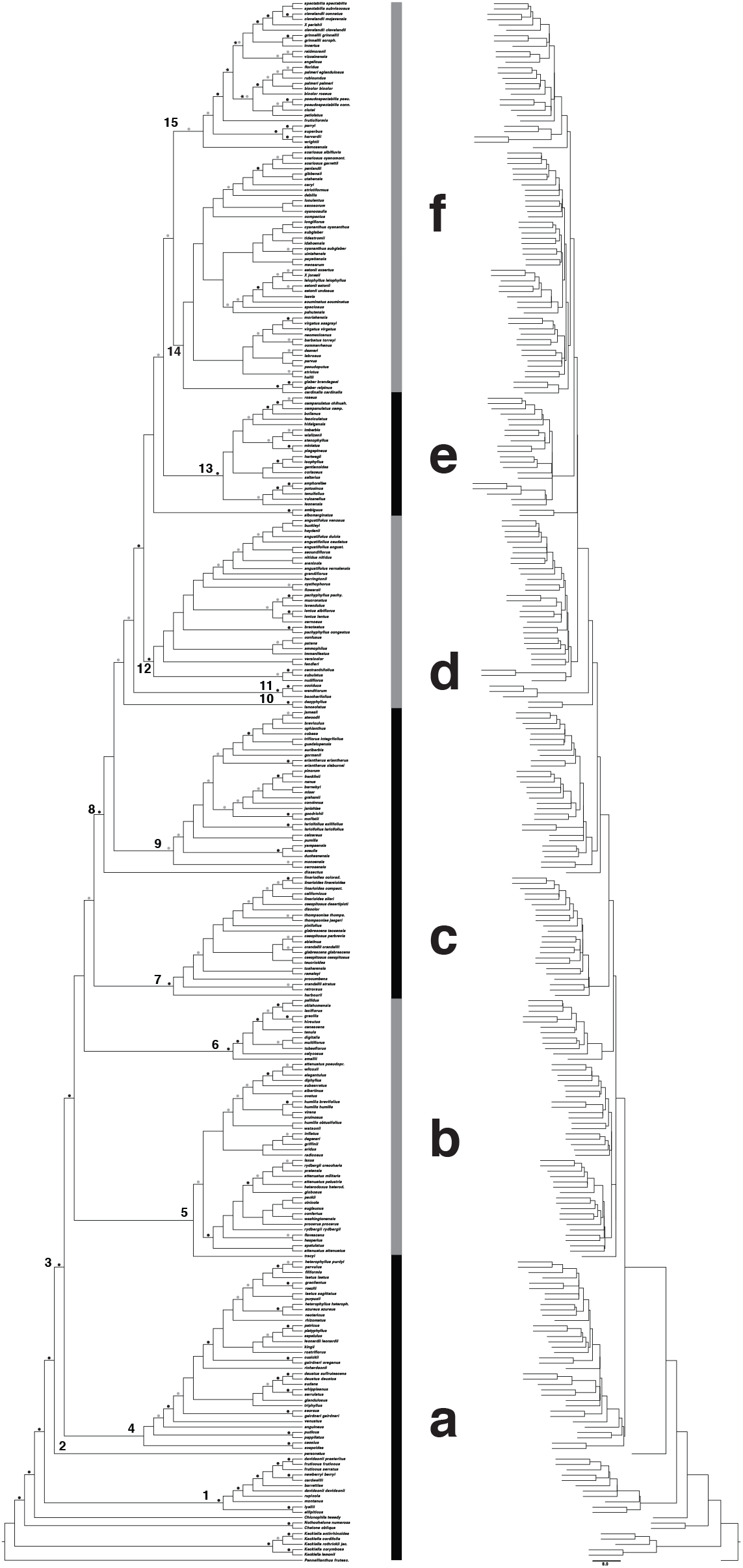
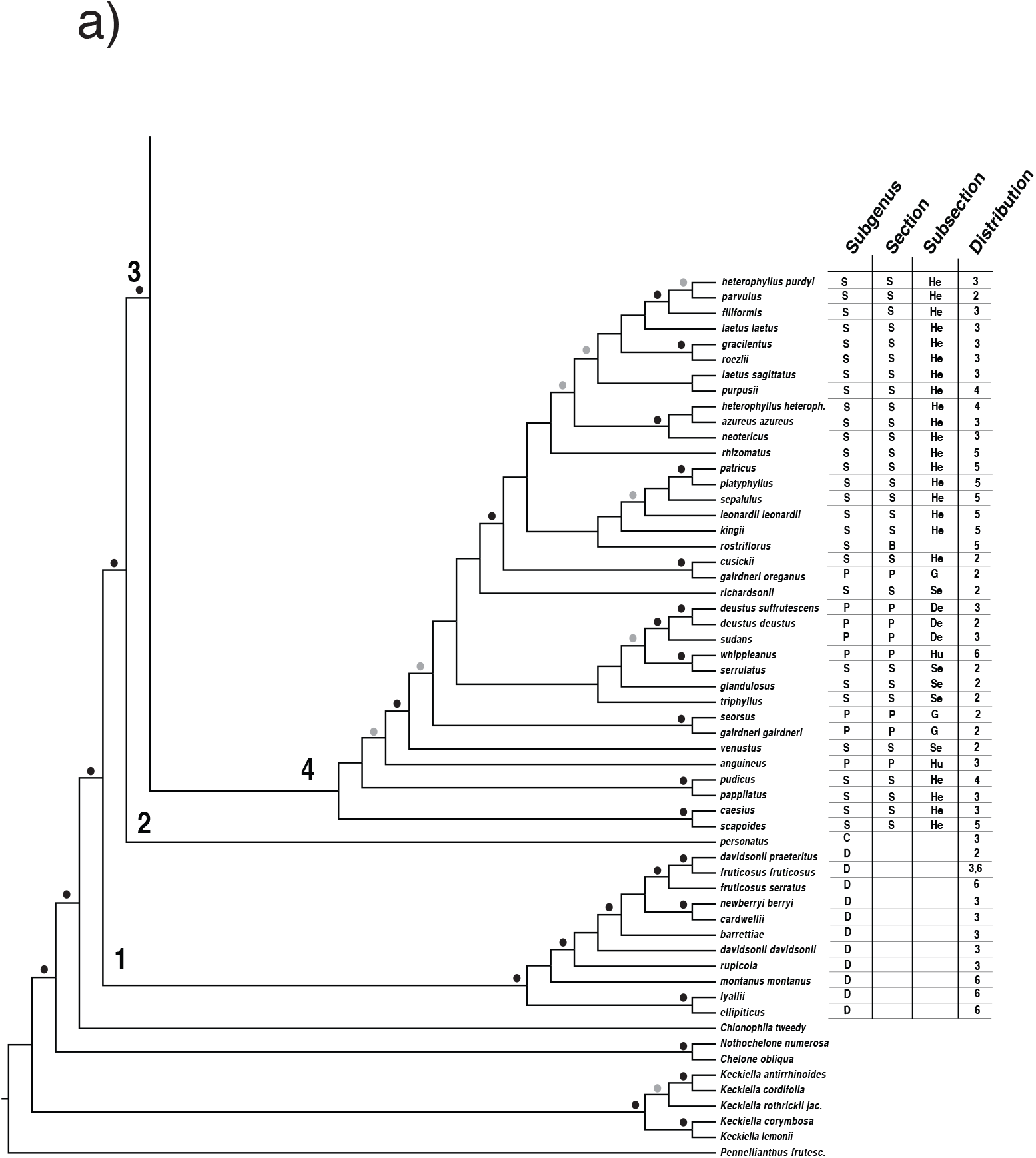
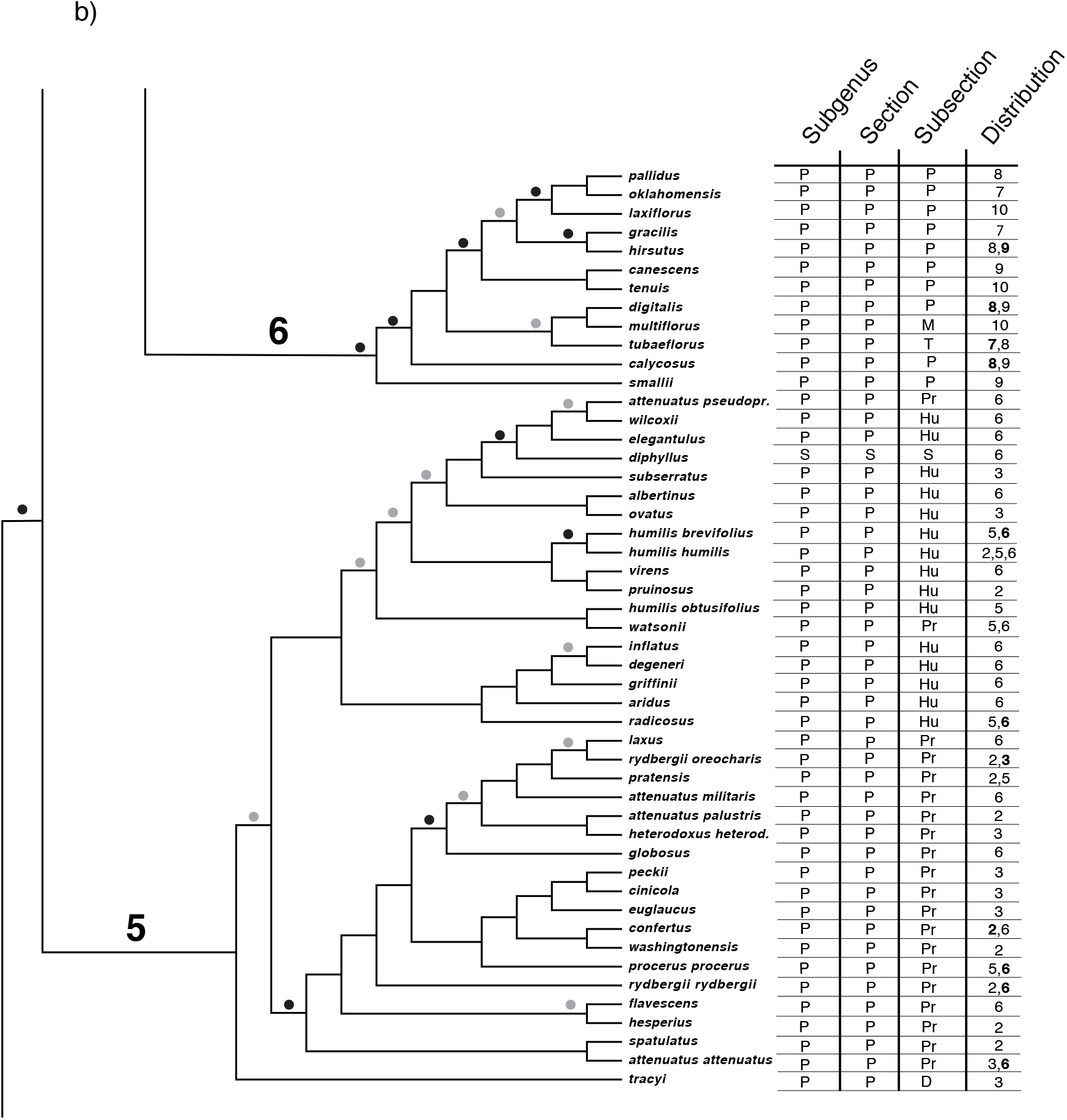
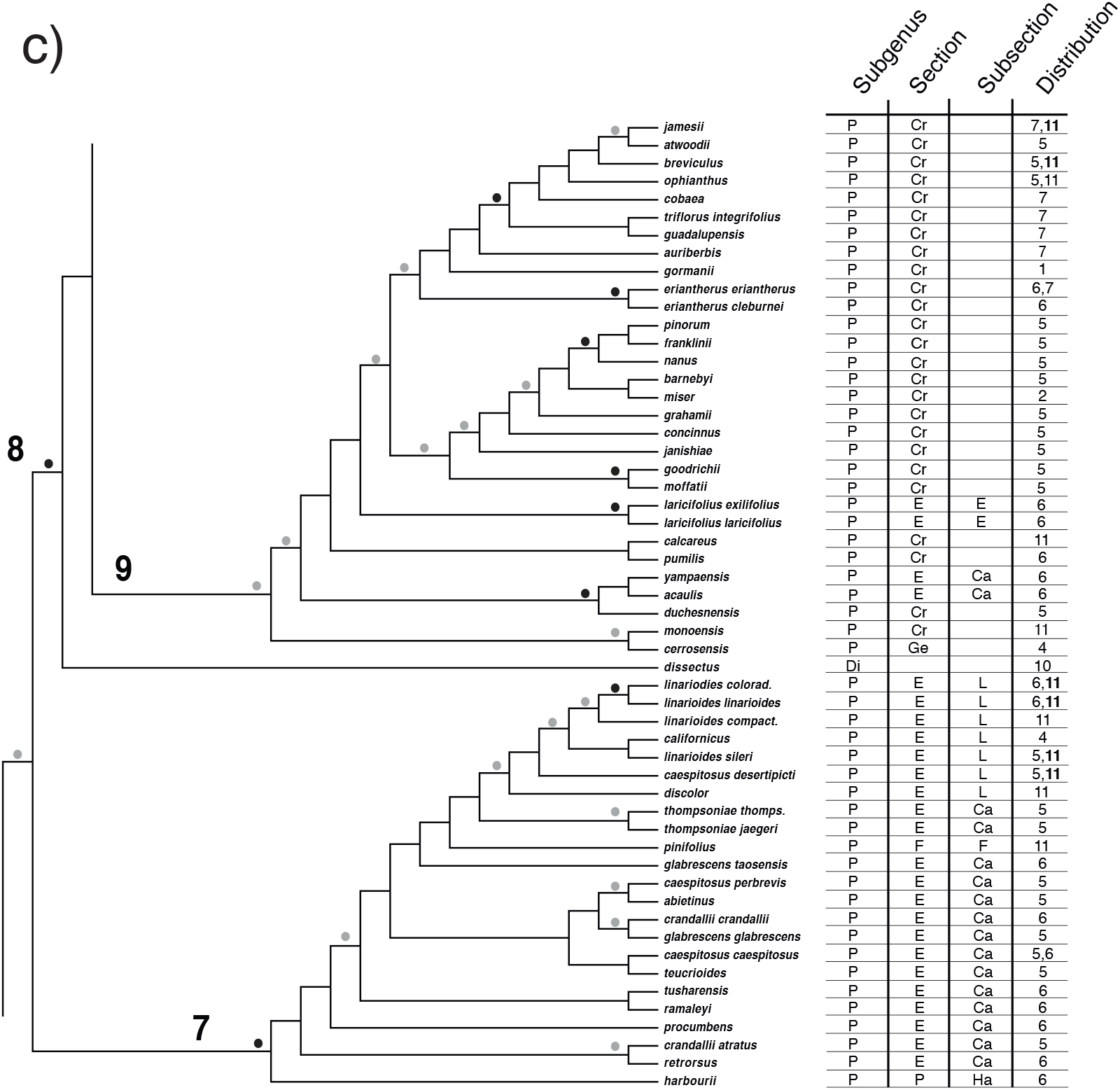
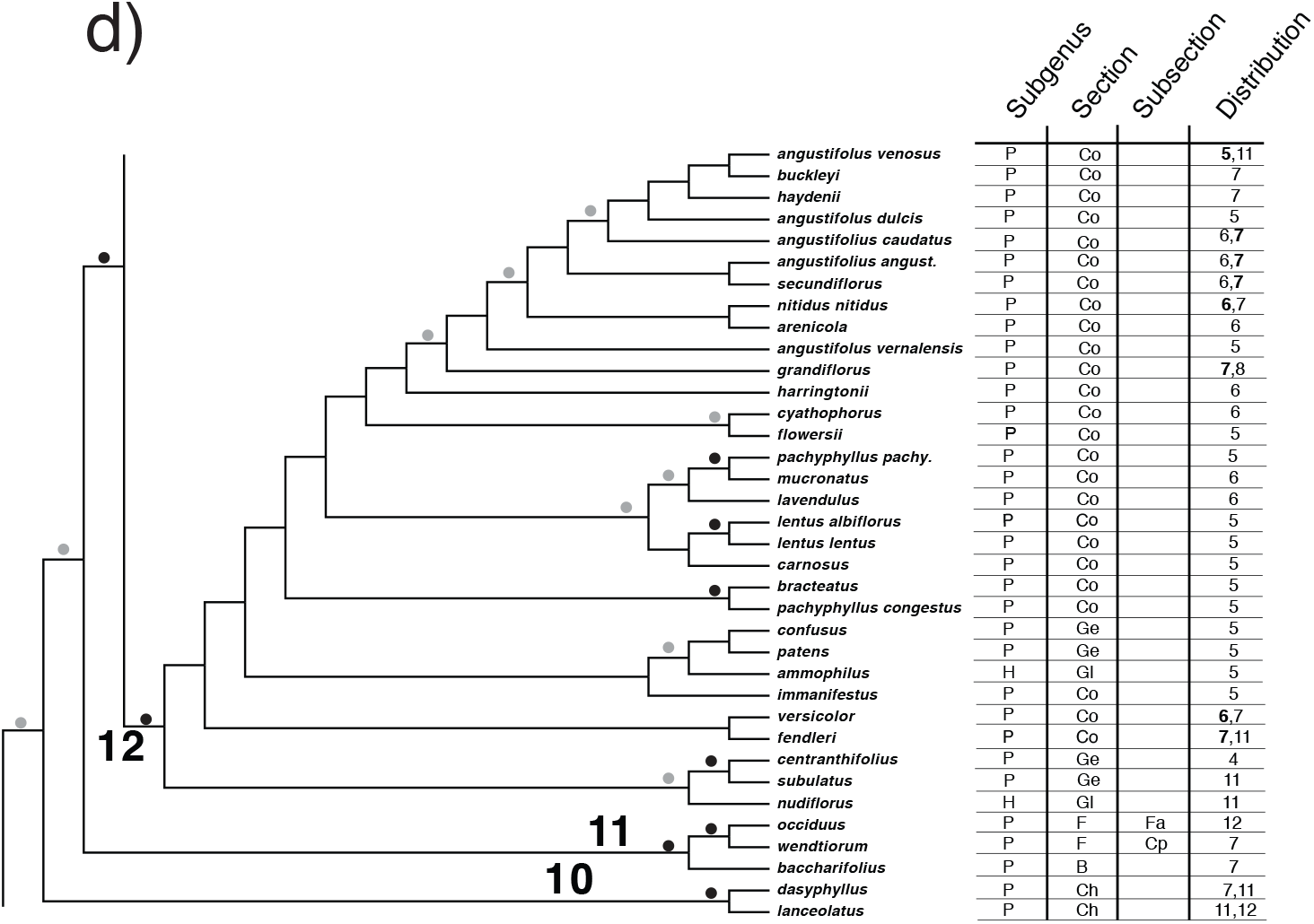
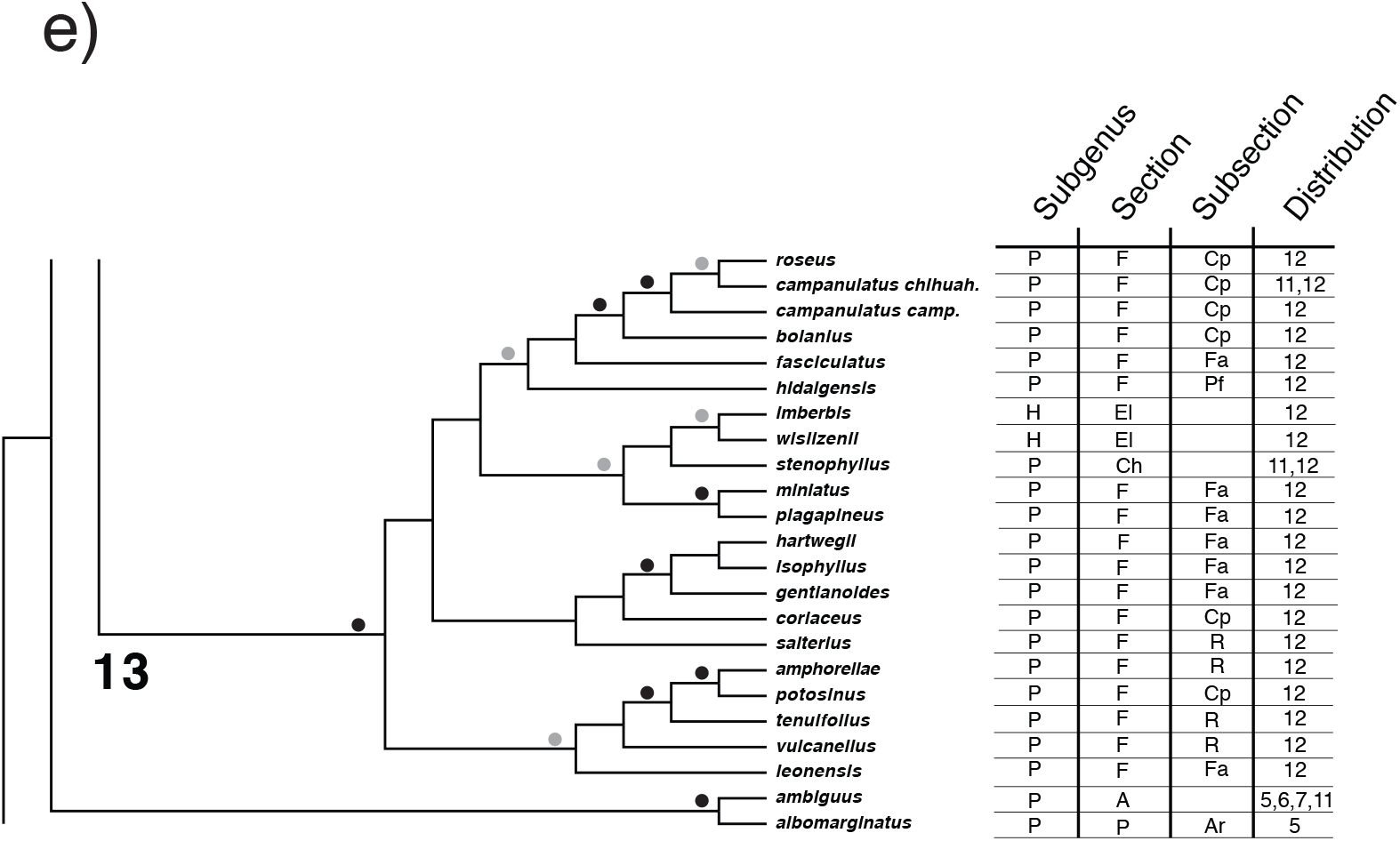
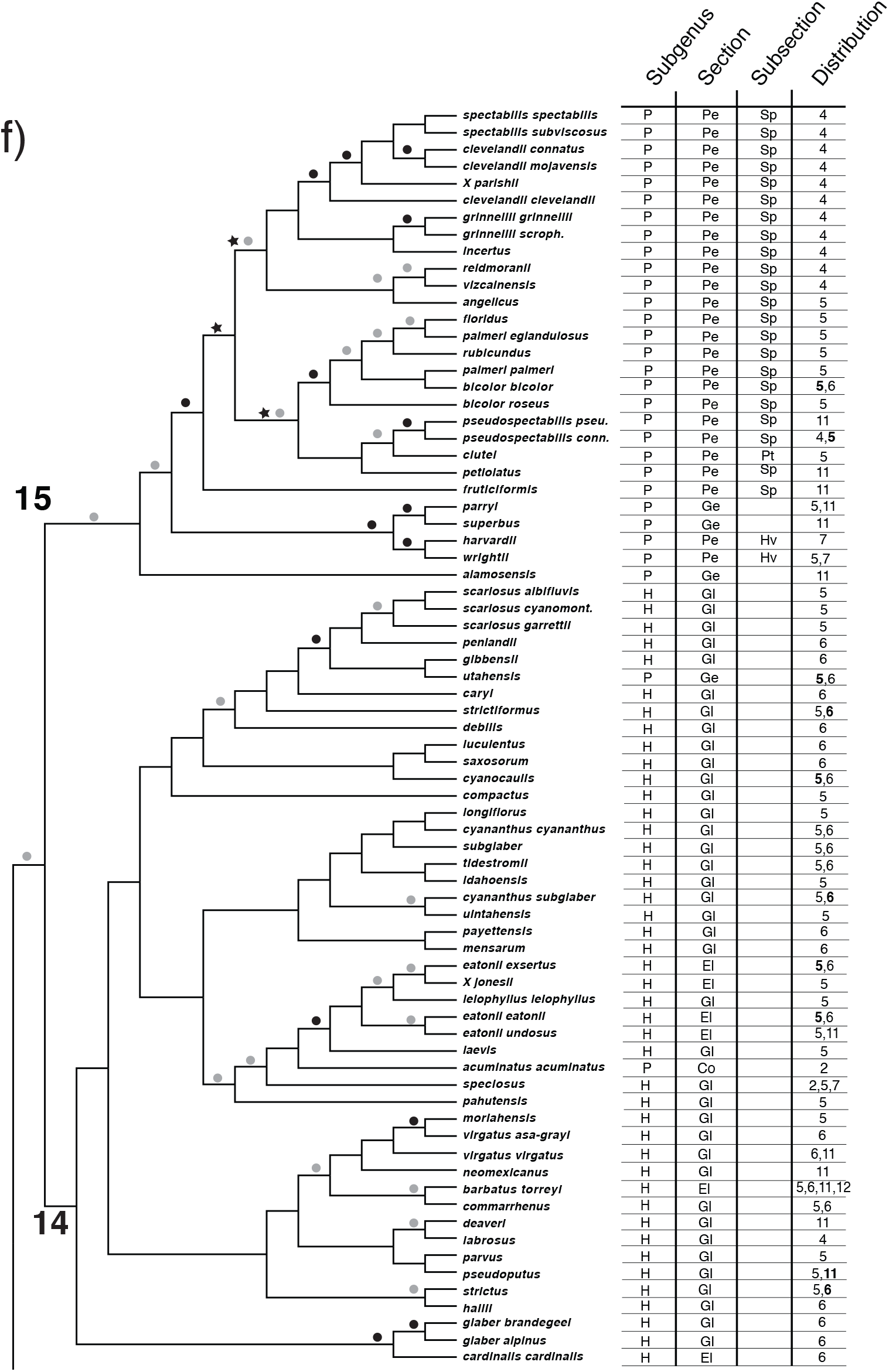
Phylogeny of *Penstemon* and related members of Cheloneae based on the Astral analysis. Relative branch lengths are shown on the right. The vertical bar with lettering refers to insets 1a–1f, which show details of the phylogenetic tree.

### Phylogenetic Inference

To infer a phylogeny for *Penstemon*, we used a coalescent-based approach. We inferred individual gene trees for each locus using FastTree v2.1.11 (Price et al. 2010), which we then used as input for ASTRAL v5.6.3 (ASTRAL-III; Zhang et al. 2018). Gene trees were inferred using the GTR+GAMMA model and ASTRAL-III was run using default options. We also ran a concatenation-based, maximum likelihood (ML) analysis using RAxML v8.2.10 (Stamatakis 2014). For our ML analysis, we concatenated all processed and aligned sequences using Phyutility v2.7.1 (Smith and Dunn 2008), fit separate GTR+GAMMA parameters for each gene using a partitions file, and performed 1000 rounds of rapid bootstrapping to assess support (Stamatakis et al. 2008). Previous phylogenetic work has placed *Pennellianthus frutescens* as sister to the rest of the Cheloneae, and we used it here as an outgroup for both analyses (Wolfe et al. 2002, 2006).

### Taxonomic Bootstrap

To assess support for the current taxonomic classification of *Penstemon* at the level of subgenus, section, and subsection, we performed bootstrap resampling of taxa from within their respective taxonomic ranks. For an individual bootstrap replicate, a single representative was sampled from each named group at the chosen taxonomic rank and a phylogeny was inferred using these individuals’ sequence data. At each taxonomic level (subgenus, section, and subsection), we performed 1000 rounds of bootstrap resampling with phylogenetic inference conducted using SVDQuartets (Chifman and Kubatko 2014) on each bootstrapped data set. Replicates were then combined into a majority rule extended consensus tree using RAxML v8.2.10 (Stamatakis 2014). The Python script for performing this ‘taxonomic bootstrapping’ is available in Supplemental Materials on Dryad (*sample_from_taxonomy.py*).

### Time Calibration and Analysis of Diversification Rates

To obtain a time-calibrated phylogeny, we first used BEAST v2.6.0 with fossils for previously inferred divergence events for families within Lamiales from the literature as calibration points to date the splits between lineages in the tribe Cheloneae (Vargas et al. 2014). To do this, we downloaded publicly available sequences for the ITS region for three pairs of species with fossils supporting their minimum divergence time: *Kigelia africana* (Bignoniaceae) + *Catalpa duclouxii* (Bignoniaceae) [28.4 mya], *Veronica persica* (Plantaginaceae) + *Plantago lanceolata* (Plantaginaceae) [11.6 mya], and *Gratiola neglecta* (Plantaginaceae) + *Bacopa eisenii* (Plantaginaceae) [5.3 mya]. We then combined the sequences from these six species with the ITS region for samples from Cheloneae (Wolfe et al. 2002, 2006). In addition, we included sequences for trnC-D and trnT-L for all included Cheloneae samples (Table S1). Each locus was aligned with MAFFT (Katoh 2013) using the ‘--auto’ option and then all genes were concatenated with Phyutility (Smith and Dunn 2008).

Program settings for the analysis with BEAST were specified within BEAUti (Bouckaert et al. 2019). Each locus was given an independent substitution model (GTR + Gamma with four rate categories) and clock model (relaxed log-normal; Drummond et al. 2006) but were constrained to all follow a single tree topology. For the three fossil calibrations, we used log-normal distributions with the following ranges: mean = 3.346 and standard deviation = 0.2 (95% quantile range: 20.4–39.4 my) for *K. africana* + *C. duclouxii*; mean = 2.45 and standard deviation = 0.2 (95% quantile range: 8.34–16.1 my) for *V. persica* + *P. lanceolata*; and mean = 1.667 and standard deviation = 0.2 (95% quantile range: 3.81–7.36 my) for *G. neglecta* + *B. eisnii*. All other prior specifications were left at their default options. Parameters were sampled for 2 x 10^7^ generations and were logged every 1,000 generations. Posterior samples were examined in Tracer v1.7.1 (Rambaut et al. 2018) to assess convergence and to estimate effective sample sizes (ESS). Results were then summarized into a maximum clade credibility (MCC) tree using TreeAnnotator after discarding 25% of the samples as burn in.

After estimating divergence times for the major Cheloneae lineages with BEAST, we used three of the 95% highest posterior density estimates as secondary calibration bounds to date the entire 293 taxon RAxML phylogeny using treePL (Smith and O’Meara 2012). The secondary calibration points were defined for the most recent common ancestors for *Pennellinathus frutenscens* + *Chelone glabra* (4.375–19.04 my), *K. breviflora* + *P. montanus* (1.89–8.539 my), and *P. montanus* + *P. personatus* (1.419–6.416 my). We first ran treePL with the ‘prime’ option to find the best optimization settings. Then, we used these settings, along with the ‘thorough’ option, to estimate divergence times across the tree.

Using this time-calibrated phylogeny, we also estimated diversification rates with BAMM v2.5.0 (Rabasky 2014). Priors for the expectedNumberOfShifts, lambdaInitPrior, lambdaShiftPrior, and muInitPrior distributions were set using the setBAMMprior function in the BAMMtools R package v2.1.7 (Rabosky et al. 2014). All other priors were left at their default value. We then sampled parameters for 1 x 10^6^ generations, logging them every 1,000 generations. After discarding 10% of the samples as burn-in, the remaining posterior samples were then processed in R v3.6.1 (R Core Team 2019) using the coda package (Plummer et al. 2006) to assess convergence and the BAMMtools package to summarize estimated rate shifts, net diversification rates, and sampled placements of shifts on the phylogeny. We also estimated diversification dynamics using MEDUSA v1.41 (Alfaro et al. 2009), implemented in the R package geiger v2.0.6.4 (Pennell et al. 2014), assuming a birth-death model.

### Analysis of Biogeographic Distribution

To estimate biogeographic patterns in *Penstemon*, we used BioGeoBEARS v1.1.2 (Matzke 2013a) to infer ancestral areas as well as to compare competing biogeographic hypotheses using model selection criteria (e.g., Akaike information criterion, AIC; Akaike 1974). Within BioGeoBEARS, we fit the DIVALIKE (Ronquist 1997), DEC (Ree and Smith 2008), and BAYAREALIKE (Landis et al. 2013) models, along with their jump dispersal (“+J”) counterparts, for a total of six models (Matzke 2013b, 2014). Areas were defined initially by the 12 regions from Wolfe et al. (2006) with updates based on the physiographic regions of North America (Fig. S1; Table 2; Barton et al. 2003). These were split into three separate sets to help with computational feasibility and to reflect geographic proximity of regions. The first set of analyses included areas [1-6], the second set included areas [4-7,11,12], and the third set included areas [6-11]. The maximum number of ancestral areas for each analysis was set as the maximum number of areas currently occupied by the sampled taxa in a given set and all other parameters were kept at their default values. R code for running these analyses is available on dryad.

**Table 2.** Biogeographic regions for *Penstemon* (modified from Wolfe et al. 2006) used in Figure 1, and for BioGeoBears analyses.

| Region Description | Designation |
| --- | --- |
| Intermontane Plateau – extreme north | 1 |
| Pacific Northwest, exclusive of Eastern and Western Cordilleras | 2 |
| Cascade-Sierra and Northwestern Cordillera | 3 |
| Southwestern Cordillera | 4 |
| Intermountain Region | 5 |
| Eastern Cordillera | 6 |
| Great Plains | 7 |
| Interior Lowlands | 8 |
| Appalachian Mountain System | 9 |
| Coastal Plains | 10 |
| American Southwest | 11 |
| Mexican Highlands | 12 |

To further dissect evolutionary pattern and process for *Penstemon*, we also downloaded distribution information for every species of *Penstemon* available in the SEINet data portal (http://www.swbiodiversity.org). Only Federally- or State-listed species were excluded from our data set because these locations are restricted from public access. This information was collected as Keyhole Markup Language (KML) files which were then imported into Google Earth Pro (https://www.google.com/earth/versions/#earth-pro). Using our time-calibrated phylogeny, we plotted two patterns in Google Earth Pro: 1) the individual patterns of distribution of taxa from the earliest to most recently diverged lineages, and 2) a collective distribution map based on time of divergence estimations for the terminal lineages in each clade (Table S2). These time slices were 2.5 mya, 2.0 mya, 1.8 mya, and forward to 0.1 mya in 100,000-year increments. The time of divergence was rounded up or down to fit into a time increment. Animations of the distributions in these two map sets were assembled from the individual or collective map images. The pattern of these species’ distributions was then examined in the context of inferred glaciation cycles during the Pleistocene (Ehlers and Gibbard 2007; Ehlers et al. 2018).

## Results

### Amplicon Sequencing

Of the 48 loci targeted for sequencing, 43 were successfully amplified in a sufficient number of our samples and were used for downstream analyses (Table S1; see filtering criteria in the Appendix 2 in Supplement Materials). Our final data matrix contained 17,518 sites (7171 variable, 4049 parsimony informative), 293 taxa (282 from *Penstemon*), and 30.4% missing data. The average number of non-missing sites per taxon was 12,191.2.

### Phylogenetic Inference

Tribe Cheloneae is circumscribed by the genera *Pennellianthus, Collinsia, Keckiella, Chelone, Nothochelone, Chionophila,* and *Penstemon*. Our phylogeny based on 43 nuclear genes recovered a topology for the tribe consistent with earlier studies (Fig. 1; Wolfe et al. 2002, 2006), but with higher relative nodal support throughout the tree, and greater resolution within clades and along the backbone. Relationships among early diverging lineages in *Cheloneae* in this study also agree with previous work and support *Penstemon* as monophyletic with *Chionophila tweedyi* as sister to the entire genus (prior >0.8) (Fig. 1a).

Within *Penstemon*, we infer subgenus *Dasanthera* as monophyletic, as has also been found previously (Wolfe et al. 2002, 2006). Major differences between the phylogeny presented here and earlier studies include the well-resolved placement of subgenus *Dasanthera* as sister to the rest of the genus (Fig.1a position 1), the position of *P. personatus* (subg. *Cryptostemon*; Fig. 1a position 2) as the next diverging lineage, a strongly supported clade containing the rest of the genus (referred to below as the crown clade; Fig 1a position 3), strong support for the majority of the group of penstemons classified as subgenus *Saccanthera* (Fig. 1a position 4; Freeman 2019), affinities for sect. *Penstemon* subsections *Proceri* and *Humiles* (Fig. 1b position 5), strong support for sect. *Penstemon* subsect. *Penstemon* to include subsections *Multiflori* and *Tubaeflori* (Fig. 1b position 6), and strong support for the majority of taxa traditionally in sect. *Ericopsis* (Fig. 1c position 7). *Penstemon dissectus* is the first diverging lineage for remaining group of penstemons (Fig. 1c position 8). Additional differences are the moderate to strong support for much of the backbone for the clade of penstemons sister to *P. dissectus* (Figs. 1c–1f). A clade consisting of section *Coerulei* and species from sections *Gentianoides* and *Glabri* has strong nodal support (Fig. 1d position 12). There is strong support for section *Fasciculus* (Fig. 1e position 13), and resolution of groups within the terminal clade (Fig. 1f) with moderate to strong bootstrap support. Membership of this clade is consistent with the inclusion of members of sect. *Gentianoides* with previous studies (Wessinger et al. 2016). However, our study shows members of *Habroanthus* as part of this terminal clade, also. The remaining taxa in *Habroanthus* group in a clade that includes members of subgenus *Penstemon* sections *Coerulei* and *Gentianoides* (Fig. 1f position 14), which is sister to a clade containing members of subgenus *Penstemon* sections *Peltanthera* and *Gentianoides*. Relative nodal support was moderate to strong within the *Peltanthera/Gentianoides* clade (Fig. 1f position 15) but was mostly lacking or moderate in the traditional clade representing subgenus *Habroanthus*. There was no taxonomic distinction to separate the red-flowered subg. *Habroanthus* sect. *Elmigera* from sect. *Glabri* (Fig. 1f position 14).

### Taxonomic Bootstrap

Three analyses were conducted to test the taxonomic circumscription of the genus: 1) subgenus, 2) section, and 3) subsection (Fig. 2). In the subgenus taxonomic bootstrap (Fig. 2a), support for the backbone is robust, and shows strong support for the major groupings recognized in *Penstemon,* except for the relationship among taxa in subgenera *Penstemon* and *Habroanthus*. The pattern of relationships among subgenera are as follows: subg. *Dasanthera* as the earliest diverging lineage and sister to a clade containing all the other subgenera. Subgenus *Cryptostemon* is sister to the core group of penstemons, and there is strong support for the grouping of taxa within subg. *Saccanthera*. It is important to note that this is a test of the current classification of major groups, and not the phylogenetic tree for the genus. However, the general order of branching for major groups matches those seen in the 43-gene phylogeny presented in Figure 1, and this is the same pattern seen for the section and subsection bootstrap analyses. For the section bootstrap analysis (Fig. 2b), the earliest diverging lineages show the same robust support as the 43-gene phylogeny, and sections within subg. *Saccanthera* are strongly supported as sister taxa. Support for taxa grouping in all recognized sections of the rest of *Penstemon* is nonexistent, but the support for subg. *Dissecti* as sister to what Wessinger et al (2016, 2019) refer to as the “crown group” of *Penstemon* has strong nodal support.

**Figure 2.**
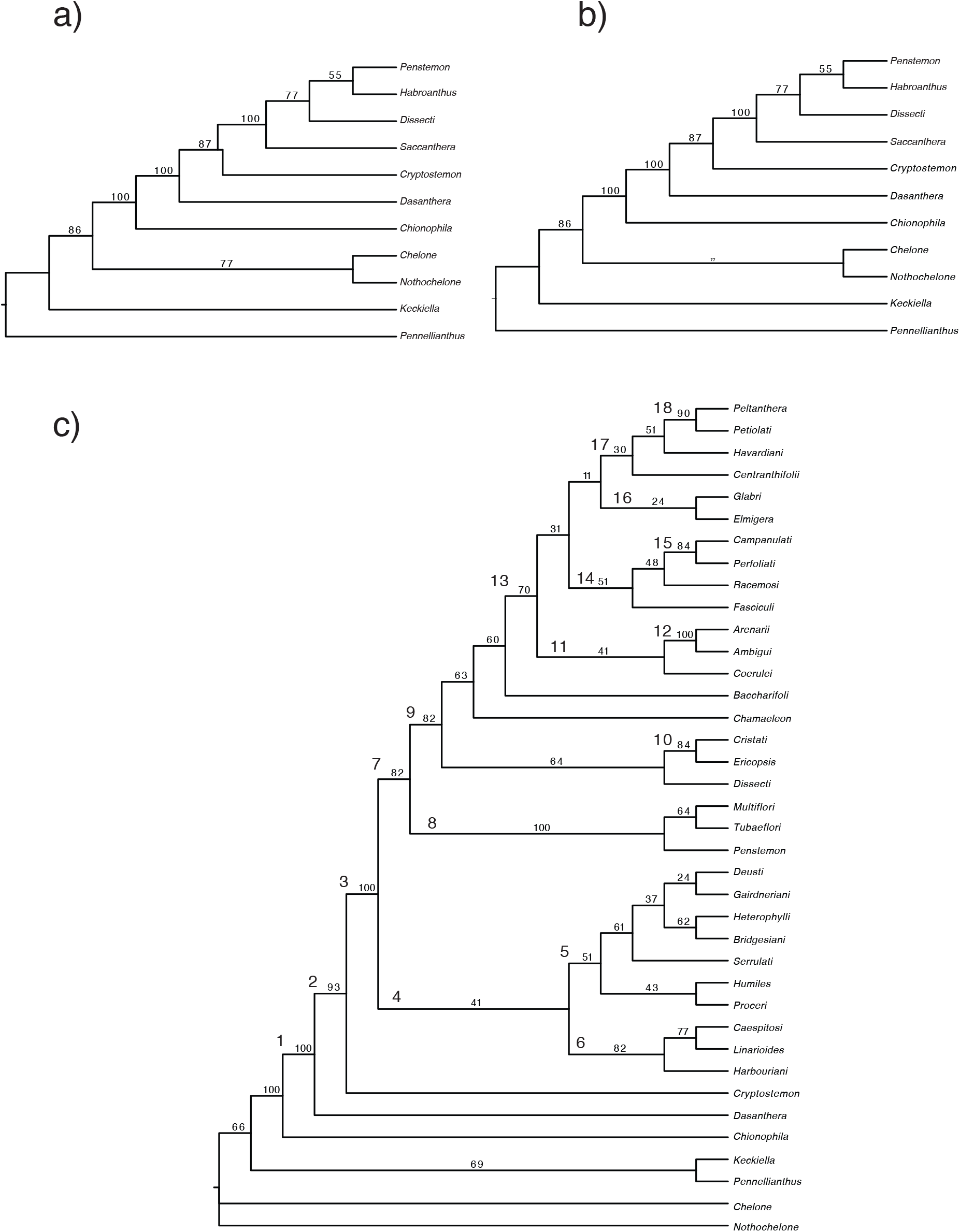
Taxonomic bootstrap results for a) subgenera, b) sections, and c) subsections.

The subsection bootstrap analysis (Fig. 2c) reveals that current classification for the genus is not well supported by phylogenetic relationships. The topology of the early diverging lineages is concordant with the overall phylogeny (Fig. 2c nodes 1, 2, and 3). In this analysis, subgenus *Saccanthera* does not hold together as a monophyletic group but is grouped with members of subg. *Penstemon* sect. *Penstemon* (Fig. 2c node 5), which is sister to three of the four subsections of sect. *Ericopsis* (Fig. 2c node 6). The topology of the latter group has strong bootstrap support. The monotypic sect. *Penstemon* subsect. *Harbouriani* is grouped with two subsections of sect. *Ericopsis* (Fig. 2c. node 6), which is consistent with the Wolfe et al. (2006) study and our 43-nuclear-gene tree (Fig. 1). Subgenus *Penstemon* sect. *Penstemon* is polyphyletic in the subsection bootstrap tree (Fig. 2c nodes 4 and 8). The relationship among subsections for the group representing members of subg. *Saccanthera*, subsects. *Deusti*, *Gairdneri*, *Humiles, Proceri, Caespitosi, Linarioides*, and *Harbouriani* to the rest of *Penstemon* has strong nodal support (Fig. 2c node 7). Three of the sect. *Penstemon* subsections are united by strong bootstrap support (Fig. 2c node 8). The position of subg. *Dissecti* is unresolved in the subsection bootstrap analysis (Fig. 2c node 9), but sect. *Ericopsis* subsect. *Ericopsis* is strongly supported as grouped with sect. *Cristati* (Fig. 2c. node 10). Taxonomic affinities for sect. *Coerulei* are uncertain (Fig. 2c node 11), but there is strong support for an affinity between sect. *Penstemon* subsect. *Arenarii* and sect. *Ambiguii* (Fig. 2c node 12). The subsection topology for the crown clade of *Penstemon* (Fig. 2c node 13) is not well resolved. Subsections of sect. *Fasciculus* are grouped with a bootstrap value of 50 (Fig. 2c node 14). However, two of the subsections group with strong support (Fig. 2c node 15). Subgenus *Habroanthus* sections *Glabri* and *Elmigera* have no bootstrap support (Fig. 2c, node 16), and relationships among what the American Penstemon Society refers to as sect. *Peltanthera*, which includes subsect. *Centranthifolii* (sect. *Gentianioides*), subsect. *Havardiani* (sect. *Gentianioides*), subsect. *Peltanthera* (sect. *Spectabiles*), and subsect. *Petiolati* (sect. *Spectabiles*) are grouped at node 17 of Figure 2c. The only group in this clade with strong support consists of subsect. *Peltanthera* and *Petiolati* (Fig. 2c node 18).

### Time Calibration and Analysis of Diversification Rates

Divergence dating with BEAST using three fossils with minimum age estimates for members of the families Bignoniaceae and Plantaginaceae produced a well-supported phylogeny for the Cheloneae, with all sampled parameters having ESS values greater than 200 and no signs of a lack of convergence in the trace plots (Fig. 3). Based on this time calibration, we infer the age of the tribe to be 10.85 my (95% HPD interval: 4.375–19.04 my), with an estimated age for the origin of *Penstemon* 3.62 mya (95% HPD interval: 1.419–6.416). Using the 95% HPD intervals for these two age estimates as secondary calibration points, plus a third calibration point for members of the Cheloneae without *Pennellianthus*, we were able to time calibrate our entire 293-taxon phylogeny with treePL. Dates inferred by treePL were slightly older for the origin of the tribe (14.94 mya) but remained roughly the same for the origin of Penstemon (3.66 mya). The divergence for the crown clade of Penstemon (excluding subg. *Dasanthera* and *P. personatus*) was placed at 2.51 mya, demonstrating that the majority of the diversification within *Penstemon* has occurred since the early Pleistocene.

**Figure 3.**
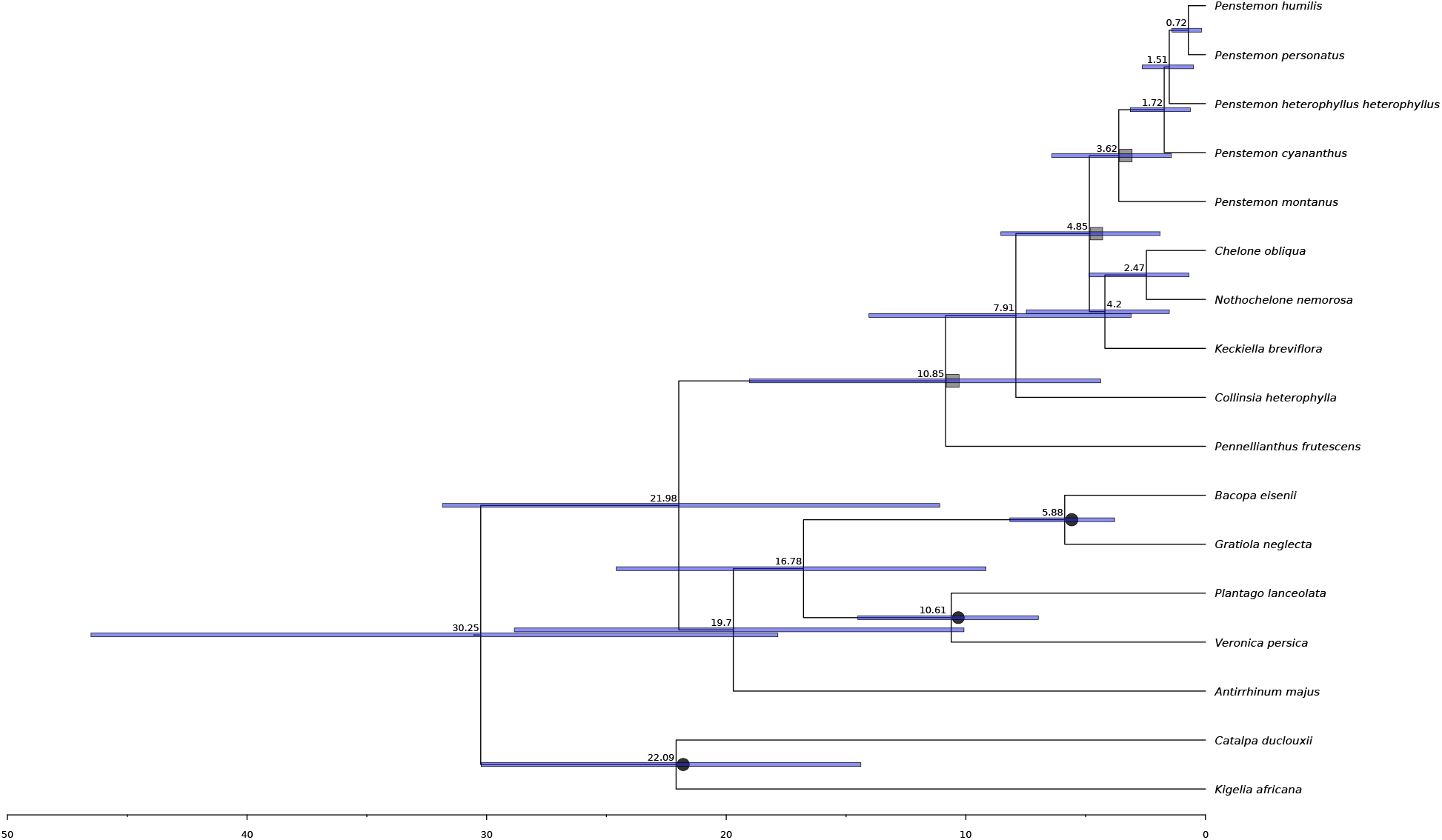
Time-calibrated tree for Chelonae and *Penstemon* based on fossils for members of Lamiales. Dark circles mark nodes that were calibrated using fossils from Vargas et al. (2014). Grey squares mark nodes that were used as secondary calibration points for dating the entire 293-taxon phylogeny with treePL.

Diversification analyses with BAMM and MEDUSA inferred similar patterns of macroevolutionary dynamics, showing a large increase in speciation rate at the base of *Penstemon* (Fig. 4). There was also some support in the posterior sample in BAMM for the placement of this shift in diversification at the base of the crown clade of *Penstemon* (Fig. S2). However, regardless of placement, both analyses support a single shift in diversification rate over more complex models with multiple shifts. While there is still a fair amount of controversy regarding the analysis of speciation and extinction rates from time-calibrated, molecular phylogenies (Moore et al. 2016; Louca and Pennell 2020), this result is consistent with the hypothesis from Wolfe et al. (2006) that *Penstemon* is a recent and rapid evolutionary radiation.

**Figure 4.**
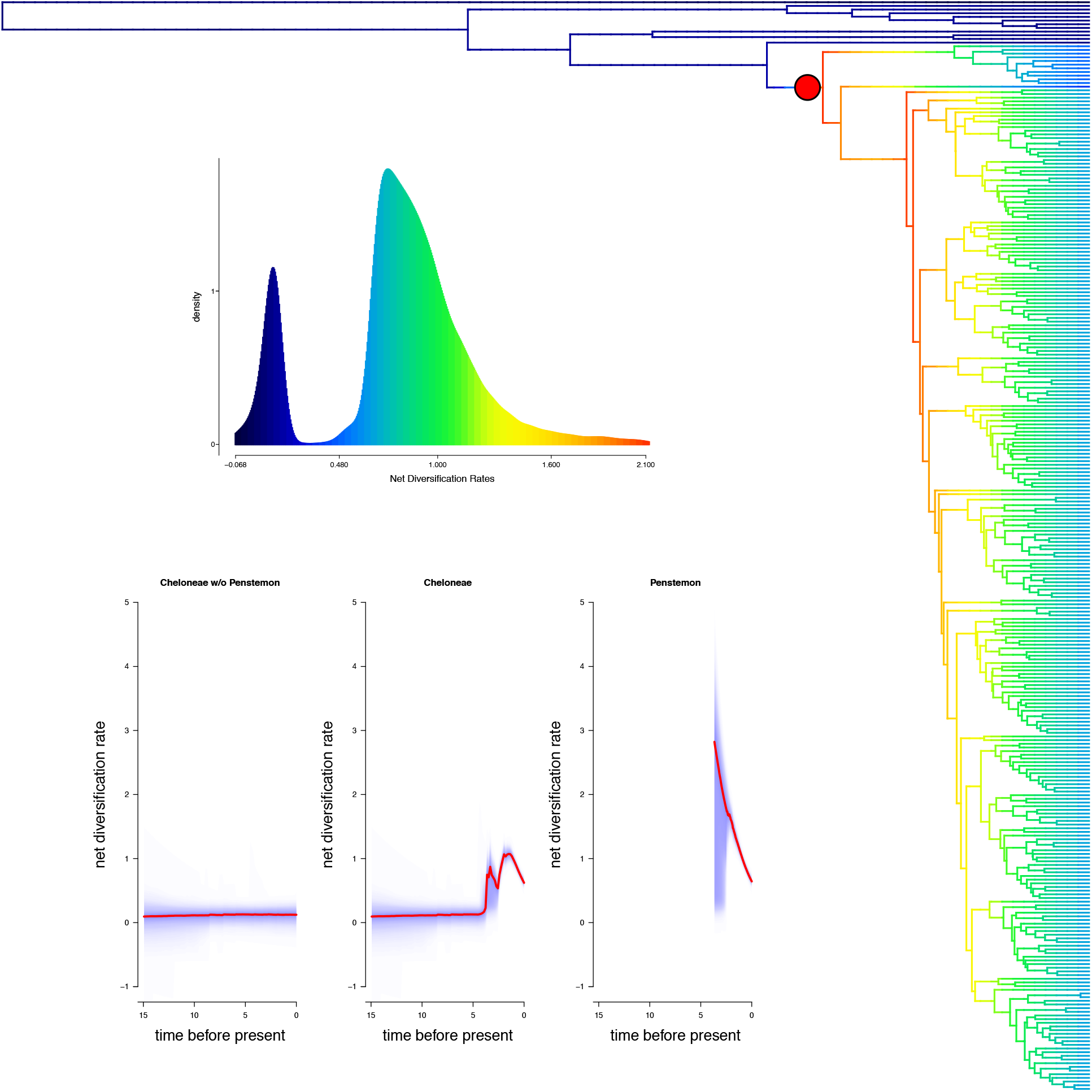
Inferred patterns of diversification using BAMM. The phylogeny shows the reconstructed rates of net diversification inferred across the tree with a single shift in diversification (best shift configuration) at the base of *Penstemon*. The top inset plot shows the distribution of net diversification rates for the entire phylogeny and the bottom inset plot shows the inferred net diversification rate through time for Chelonae without *Penstemon* (left), the Chelonae with *Penstemon* (center), and *Penstemon* alone (right).

### Biogeographic Distribution

Analyses with BioGeoBEARS consistently chose the BAYAREALIKE+J model as the best fit for all three sets of areas (Table 3; Fig. S3). The DEC+J and DIVALIKE+J models were also either the second or third best models, demonstrating that founder-event speciation events were likely to be an important process during the diversification of *Penstemon*. For ‘+J’ models, estimates of the rate of dispersal, extinction, and founder-events were all fairly consistent across sets of analyses, with extinction rates estimated to be close to 0. For analyses without the founder-event parameter, estimates of these rates varied more greatly. In particular, the BAYAREALIKE model tended to infer a higher extinction rate by almost an order of magnitude when founder-event speciation was not modeled. Despite this variation, ancestral area reconstruction for these analyses placed the most likely origin of *Penstemon* in the Eastern Cordillera (region 6), which agrees with previous hypotheses (Wolfe et al. 2002, 2006).

**Table 3.** Model comparison results for BioGeoBEARS analyses.

|  | Model | Log-Likelihood | AICc Weight | d | e | j |
| --- | --- | --- | --- | --- | --- | --- |
| Set 1 | BAYAREASLIKE+J | -372.4 | 0.59 | 0.019 | 1e-7 | 0.039 |
|  | DEC+J | -372.9 | 0.38 | 0.023 | 1e-12 | 0.036 |
|  | DIVALIKE+J | -375.6 | 0.025 | 0.026 | 1e-12 | 0.036 |
|  | DEC | -442.5 | 6.2e-31 | 0.053 | 0.038 | 0 |
|  | BAYAREALIKE | -446.9 | 7.2e-33 | 0.023 | 0.52 | 0 |
|  | DIVALIKE | -449.7 | 7.0e-34 | 0.068 | 0.024 | 0 |
| Set 2 | BAYAREALIKE+J | -424.0 | 0.92 | 0.032 | 1.0e-07 | 0.033 |
|  | DEC+J | -426.5 | 0.079 | 0.039 | 1.0e-12 | 0.029 |
|  | DIVALIKE+J | -431.8 | 0.0004 | 0.042 | 1.0e-12 | 0.029 |
|  | DEC | -479.6 | 1.9e-24 | 0.061 | 0.012 | 0 |
|  | BAYAREALIKE | -489.1 | 1.4e-28 | 0.022 | 0.54 | 0 |
|  | DIVALIKE | -492.1 | 6.9e-30 | 0.078 | 1.0e-12 | 0 |
| Set 3 | BAYAREALIKE+J | -160.3 | 1.0 | 0.0072 | 0.0028 | 0.030 |
|  | BAYAREALIKE | -250.9 | 1.4e-39 | 0.013 | 0.26 | 0 |
|  | DEC+J | -378.3 | 2.1e-95 | 0.10 | 1.0e-12 | 0.025 |
|  | DIVALIKE+J | -385.8 | 1.3e-98 | 0.11 | 1.0e-12 | 0.025 |
|  | DEC | -388.5 | 2.4e-99 | 0.11 | 1.0e-12 | 0 |
|  | DIVALIKE | -402.7 | 1.7e-105 | 0.13 | 0.016 | 0 |

One hundred sixty-five distribution map images were assembled across the phylogeny of *Penstemon* from this study. Individual images are in Figure S4, and the collective maps for the 100,000-year time slices inferred from a dated tree are in Figure S5. The time slices were inferred from the dated tree for the genus (Fig. S6). Animations of these maps in the order of appearance for either the clade position or the time slice can be seen at https://www.youtube.com/watch?v=lMLn7Gq_ZPw. Biogeographic regions (Fig. S1, Table 2) for taxa were determined by the distribution of populations shown in the individual KML files collected from the SEINet data portal (www.swbiodiversity.org). These regions were mostly based on the physiographic regions of North America (Barton et al. 2003) and were modified from Wolfe et al. (2006) to better reflect digitized collection data.

## Discussion

### Phylogenetic Inference and Time Calibration

This study is the most comprehensive phylogenetic analysis of *Penstemon* to date, with 239 of ca. 285 species included. Previous studies have established that this genus represents a large continental radiation of recent origin. Our time calibration and divergence time analyses confirm this pattern, with *Penstemon* originating around the Pliocene/Pleistocene boundary. Given the large number of species diverging in a relatively short amount of time, we estimate that *Penstemon* has one of the highest rates of diversification of any plant genus in a continental setting (Breitkopf et al. 2015; Schwery et al. 2015; Tank et al. 2015; Verboom et al. 2015; Kriebel et al. 2019). We hypothesize that this may be due to a pattern of adaptive radiation, which will be examined thoroughly in another study.

The difference in resolution from phylogenetic analyses based on ITS and cpDNA (Wolfe et al. 2006), and the current 43-nuclear-gene loci is notable, but not surprising. One would expect better tree topology resolution with more data. However, with a relatively young genus such as *Penstemon*, it is also not surprising that relative nodal support is lacking in some areas of the tree, despite the use of a large dataset. The Wessinger et al. (2019) study with the largest number of taxa (120 total species, 104 species in the crown clade) included 2306 and 2051 loci, respectively, with 72% and 68% missing data. A consensus locus was aligned “if there were at least 20 taxa per locus, no more than 50 SNPs and 8 indels per locus, and no more than 8 shared heterozygous sites across samples” (Wessinger et al. 2019). The trees in Wessinger et al. (2016, 2019) had relatively strong nodal support along the backbone of the tree, strong bootstrap support in some terminal clades, but not complete resolution of relationships throughout the tree. Comparing nuclear-gene amplicon sequences to MSG SNP data is difficult, but there are areas of agreement between the phylogenies produced by these studies as well as areas where the topologies differ. The general patterns from the phylogenies are similar enough that where backbone support is not as strong in our 43-gene tree, but in agreement with the MSG SNP results, we infer that the clade topologies represent relationships as shown in our results (Figs. 1c–1f).

Patterns of genealogical discordance and rapid rates of evolution are also apparent in our data, as evidenced by the short branches in the ASTRAL-III tree (Figs. 1, S7). Given the recent radiation of the group, such patterns are expected and are likely a result of incomplete lineage sorting caused by rapid speciation. Other processes such as hybridization and allopolyploidy have also been documented in *Penstemon* (Keck 1945; Wolfe et al. 1998a,b; Broderick et al. 2011) and further complicate the accurate inference of phylogeny where they occur. However, the coalescent branch lengths in the ASTRAL-III tree (Figs. 1, S7) indicate that high levels of discordance are more prevalent towards the tips of the tree, with little evidence of discordance among the major lineages of the Cheloneae. In general, it is important to consider the effect of these sources of discordance on downstream inferences. For the analyses we conducted here (divergence dating, diversification, and biogeography), we are primarily focused on nodes deeper in the phylogeny that are less affected by the incongruence seen in other parts of our tree. Because of this, we argue that our results regarding the timing and diversification of the genus are robust, especially considering the agreement between our work and previous studies, as well as our dense sampling and the comparatively low amount of missing data in our analyses.

Our inferred timing for the biogeographic and diversification history (Table S2) of *Penstemon* in the Pliocene/Pleistocene supports previous hypotheses regarding its spread across North America during these epochs’ dynamic periods of glaciation (Wolfe et al. 2006, Ehlers and Gibbard 2007; Ehlers et al. 2018). And while the age we infer for *Penstemon* places its origin slightly farther back in time than previously thought (3.66 Mya versus ~2.5 Mya; Wolfe et al. 2006), the timing of divergence for the majority of species in the crown clade still coincides with the onset of the Pleistocene. Absent more closely related fossils than those from Vargas et al. (2014) at the family level, our divergence dating with secondary calibration points represents the only fossil-based divergence estimation conducted for the tribe. This time-calibrated phylogeny was crucial for understanding the pattern and timing of diversification, showing that *Penstemon* not only diversified at a much higher rate than other members of Cheloneae (Fig. 4) but that the shift in diversification rate occurred at the base of the clade, coinciding with its inferred origin in the Eastern Cordillera and subsequent dispersal through founder events across the continent. Compared with other groups of angiosperms, the net diversification rate for *Penstemon* is consistent with estimates for the Lamiales (Magallón and Sanderson 2002; Magallón and Castillo 2009); however, *Penstemon* is noteworthy as a particularly exceptional radiation when considering that its diversity has arisen in only the last 3.66 My, with the bulk of diversification occurring within the past 1.8 My. And while criticisms of diversification analyses are continuing to raise important considerations for the interpretability of absolute estimates of diversification dynamics (Moore et al. 2016; Louca and Pennell 2020), especially with regard to extinction rates, the relative pattern of rapid diversification in *Penstemon* compared to its sister lineages remains clear.

### Taxonomic Implications

Taxonomic bootstrapping of the subgenera, sections, and subsections of *Penstemon* reveals some interesting patterns, which speak to the need for taxonomic revision of the genus. In the subgenus taxonomic bootstrap (Fig. 2a), the traditional categories for subg. *Dasanthera*, *Cryptostemon*, *Saccanthera*, and *Dissecti* are not in conflict. The low bootstrap value for the node representing subgenera *Penstemon* and *Habroanthus* indicates some taxonomic conflict with placement of taxa in these categories. The taxonomic bootstrap of sections (Fig. 2b) shows the same early branching pattern support for *Dasanthera*, *Cryptostemon*, and the grouping of sections within subg. *Saccanthera*. Indications of taxonomic conflict occur throughout the remainder of the tree as evidenced by low to moderate bootstrap values for all nodes beyond subg. *Saccanthera*.

The subsection bootstrap analysis (Fig. 2c) yields insights into current taxonomic structure of the genus. Nodes 1–3 show the same strong support for the early diverging lineages (subg. *Dasanthera* and subg. *Cryptostemon* relationship and their placement as sister to the rest of the genus). Node 4 reveals conflict in taxonomic placement of some taxa within subgenera *Saccanthera* and *Penstemon*, particularly with the inclusions of subg. *Penstemon* sect. *Penstemon* subsections *Deusti* and *Gairdneri* with sections and subsections of subg. *Saccanthera*. Low bootstrap values for the node representing sect. *Penstemon* subsections *Humiles* and *Proceri* also indicate taxonomic problems within those groups. Node 6 indicates that two of the subsections of section *Ericopsis* are well defined, but these also group with the monotypic sect. *Penstemon* subsect. *Harbouriani*. Three of the subsections of section *Penstemon* have high nodal support (Fig. 2c, Node 8). Other indicators of taxonomic conflict are represented by nodes 10 (placement of sect. *Ericopsis* subsect. *Ericopsis* as sister to sect. *Cristati*), 12 (sect. *Penstemon* subsect. *Arenarii* as sister to sect. *Ambiguii*), 14 (low nodal support for taxa within section *Fasciculus*), 16 (no support for subg. *Habroanthus* sections *Glabri* and *Elmigera*), and 17 (placement of taxa within section *Gentianioides* with no nodal support).

In the context of the phylogeny presented here (Fig. 1), clearly there is a need to revise the circumscription of the genus *Penstemon.* Similar patterns of taxa grouping outside their assigned sections and subsections have been seen in other recent studies with subsets of species (Wessinger et al. 2016, 2019). Based on the current study, together with the Wessinger et al studies (2016, 2019), the number of subgenera should be decreased from six to four: *Dasanthera*, *Cryptostemon*, *Saccanthera*, and *Penstemon*. The traditional subgenus *Habroanthus* should be designated as a section, sans designated subsections *Glabri* and *Elmigera*. The traditional monotypic subgenus *Dissecti*, should be section *Dissecti*, given its phylogenetic placement (Fig. 1c). Many taxa need to be shuffled from their traditional assigned subgenera, subsections, or sections to other groupings. Recommended changes based on phylogenetic studies can be found in Table S3.

### Biogeography

Biogeographic analysis of *Penstemon* with BioGeoBEARS (Fig. S3) confirmed previous hypotheses for the origin of the genus in the eastern Cordillera of North America and its subsequent spread across the continent (Straw 1966; Wolfe et al. 2002). The importance of founder-event speciation for this process is demonstrated here for the first time, indicating that dispersal to new, ancestrally unoccupied areas was a key mechanism for the evolution and expansion of *Penstemon*. Given the dynamic nature of glaciation in North America during the diversification of the genus (see more details below), this pattern makes sense and is likely connected with the recurrent formation of “sky islands” in western and southwestern North America (Rehfeldt 1999; Knowles 2001; Hewitt 2004). As with our analyses of diversification with BAMM and MEDUSA, we find that the inferred rate of extinction is effectively zero for the most well-supported models. However, we do note that the inference of extinction rates from molecular phylogenies can be problematic (Rabosky 2010; Louca and Pennell 2020), so we interpret this absence of extinction with caution, especially considering the paucity of fossil evidence to corroborate diversification patterns for the Cheloneae. Nevertheless, the consistent pattern of elevated net diversification in *Penstemon* and its probable connection with the importance of founder-event speciation as inferred by BioGeoBEARS helps point to a possible mechanism for its rapid radiation.

The mapped biogeographic patterns for *Penstemon* are associated with the glaciation cycles of the Pleistocene (Table S2, Figs. S4, S5; https://www.youtube.com/watch?v=lMLn7Gq_ZPw). There were 13 major glaciations and nine minor glaciations during the 2.7 years of the Pleistocene (Ehlers et al. 2018). Although most biogeographic histories emphasize the effect of the last glacial maximum, we argue that each cycle of glaciation had an impact on the diversification of *Penstemon*. The first four major glaciations occurred between 2.7 mya and 2.0 mya. During that time frame the earliest diverging lineage, subgenus *Dasanthera*, spread through the northeastern cordillera into the Cascade-Sierra cordillera and Pacific Northwest. The next major glaciation occurred between 2.0–1.8 mya. During this timeframe, *P. dissectus*, endemic to Georgia, appears in the eastern coastal plain. Given the phylogenetic position of *P. dissectus* in the middle of the phylogenetic tree (Figs. 1c, S2), one can confidently infer that distribution of species was more widespread throughout the southern reaches of the North American continent, but that extinction events occurred during the first million years of the Pleistocene. This is supported in our BAMM analyses (Fig. 4) where diversification shows an early burst along the backbone of the phylogenetic tree, followed by a decrease and/or extinction as the genus evolved. Figure 4 also illustrates two periods of diversification for tribe Cheloneae including *Penstemon*. There was an initial burst followed by a dramatic decline in net diversification rate, and a second burst and decline corresponding with the phylogenetic time frame represented in Table S2.

Between 1.8 mya and 1.2 mya there was one major and four minor glaciation episodes. During this timeframe, most of the diversification was taking place in the eastern cordillera and intermountain region (Table S2), except for the appearance of *P. smallii* in the Appalachian region. Extension of *Penstemon* into the southwest region, southern Great Plains, and highlands of Mexico had just begun. From 1.2 to 1.1 mya two major glaciation cycles took place. The diversification into regions shows a north-south oscillation from the Cascade-Sierra and eastern cordillera ranges as far north as Alaska, and into the southwest and Mexico highlands. The southern Great Plains in the west also showed activity as did the southern Intermountain Region. From 1.1 mya to 1.0 mya there was one minor glaciation. There appeared to be a pattern of diversification from the eastern Pacific Northwest, southeast into the eastern cordillera and western Great Plains, and east into the Intermountain region.

Diversification of *Penstemon* was most active between 1.0 mya to 0.5 mya (Table S2, Figs. S4, S5). During this time frame there were two major glaciations (0.9–0.8 mya and 0.7–0.6 mya) and one minor glaciation (1.0–0.9 mya). During the latter time period, there was an expansion from the southern Cascade-Sierra and intermountain regions north and east into the eastern cordillera and northern Cascade-Sierra. In the east there was an expansion from the Appalachian region into the southern interior lowlands and eastern Great Plains, and expansion through the coastal plain. From the southeastern cordillera there was an expansion into the Intermountain Region, then southwest and back into the Eastern Cordillera. During the first major glaciation cycle of this time period, there was an expansion south from the Cascade-Sierra to the Southwest Cordillera and Baja California. This was concurrent with a trajectory from the Intermountain region into the Southwest and Southwest Cordillera. During the subsequent interglacial period, the diversification followed a track northward into the Eastern Cordillera and Intermountain Region, and south into the Mexico Highlands. There was also an expansion from the Eastern Cordillera into the Great Plains, and in the east, there was an expansion through the eastern interior lowlands and northward in the Appalachian region. During the second major glaciation of this period, there was another cycle of diversification to the south (Intermountain region to Southwest, and Mexico Highlands), and diversfication into the southern Great Plains. The next glacier-free 100,000 years had activity in the Pacific Northwest and Cascade-Sierra, and an eastward movement of diversification in the Great Plains. There was also diversification in the Interior Lowlands.

The diversification of *Penstemon* slowed down during the most recent 500,000 years, with four major glaciations and two minor glaciation cycles, including the Last Glacial Maximum period. During this time period, most of the action was in the southern areas of the North America continent, notably in the Intermountain Region, Southwest, Southwest Cordillera and Baja California, and the Mexico Highlands.

## Conclusions

*Penstemon* is a remarkable genus in its rapid diversification into most ecological regions found in North America (Fig. S1). The diversification, phylogeny, and biogeographic history suggest a rapid evolutionary radiation throughout North America during the Pleistocene. The pulses of diversification associated with major and minor glaciation periods, in the context of the models supported by our BioGeoBEARS analyses, are consistent with the colonization of newly available ecological niches during interglacial cycles. Considering the immense diversity in corolla, anther, staminode, leaf, inflorescence, and habit morphology, taken together with the large amount of variation in habitats, it is likely that *Penstemon* has undergone an adaptive radiation in a continental setting.

## Acknowledgements

The authors thank members of the Wolfe and Kubatko labs for comments that helped to improve this manuscript. We also thank N. Holmgren, M. Stevens, G. Moffit, past and current members of the Wolfe Lab, and members of the American Penstemon Society for help with collecting specimens. In addition, we thank the Wolfe Lab graduate and undergraduate students, past and current, for their assistance with DNA extractions, database management, and sequencing preparation. Without their dedicated contributions this study would never have been possible.

## Data Availability

Raw sequencing reads, aligned DNA sequences, data matrices, tree files, and code for all analyses is available on Dryad (doi:XXXX).

