## Appendix 1 for "Phylogenetics of a Rapid, Continental Radiation: Diversification, Biogeography, and Circumscription of the Beardtongues (*Penstemon*; Plantaginaceae)"

**Appendix 1**. Taxon list for DNA sampling.

| Taxon | DNA Accession Number | Collector Accession/Herbarium |
|---|---|---|
| *Chelone obliqua* | 96-0073 | A. D. Wolfe 586/OS |
| *Chionophila tweedyi* | 99-1511 | S. L. Datwyler 110/OS |
| *Keckiella antirrhinoides* | 00-0135 | Paul Wilson /M.S. Vanenzuela Oak Grove 4Jun.98/SVF |
| *Keckiella cordifolia* | 97-0181 | Paul Wilson 3513/SVF |
| *Keckiella corymbosa* | 00-0133 | Paul Wilson 3549/SVF |
| *Keckiella lemonii* | 16-0101 | A. D. Wolfe 1518/OS |
| *Keckiella rothrockii* var. *jacintensis* | 97-0180 | Paul Wilson 3512/SVF |
| *Nothochelone numerosa* | 00-0129 | Paul Wilson 3552/SVF |
| *Pennellianthus frutescens* | 98-0549 | Wolfe s.n./OS |
| *Penstemon abietienus* | 11-0385 | A. Wenzel 08/OS |
| *Penstemon acaulis* var. *acaulis* | 12-0192 | Jouse 19/OS |
| *Penstemon acaulis* var. *yampaensis* | 12-0184 | Jouse 11/OS |
| *Penstemon acuminatus var. acuminatus* | 96-0186 | S. L. Datwyler 03/OS |
| *Penstemon alamosensis* | 99-1391 | A. D. Wolfe 813/OS |
| *Penstemon albertinus* | 15-0469 | A. Wenzel 41/OS |
| *Penstemon albomarginatus* | 10-0101 | A. D. Wolfe 1358/OS |
| *Penstemon ambiguus* | 14-0146 | A. D. Wolfe 1433/OS |
| *Penstemon ammophilus* | 15-0305 | A. D. Wolfe 1464/OS |
| *Penstemon amphorellae* | 14-0111 | J. Gutiérrez G. 392/IEB |
| *Penstemon angelicus* | 15-0192 | Rebman/ SD 228971 |
| *Penstemon anguineus* | 16-0064 | A. D. Wolfe 1505/OS |
| *Penstemon angustifolius* var. *angustifolius* | 99-1741 | Mt. West Enviro Services 7948/OS |
| *Penstemon angustifolius* var. *caudatus* | 99-1653 | Mt. West Enviro. Services 7947/OS |
| *Penstemon angustifolus* var. *dulcis* | 15-0282 | A. D. Wolfe 1457/OS |
| *Penstemon angustifolus* var. *venosus* | 15-0334 | A. D. Wolfe 1475/OS |
| *Penstemon angustifolus* var. *vernalensis* | 15-0250 | A. D. Wolfe 1450/OS |
| *Penstemon arenicola* | 99-1654 | Mt. West Enviro. Services 7957/OS |
| *Penstemon aridus* | 99-1492 | A. Lutz Billy Creek/OS |
| *Penstemon attenuatus* var. *attenuatus* | 12-0347 | P. D. Blischak 19/OS |
| *Penstemon attenuatus* var. *militaris* | 12-0275 | P. D. Blischak 12/OS |
| *Penstemon attenuatus* var. *palustris* | 15-0690 | P. D. Blischak 61/OS |
| *Penstemon attenuatus* var. *pseudoprocerus* | 15-0485 | P. D. Blischak 42/OS |
| *Penstemon atwoodii* | 15-0211 | A. D. Wolfe 1439/OS |
| *Penstemon auriberbis* | 99-1485 | A. D. Wolfe 843/OS |
| *Penstemon azureus* var. *azureus* | 17-0004 | A. D. Wolfe 59/OS |
| *Penstemon baccharifolius* | 03-0167 | Paul Wilson 3938/SVF |

| | | |
|---|---|---|
| *Penstemon barbatus* | 98-0495 | A. D. Wolfe 778/OS |
| *Penstemon barnebyi* | 15-0355 | A. D. Wolfe 1494/OS |
| *Penstemon barrettiae* | 96-0047 | A. D. Wolfe 605/OS |
| *Penstemon bicolor* var. *bicolor* | 99-0368 | A. D. Wolfe 795/OS |
| *Penstemon bicolor* var. *roseus* | 99-1697 | A. D. Wolfe 805/OS |
| *Penstemon bolianus* | 14-0083 | A. G. Zacaría-Correa 87/IEB |
| *Penstemon bracteatus* | 15-0405 | A. D. Wolfe 1489/OS |
| *Penstemon breviculus* | 15-0349 | A. D. Wolfe 1479/OS |
| *Penstemon buckleyi* | 16-0371 | R. Rodriguez 29/OS |
| *Penstemon caesius* | 98-0056 | Paul Wilson 3482/SVF |
| *Penstemon caespitosus* var. *caespitosus* | 11-0196 | A. J. Wenzel 01/OS |
| *Penstemon caespitosus* var. *desertipicti* | 11-0557 | A. J. Wenzel 15/OS |
| *Penstemon caespitosus* var. *perbrevis* | 11-0307 | A. J. Wenzel 05/OS |
| *Penstemon calcareus* | 15-0194 | Howe/ SD 113661 |
| *Penstemon californicus* | 12-0074 | A. J. Wenzel 42/OS |
| *Penstemon calycosus* | 15-0426 | P. D. Blischak 69/OS |
| *Penstemon campanulatus* var. *campanulatus* | 14-0095 | A. G. Zacarías-Correa 83/IEB |
| *Penstemon campanulatus* var. *chihuahuensis* | 00-0121 | Paul Wilson 3595/SVF |
| *Penstemon canescens* | 99-1468 | A. D. Wolfe 854/OS |
| *Penstemon cardinalis* var. *cardinalis* | 03-0169 | Paul Wilson 3957/SVF |
| *Penstemon cardwellii* | 96-0044 | A. D. Wolfe 602/OS |
| *Penstemon carnosus* | 98-0508 | ADW 757/OS |
| *Penstemon caryi* | 99-0298 | A. W. Lutz C/OS |
| *Penstemon centranthifolius* | 27 | A. D. Wolfe 209/OS |
| *Penstemon cerrosensis* | 15-0195 | Moran/ SD 53963 |
| *Penstemon cinicola* | 12-0577 | P. D. Blischak 32/OS |
| *Penstemon clevelandii* var. *clevelandii* | 98 | A. D. Wolfe 186/OS |
| *Penstemon clevelandii* var. *connatus* | 18 | A. D. Wolfe 313/OS |
| *Penstemon clevelandii* var. *mojavensis* | 00-0137 | Wilson s.n./OS |
| *Penstemon clutei* | 99-1388 | A. D. Wolfe 810/OS |
| *Penstemon cobaea* | 99-1452 | A. D. Wolfe 838/OS |
| *Penstemon commarrhenus* | 01-0177 | Mt. West Enviro Services 8780/OS |
| *Penstemon compactus* | 15-0694 | Mancuso 235/ORED |
| *Penstemon concinnus* | 16-0131 | M. Stevens 208/BRY-V |
| *Penstemon confertus* | 12-0327 | P. D. Blischak 18/OS |
| *Penstemon confusus* | 14-0148 | A. D. Wolfe 1435/OS |
| *Penstemon coriaceus* | 14-0001 | A. G. Zacaría-Correa 102/IEB |
| *Penstemon crandallii* var. *atratus* | 11-0667 | A. J. Wenzel 19/OS |
| *Penstemon crandallii* var. *crandallii* | 11-0697 | A. J. Wenzel 20/OS |

| | | |
|---|---|---|
| *Penstemon cusickii* | 16-0132 | J. F. Smith/BRV-Y |
| *Penstemon cyananthus* var. *cyananthus* | 98-0514 | A. D. Wolfe 672/OS |
| *Penstemon cyananthus* var. *subglaber* | 16-0465 | A. D. Wolfe 1540/OS |
| *Penstemon cyanocaulis* | 98-0091 | F. R. Stermitz s.n./OS |
| *Penstemon cyathophorus* | 99-1160 | Mt. West Enviro Services 7982/OS |
| *Penstemon dasyphyllus* | 01-0104 | A. D. Wolfe 918/OS |
| *Penstemon davidsonii* var. *davidsonii* | 96-0226 | A. D. Wolfe 614/OS |
| *Penstemon davidsonii* var. *praeteritus* | 96-0021 | A. D. Wolfe 662/OS |
| *Penstemon deaveri* | 16-0133 | G. L. Pyrah 166/BRY-V |
| *Penstemon debilis* | 97-0189 | McMullen s.n./OS |
| *Penstemon degeneri* | 113 | C. English LPN 113/University of Denver |
| *Penstemon deustus* var. *deustus* | 98-0498 | A. D. Wolfe 781/OS |
| *Penstemon deustus* var. *suffrutescens* | 00-0141 | Paul Wilson 3551/SVF |
| *Penstemon digitalis* | 99-0096 | C. P. Randle 72/OS |
| *Penstemon diphyllus* | 010-0102 | A. D. Wolfe 919/OS |
| *Penstemon discolor* | 12-0167 | Reichenbacher 50/OS |
| *Penstemon dissectus* | 98-0542 | Leege s.n./OS |
| *Penstemon duchesnensis* | 11-0951 | A. J. Wenzel 33/OS |
| *Penstemon eatonii* var. *eatonii* | 10-0175 | Mt. West Enviro Services 8760/OS |
| *Penstemon eatonii* var. *exsertus* | 16-0127 | A. D. Wolfe 1526/OS |
| *Penstemon eatonii* var. *undosus* | 15-0313 | A. D. Wolfe 1567/OS |
| *Penstemon elegantulus* | 15-0698 | Gray 4231/ORED |
| *Penstemon ellipiticus* | 96-0208 | J. Walker 259 |
| *Penstemon eriantherus* var. *cleburnei* | 99-1755 | Mt. West Enviro Services 7958/OS |
| *Penstemon eriantherus* var. *eriantherus* | 99-0016 | S. L. Datyler 47/OS |
| *Penstemon euglaucus* | 12-0562 | P. D. Blischak 33/OS |
| *Penstemon fasciculatus* | 14-0094 | MCD 21/IEB |
| *Penstemon fendleri* | 99-1450 | A. D. Wolfe 836/OS |
| *Penstemon filiformis* | 16-0067 | A. D. Wolfe 1506/OS |
| *Penstemon flavescens* | 15-0737 | P. D. Blischak 63/OS |
| *Penstemon floridus* | 11-0355 | G. Moore s.n./OS |
| *Penstemon flowersii* | 15-0236 | A. D. Wolfe 1446/OS |
| *Penstemon franklinii* | 15-0411 | A. D. Wolfe 1491/OS |
| *Penstemon fruticiformis* | 99-1397 | A. D. Wolfe 819/OS |
| *Penstemon fruticosus* var. *fruticosus* | 99-0169 | S. L. Datwyler 50/OS |
| *Penstemon fruticosus* var. *serratus* | 02-0470 | Mt. West Enviro Services 9285/OS |
| *Penstemon gairdneri* var. *gairdneri* | 12-0326 | P. D. Blischak 17/OS |
| *Penstemon gairdneri* var. *oreganus* | 96-0003 | A. D. Wolfe 660/OS |
| *Penstemon gentianoides* | 14-0109 | Saimain 2014-022/IEB |
| *Penstemon gibbensii* | 99-1657 | Mt. West Enviro Services 7973/OS |
| *Penstemon glaber* var. *alpinus* | 99-1761 | Mt. West Enviro Services 8031/OS |
| *Penstemon glaber* var. *brandegeei* | 99-1459 | A. D. Wolfe 845/OS |
| *Penstemon glabrescens glabrescens* | 11-0770 | A. J. Wenzel 25/OS |
| *Penstemon glabrescens* var. *taosensis* | 12-0133 | A. J. Wenzel 47/OS |
| *Penstemon glandulosus* | 96-0195 | A. D. Wolfe 639/OS |
| *Penstemon globosus* | 16-0584 | A. D. Wolfe 1566/OS |
| *Penstemon goodrichii* | 15-0231 | A. D. Wolfe 1445/OS |

| | | |
|---|---|---|
| *Penstemon gormanii* | 99-1364 | Armbruster s.n./ALA |
| *Penstemon gracilentus* | 16-0036 | A. D. Wolfe 1500/OS |
| *Penstemon gracilis* | 99-1409 | A. D. Wolfe 830/OS |
| *Penstemon grahamii* | 15-0241 | A. D. Wolfe 1447/OS |
| *Penstemon grandiflorus* | 99-1762 | Mt. West Enviro Services 8065/OS |
| *Penstemon griffinii* | 99-0173 | Pate and Porter 10299/RM |
| *Penstemon grinnellii* var. *grinnellii* | 62 | A. D. Wolfe 303/OS |
| *Penstemon grinnellii* var. *scrophularioides* | 59 | A. D. 464/OS |
| *Penstemon guadalupensis* | 16-0168 | Webster and Westlund 34072/TEX |
| *Penstemon hallii* | 02-0473 | Mt. West Enviro Services 9502/OS |
| *Penstemon harbourii* | 98-0529 | J. Thomson s.n./OS |
| *Penstemon harringtonii* | 98-0482 | A. D. Wolfe 784/OS |
| *Penstemon hartwegii* | 14-0119 | A. G. Zacaría-Correa 108/IEB |
| *Penstemon harvardii* | 99-1463 | A. D. Wolfe 849/OS |
| *Penstemon haydenii* | 17-0013 | Stubbendeck s.n./OS |
| *Penstemon hesperius* | 16-0128 | G. Maffit s.n./OS |
| *Penstemon heterodoxus* var. *heterodoxus* | 16-0014 | A. D. Wolfe 1498/OS |
| *Penstemon heterophyllus* | 01-0116 | L. Malessa 13/SVF |
| *Penstemon heterophyllus* var. *purdyi* | 17-0003 | A. D. Wolfe 574/OS |
| *Penstemon hidalgensis* | 16-0170 | Zamudio 13587/IEB |
| *Penstemon hirsutus* | 99-1488 | Wolfe s.n./OS |
| *Penstemon humilis* var. *brevifolius* | 98-0507 | A. D. Wolfe 573/OS |
| *Penstemon humilis* var. *humilis* | 98-0513 | A. D. Wolfe 761/OS |
| *Penstemon humilis* var. *obtusifolius* | 15-0202 | A. D. Wolfe 1430/OS |
| *Penstemon idahoensis* | 15-0699 | Atwood 13646/ORED |
| *Penstemon imberbis* | 14-0115 | A. G. Zacaría-Correa 107/IEB |
| *Penstemon immanifestus* | 15-0421 | A. D. Wolfe 1493.OS |
| *Penstemon incertus* | 17-0010 | A. D. Wolfe 465/OS |
| *Penstemon inflatus* | 99-1389 | A. D. Wolfe 811/OS |
| *Penstemon isophyllus* | 14-0125 | A. G. Zacaría-Correa 125/IEB |
| *Penstemon jamesii* | 99-1385 | A. D. Wolfe 807/OS |
| *Penstemon janishiae* | 15-0701 | A. Tiehm 12949/BRY:V |
| *Penstemon kingii* | 16-0135 | S. Goodrich 11236/BRY:V |
| *Penstemon labrosus* | 98-0073 | Hogue 87.7/OS |
| *Penstemon laetus* var. *laetus* | 96-0212 | A. D. Wolfe 665/OS |
| *Penstemon laetus* var. *sagittatus* | 00-0134 | Paul Wilson 3550/SVF |
| *Penstemon laevis* | 14-0142 | A. D. Wolfe 1429/OS |
| *Penstemon lanceolatus* | 14-0007 | A. G. Zacaría-Correa 96/IEB |
| *Penstemon laricifolius* var. *exifolius* | 12-0175 | A. D. Wolfe 1385/OS |
| *Penstemon laricifolius* var. *laricifolius* | 12-0171 | A. D. Wolfe 1383/OS |
| *Penstemon lavendulus* | 97-0031 | F. R. Stermitz s.n./OS |
| *Penstemon laxiflorus* | 97-0387 | Elisens s.n./OS |
| *Penstemon laxus* | 15-0702 | Smith 8123/ORED |
| *Penstemon leiophyllus* var. *leiophyllus* | 01-0178 | Mt. West Enviro Services 8795/OS |
| *Penstemon lentus* var. *albiflorus* | 15-0336 | A. D. Wolfe 1477/OS |
| *Penstemon lentus* var. *lentus* | 15-0368 | A. D. Wolfe 1483/OS |
| *Penstemon leonardii* var. *leonardii* | 16-0440 | A. D. Wolfe 1569/OS |
| *Penstemon leonensis* | 14-0089 | A. G. Zacaría-Correa 94/IEB |

| | | |
|---|---|---|
| *Penstemon leptanthus* | 98-0051 | A. D. Wolfe 758/OS |
| *Penstemon linarioides* var. *coloradoensis* | 14-0688 | A. J. Wenzel 49/OS |
| *Penstemon linarioides* var. *compactifolius* | 15-0056 | A. J. Wenzel 50/OS |
| *Penstemon linarioides* var. *linarioides* | 12-0114 | A. J. Wenzel 45/OS |
| *Penstemon linarioides* var. *sileri* | 11-0964 | A. J. Wenzel 35/OS |
| *Penstemon longiflorus* | 01-0176 | Mt. West Enviro Services 8766/OS |
| *Penstemon luculentus* | 15-0256 | A. D. Wolfe 1451/OS |
| *Penstemon lyallii* | 96-0095 | S. L. Datwyler 42/OS |
| *Penstemon mensarum* | 96-0134 | B. J. Thomson 96-17/OS |
| *Penstemon miniatus* | 14-0114 | MF 85/IEB |
| *Penstemon miser* | 15-0703 | Holmgren 8831/ORED |
| *Penstemon moffatii* | 15-0224 | A. D. Wolfe 1443/OS |
| *Penstemon monoensis* | 15-0197 | Everett/SD 48552 |
| *Penstemon montanus* | 96-0178 | S. L. Datwyler 51/OS |
| *Penstemon moriahensis* | 16-136 | Swenson 26/BRY-V |
| *Penstemon mucronatus* | 99-1658 | Mt. West Enviro Services 7974/OS |
| *Penstemon multiflorus* | 99-0141 | A. D. Wolfe 831/OS |
| *Penstemon nanus* | 15-0416 | A. D. Wolfe 1492/OS |
| *Penstemon neomexicanus* | 99-1407 | A. D. Wolfe 828/OS |
| *Penstemon neotericus* | 16-0105 | A. D. Wolfe 1519/OS |
| *Penstemon newberryi* var. *berryi* | 16-0046 | A. D. Wolfe 1502/OS |
| *Penstemon nitidus* var. *nitidus* | 99-1652 | Mt. West Enviro Services 7935/OS |
| *Penstemon nudiflorus* | 101 | A. D. Wolfe 916/OS |
| *Penstemon occiduus* | 16-0167 | Wehbe 216/MEX |
| *Penstemon oklahomensis* | 97-0386 | Elisens s.n./OS |
| *Penstemon ophianthus* | 99-1449 | A. D. Wolfe 835/OS |
| *Penstemon ovatus* | 17-0006 | A. D. Wolfe 608/OS |
| *Penstemon pachyphyllus* var. *congestus* | 15-0314 | A. D. Wolfe 1468/OS |
| *Penstemon pachyphyllus* var. *pachyphyllus* | 15-0214 | A. D. Wolfe 1441/OS |
| *Penstemon pahutensis* | 16-0165 | Bostick/TEX 5078 |
| *Penstemon pallidus* | 16-0188 | A. D. Wolfe 1534/OS |
| *Penstemon palmeri* | 136 | A. D. Wolfe 294/OS |
| *Penstemon palmeri* var. *eglandulosus* | 01-0174 | Mt. West Enviro Services 8759/OS |
| *Penstemon pappilatus* | 01-0095 | Kimball 121/SVF |
| *Penstemon parryi* | 15-0298 | A. D. Wolfe 1462/OS |
| *Penstemon parvulus* | 96-0048 | A. D. Wolfe 584/OS |
| *Penstemon parvus* | 16-0154 | R. L. Johnson & M. R. Stevens/BRY-V 622304 |
| *Penstemon patens* | 98-0525 | G Aldridge & H. Crandall s.n./OS |
| *Penstemon patricus* | 16-0155 | S. G. Harris 3360/BYU 462856 |
| *Penstemon payettensis* | 96-0198 | A. D. Wolfe 659/OS |
| *Penstemon peckii* | 12-0540 | P. D. Blischack 31/OS |
| *Penstemon penlandii* | 16-0156 | N. D. Atwood 33630/BRY-V 613952 |
| *Penstemon personatus* | 16-0116 | A. D. Wolfe 1522/OS |
| *Penstemon petiolatus* | 14-0147 | A. D. Wolfe 1434/OS |
| *Penstemon pinifolius* | 12-0130 | A. J. Wenzel 46/OS |
| *Penstemon pinorum* | 14-0149 | A. D. Wolfe 1436/OS |
| *Penstemon plagapineus* | 14-0097 | A. G. Zacaría-Correa 85/IEB |

| | | |
|---|---|---|
| *Penstemon platyphyllus* | 16-0589 | A. D. Wolfe 1567/OS |
| *Penstemon potosinus* | 14-0005 | A. G. Zacaría-Correa 101/IEB |
| *Penstemon pratensis* | 12-0290 | P. D. Blischak 14/OS |
| *Penstemon procerus* var. *procerus* | 12-0216 | P. D. Blischak 3/OS |
| *Penstemon procumbens* | 11-0747 | A. J. Wenzel 23/OS |
| *Penstemon pruinosus* | 16-0157 | E. C. Moran s.n./BRY-V 98608 |
| *Penstemon pseudoputus* | 99-1401 | A. D. Wolfe 823/OS |
| *Penstemon pseudospectabilis* var. *connatifolius* | 03-149 | Paul Wilson 3895/SVF |
| *Penstemon pseudospectabilis* var. *pseudospectabilis* | 98-0517 | A. D. Wolfe 767/OS |
| *Penstemon pudicus* | 16-0158 | K. A. Vincent 4425/BRY-V 399062 |
| *Penstemon pumilis* | 15-0707 | Smith 4579/ORED |
| *Penstemon purpusii* | 15-0198 | Howe/ SD 98858 |
| *Penstemon radicosus* | 99-1656 | Mt. West Enviro Services 7962/OS |
| *Penstemon ramaleyi* | 14-685 | Islam 01/OS |
| *Penstemon reidmoranii* | 14-0070 | A. D. Wolfe s.n./OS |
| *Penstemon retrorsus* | 99-0934 | Arft 9-51/COLO |
| *Penstemon rhizomatus* | 16-0159 | Arnold Tiehm 12692/BRY-V 403695 |
| *Penstemon richardsonii* | 96-0071 | A. D. Wolfe 585/OS |
| *Penstemon roezlii* | 16-0001 | A. D. Wolfe 1495/OS |
| *Penstemon roseus* | 14-0081 | A. G. Zacaría-Correa 88/IEB |
| *Penstemon rostriflorus* | 99-1387 | A. D. Wolfe 809/OS |
| *Penstemon rubicundus* | 99-1467 | A. D. Wolfe 853/OS |
| *Penstemon rupicola* | 96-0035 | A. D. Wolfe 575/OS |
| *Penstemon rydbergii* var. *oreocharis* | 12-0319 | P. D. Blischak 16/OS |
| *Penstemon rydbergii* var*. rydbergii* | 99-0034 | J. Thomson s.n./OS |
| *Penstemon saltarius* | 14-0102 | A. G. Zacaría-Correa 112/IEB |
| *Penstemon saxosorum* | 99-1759 | Mt. West Enviro Services 7981/OS |
| *Penstemon scapoides* | 99-1392 | A. D. Wolfe 814/OS |
| *Penstemon scariosus* var. *albifluvis* | 15-0249 | A. D. Wolfe 1449/OS |
| *Penstemon scariosus* var*. cyanomontanus* | 241 | M. Stevens/BRY-V 133610 |
| *Penstemon scariosus* var*. garrettii* | 15-0213 | A. D. Wolfe 1440/OS |
| *Penstemon secundiflorus* | 99-1754 | Mt. West Enviro Services 7954/OS |
| *Penstemon seorsus* | 15-0709 | Mansfield 07127/ORED |
| *Penstemon sepalulus* | 16-0160 | Welsh 638/BRY-V 221799 |
| *Penstemon serrulatus* | 96-0031 | A. D. Wolfe 610/OS |
| *Penstemon smallii* | 96-0088 | Lindgren 8/NEB |
| *Penstemon spatulatus* | 15-0666 | P. D. Blischak 56/OS |
| *Penstemon speciosus* | 98-0086 | Paul Wilson 3486/SVF |
| *Penstemon spectabilis* var*. spectabilis* | 30 | A. D. Wolfe 223/OS |
| *Penstemon spectabilis* var*. subviscosus* | 79 | A. D. Wolfe 323/OS |
| *Penstemon stenophyllus* | 15-0193 | Leon de la Luz/SD 146780 |
| *Penstemon strictiformus* | 15-0341 | A. D. Wolfe 1478/OS |
| *Penstemon strictus* | 16-0041 | A. D. Wolfe 1501/OS |
| *Penstemon subglaber* | 99-1747 | Mt. West Enviro Services 7989/OS |
| *Penstemon subserratus* | 17-0005 | A. D. Wolfe 590/OS |
| *Penstemon subulatus* | 01-0096 | Paul Wilson s.n./SVF |

| | | |
|---|---|---|
| *Penstemon sudans* | 16-0079 | A. D. Wolfe 1511/OS |
| *Penstemon superbus* | 02-0464 | Mt. West Enviro Services 9094/OS |
| *Penstemon tenuifolius* | 14-0108 | A. G. Zacaría-Correa 104/IEB |
| *Penstemon tenuis* | 99-1447 | A. D. Wolfe 833/OS |
| *Penstemon teucrioides* | 11-0844 | A. J. Wenzel 29/OS |
| *Penstemon thompsoniae* var. *jaegeri* | 12-0082 | A. J. Wenzel 43/OS |
| *Penstemon thompsoniae* var. *thompsoniae* | 15-0300 | A. D. Wolfe 1463/OS |
| *Penstemon tracyi* | 16-0047 | A. D. Wolfe 1503/OS |
| *Penstemon triflorus* var. *integrifolius* | 16-0169 | Andres dz., L. Arce, M. Martinez M. 1339/TEX |
| *Penstemon triphyllus* | 17-0002 | A. D. Wolfe 651/OS |
| *Penstemon tubaeflorus* | 17-0009 | A. D. Wolfe 271/OS |
| *Penstemon tusharensis* | 12-0156 | A. J. Wenzel 49/OS |
| *Penstemon uintahensis* | 10-0183 | Mt. West Enviro Services 8938/OS |
| *Penstemon utahensis* | 15-0293 | A. D. Wolfe 1461/OS |
| *Penstemon venustus* | 96-0002 | A. D. Wolfe 646/OS |
| *Penstemon versicolor* | 97-0332 | Stermitz 527A/CMML |
| *Penstemon virens* | 99-1745 | Mt. West Enviro Services 7953/OS |
| *Penstemon virgatus* | 17-0007 | A. D. Wolfe 829/OS |
| *Penstemon virgatus* var. *asa-grayi* | 99-1743 | Mt. West Enviro Services 8057/OS |
| *Penstemon vizcainensis* | 15-0200 | Moran/SD 104945 |
| *Penstemon vulcanellus* | 14-0106 | A. G. Zacaría-Correa 111/IEB |
| *Penstemon washingtonensis* | 16-0173 | P. D. Blischak 36/OS |
| *Penstemon watsonii* | 98-0489 | A. D. Wolfe 786/OS |
| *Penstemon wendtiorum* | 14-0003 | A. G. Zacaría-Correa 97/IEB |
| *Penstemon whippleanus* | 96-0015 | Ranker 1739 (COLO) |
| *Penstemon wilcoxii* | 15-0451 | P. D. Blischak 39/OS |
| *Penstemon wislizeni* | 14-0087 | A. G. Zacaría-Correa 78/IEB |
| *Penstemon wrightii* | 99-1403 | A. D. Wolfe 824/OS |
| *Penstemon X jonesii* | 15-0316 | A. D. Wolfe 1470/OS |
| *Penstemon X parishii* | 150 | A. D. Wolfe 224/OS |
