## Appendix 2 for "Phylogenetics of a Rapid, Continental Radiation: Diversification, Biogeography, and Circumscription of the Beardtongues (*Penstemon*; Plantaginaceae)"

### Notes on Filtering Individuals and Loci in Geneious

Below are notes on the steps that were used to filter individual samples and loci to create our final data set. The initial, completely unfiltered data are available in the ‘UnfilteredFasta/’ folder and the filtered data are available in the ‘FilteredFasta/’ folder. These notes are to help explain how we went from the unfiltered data set to the filtered set.

##### *Removing Loci*

Loci with an insufficient coverage of the taxon set were removed. This included locus *1331*, *21370*, *trnCD*, *trnTL*, and *rps12rpl20*. Locus *2919* looked a little dodgy for some taxa but we left them in because there was no obvious signal for their sequences being too short or completely misaligned. For loci with lower numbers of taxa recovered (eg, 100 or less), we kept loci if they had a decent representation of taxa from different groups or if they had outgroups represented. For example, many of the less successfully amplified loci only worked in the subgenus *Dasanthera* and nothing else, so we removed them.

##### *Removing Taxa*

After aligning with Muscle in Geneious, the sequences are sorted such that poorly aligned sequences “float” to the top of the alignment being viewed (I think it is because they have the lowest pairwise identity, or lowest identity shared with the consensus sequence). These sequences are always too short, meaning that they were poorly assembled or had poor quality read data, so they were removed.
