## Supplementary figures and images for "Phylogenetics of a Rapid, Continental Radiation: Diversification, Biogeography, and Circumscription of the Beardtongues (*Penstemon*; Plantaginaceae)"

### Figure S1

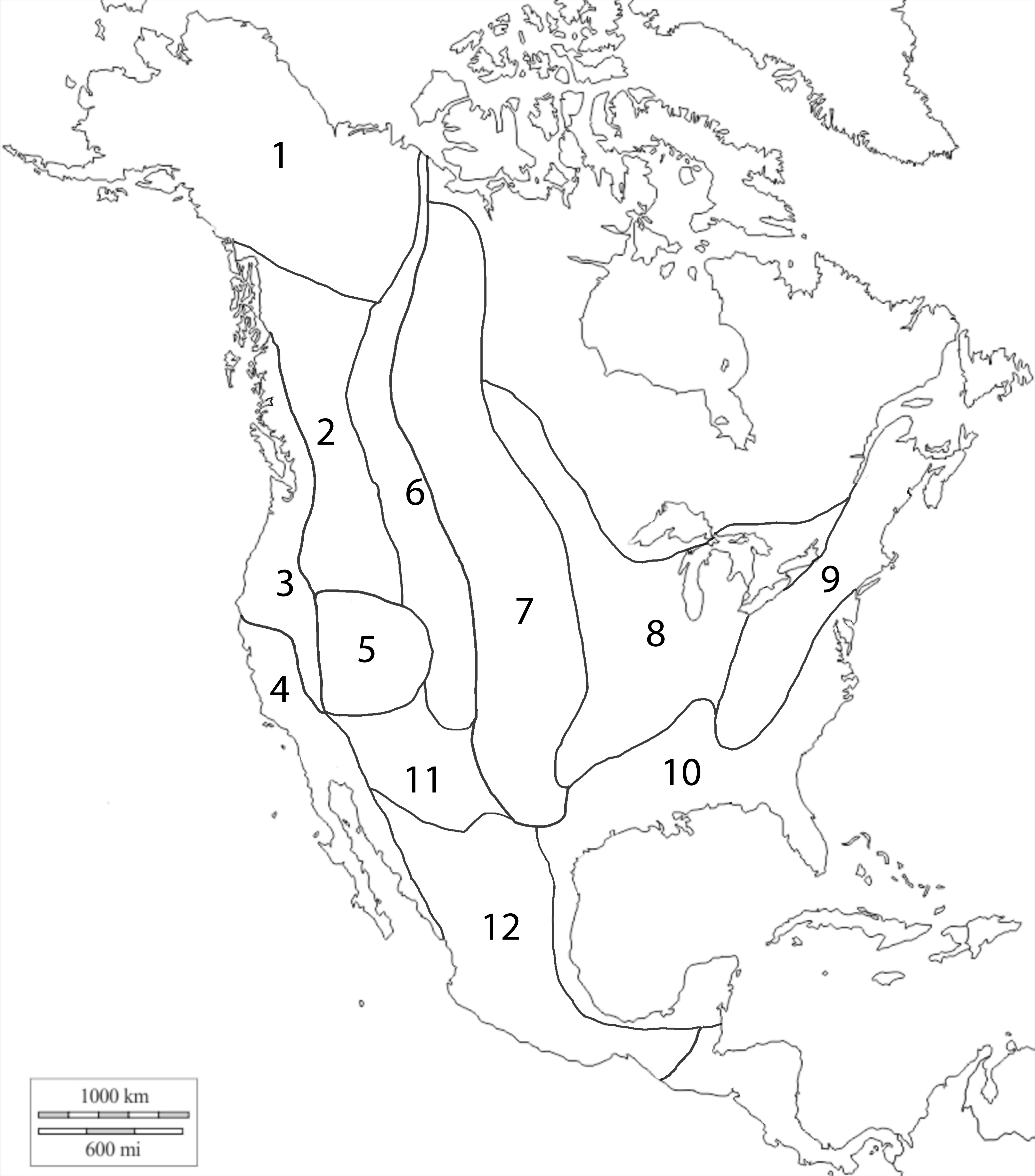

### Figure S2

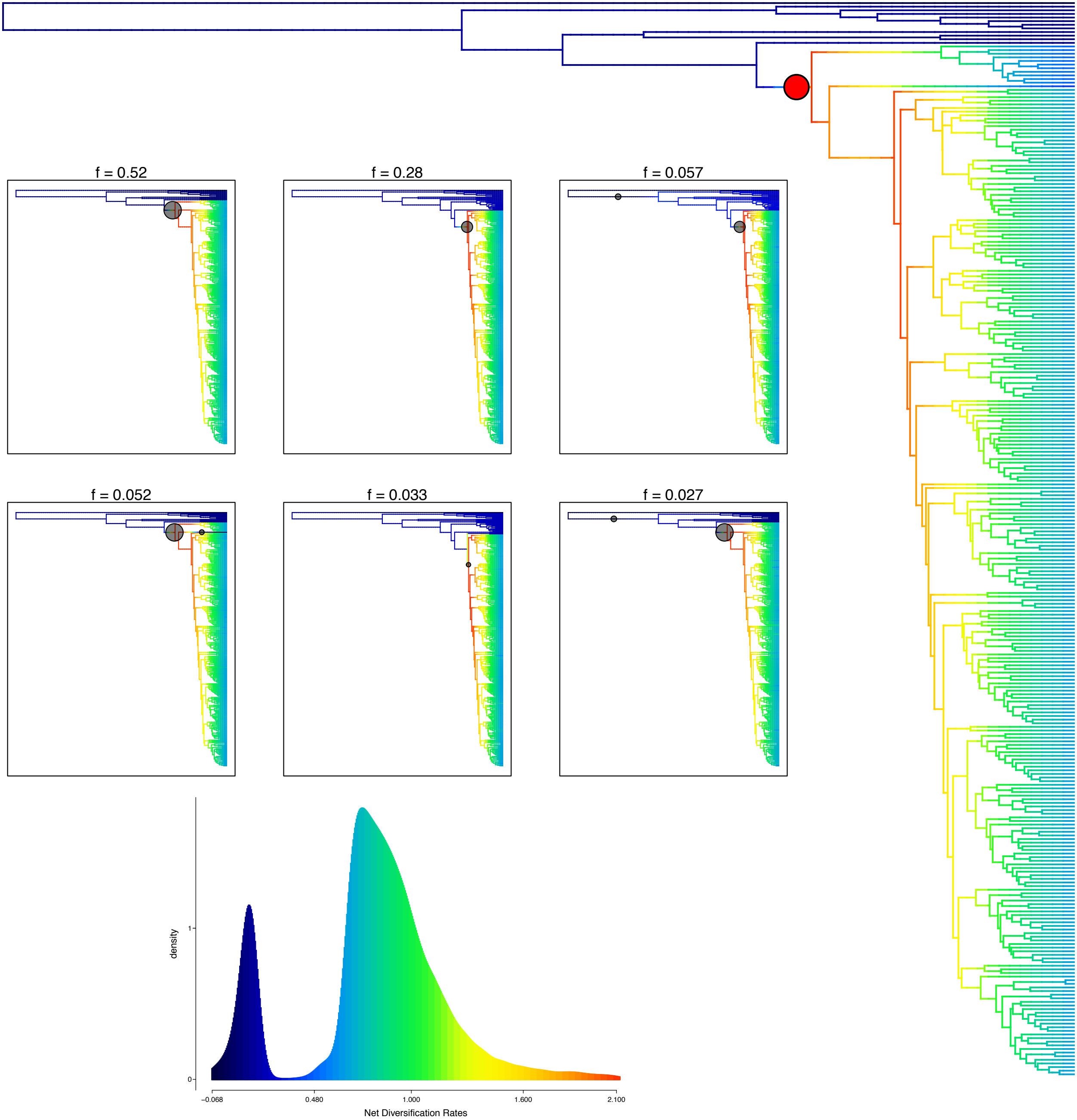

### Figure S6

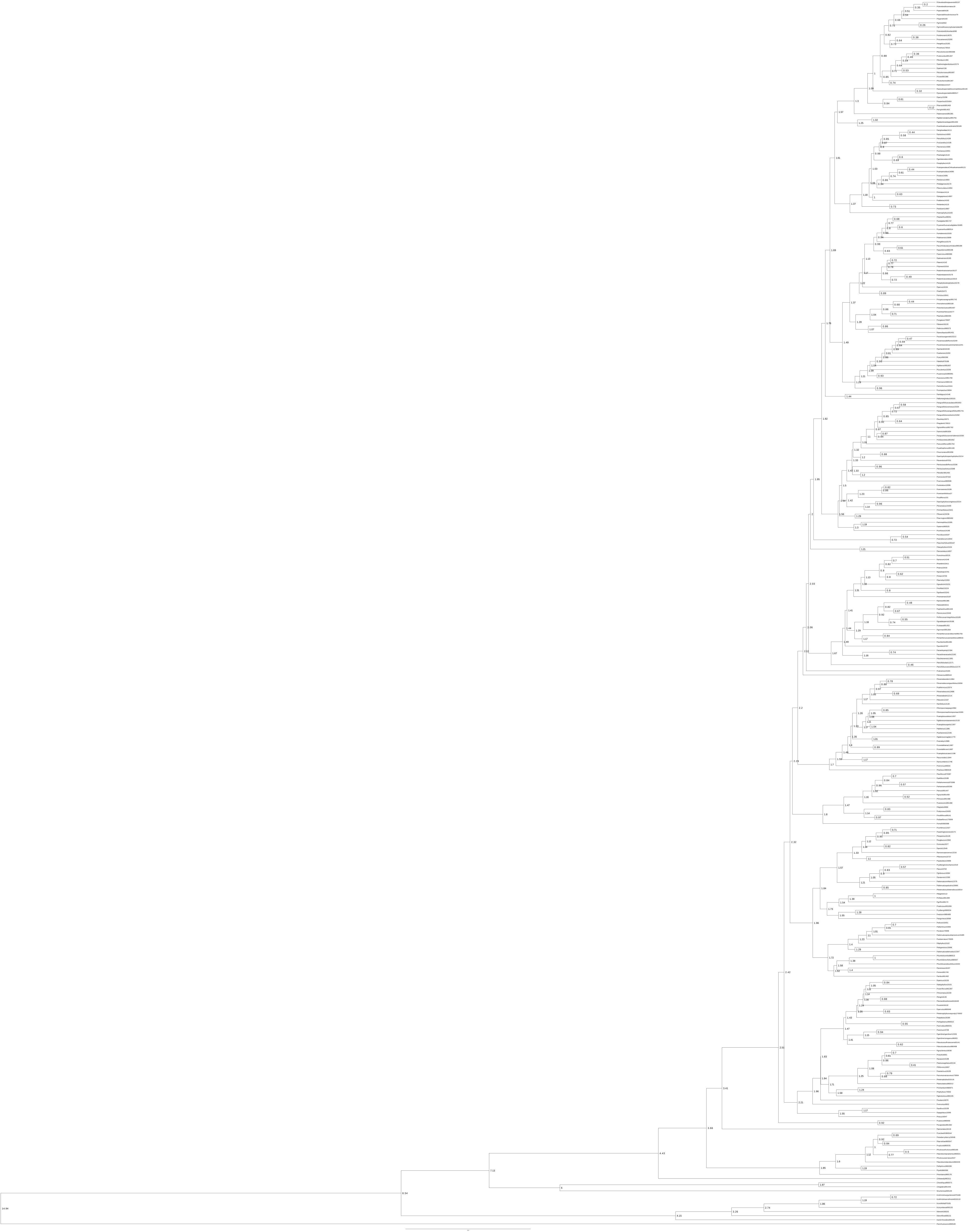

### Figure S7

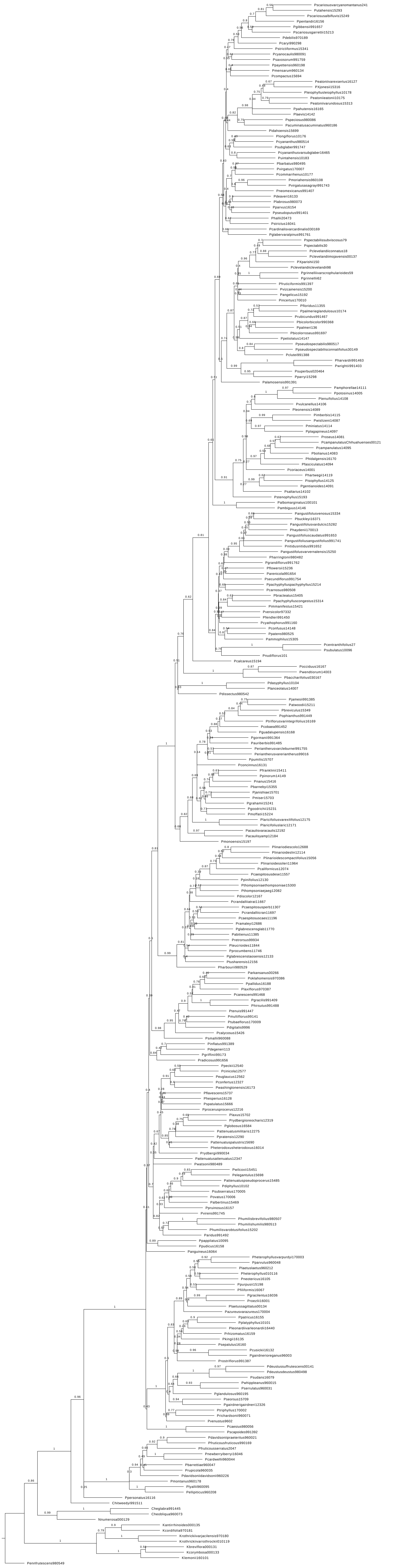
