## Supplementary material for "Phylogenetics of a Rapid, Continental Radiation: Diversification, Biogeography, and Circumscription of the Beardtongues (*Penstemon*; Plantaginaceae)": Figure S3

**BioGeoBEARS BAYAREALIKE+J on Penstemon set1 M0_unconstrained**
**ancstates: global optim, 3 areas max. d=0.0187; e=0; j=0.0394; LnL=−376.84**

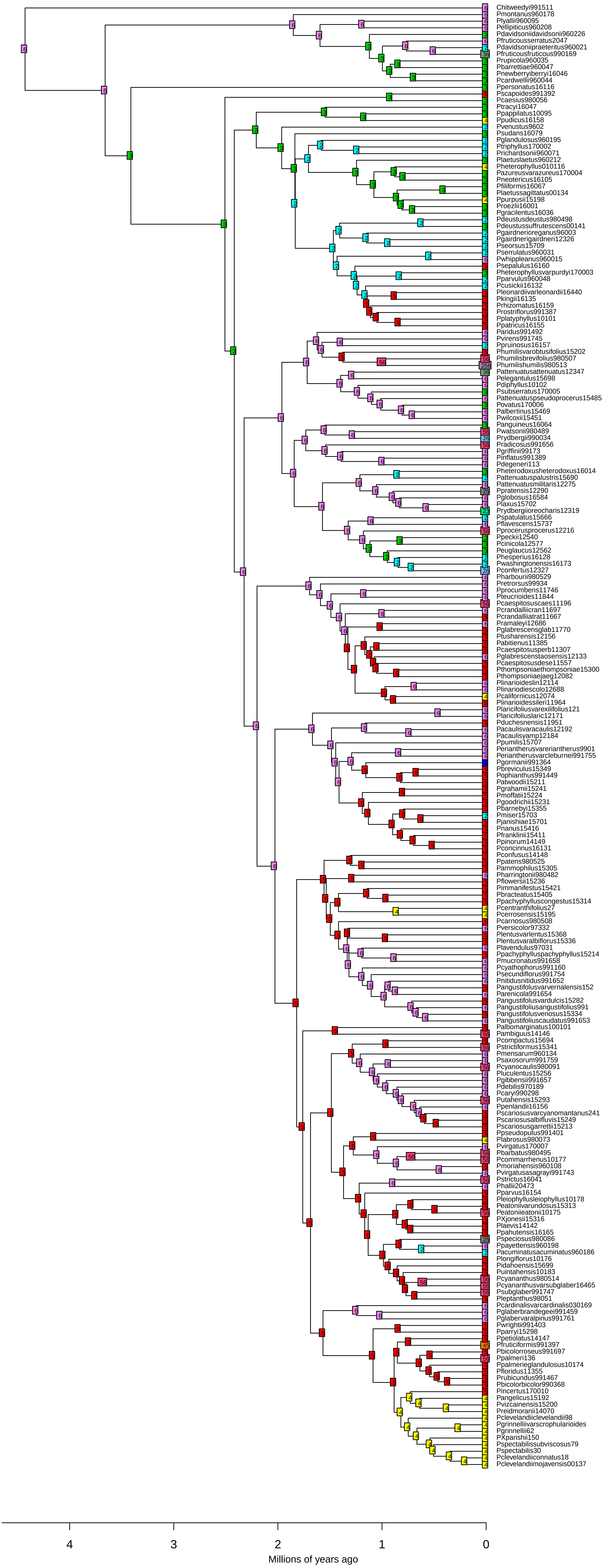

### BioGeoBEARS BAYAREALIKE+J on Penstemon set2 M0_unconstrained
### ancstates: global optim, 4 areas max. d=0.0321; e=0; j=0.0329; LnL=−424.02

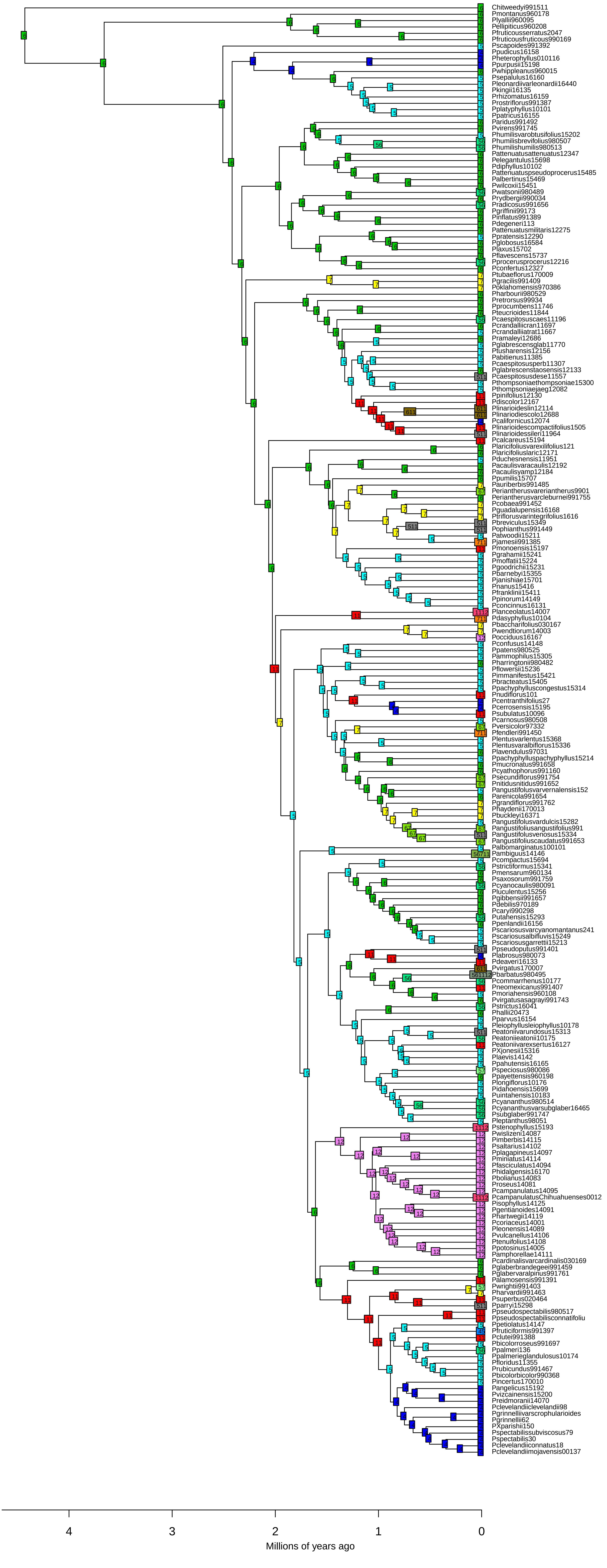

**BioGeoBEARS BAYAREALIKE+J on Penstemon set3 M0_unconstrained**
**ancstates: global optim, 3 areas max. d=0.0194; e=0; j=0.0303; LnL=−216.75**

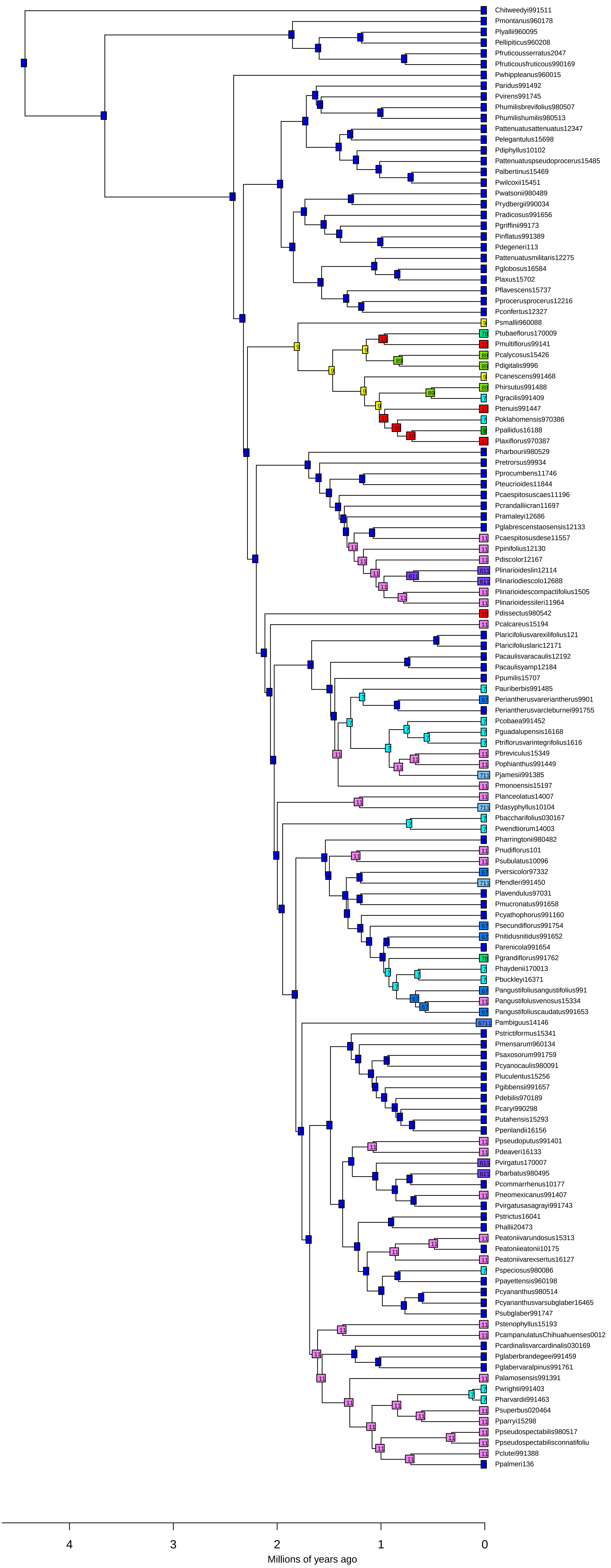
