## Supplementary material for "Phylogenetics of a Rapid, Continental Radiation: Diversification, Biogeography, and Circumscription of the Beardtongues (*Penstemon*; Plantaginaceae)": Figure S4

**Fig. S4**. Distribution of penstemons in chronological order from the Pliocene/Pleistocene border to present. Refer to Table S2 for biogeographic chronology based on time-calibrated tree.

#### Pliocene to 2.5 mya

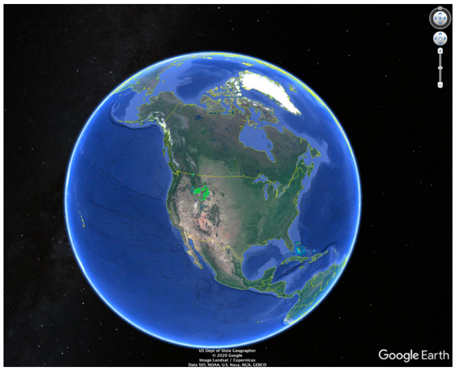

*Penstemon montanus*

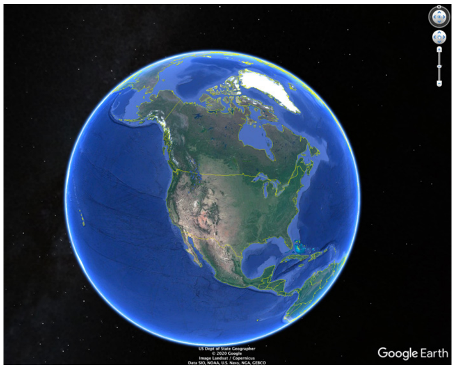

*Penstemon lyallii*

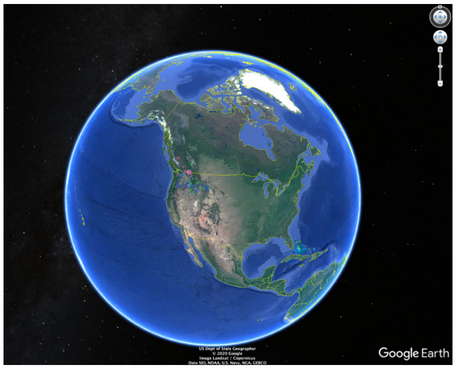

*Penstemon fruticosus*

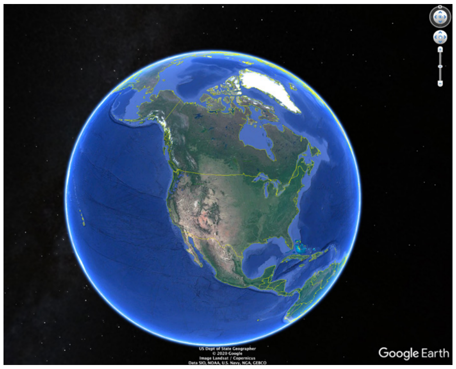

*Penstemon rupicola, P. barrettiae*

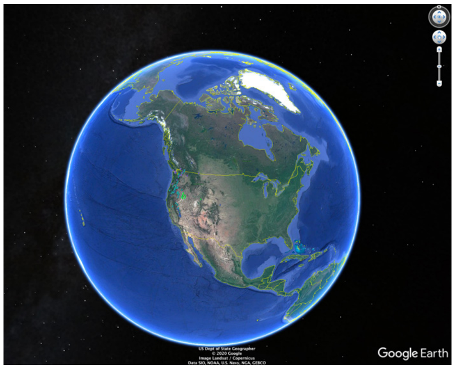

*Penstemon davidsonii*

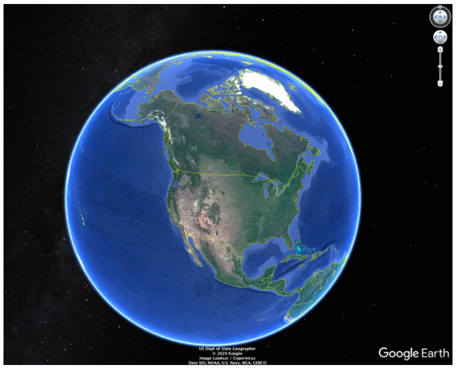

*Penstemon cardwellii*

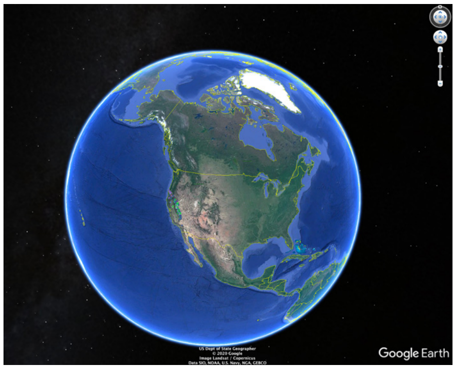

*Penstemon newberryi*

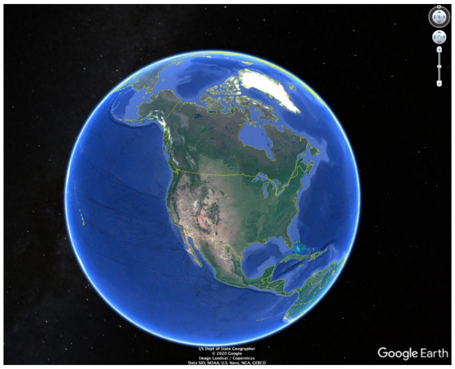

*Penstemon personatus*

#### 2.5 mya to 2.0 mya

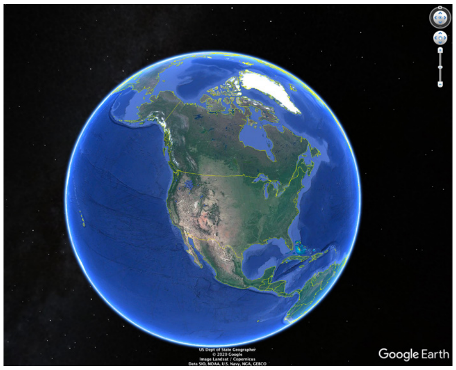

*Penstemon venustus*

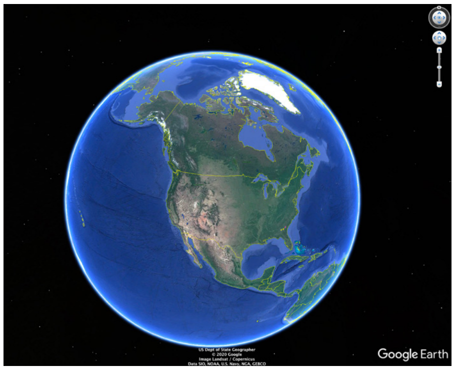

*Penstemon dissectus, P. calcareus*

#### 2.0 mya to 1.8 mya

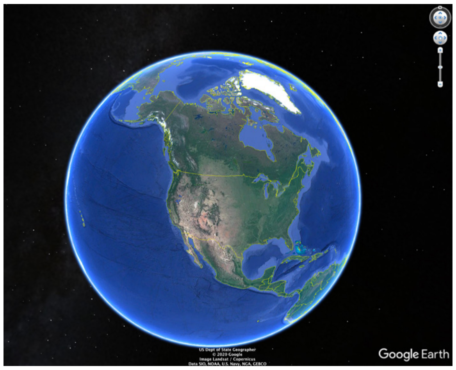


*Penstemon sudans*

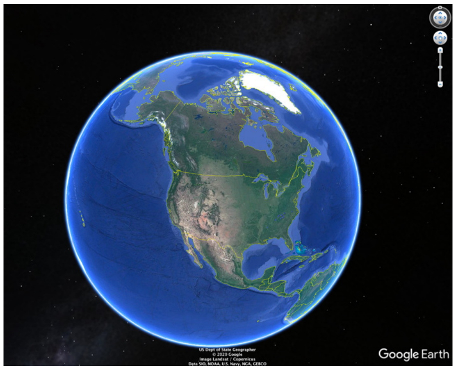


*Penstemon smallii*

#### 1.8 mya to 1.7 mya

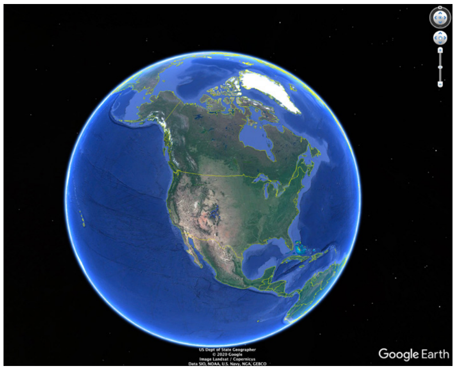


*Penstemon harbourii, retorsus*

#### 1.7 mya to 1.6 mya

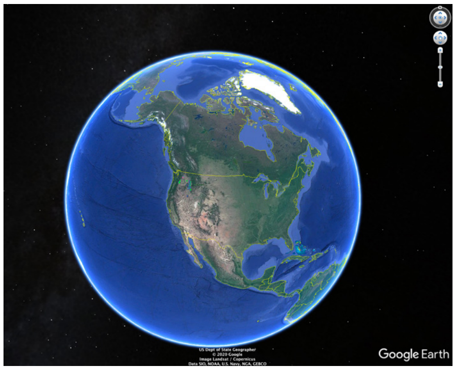


*Penstemon glandulosus*

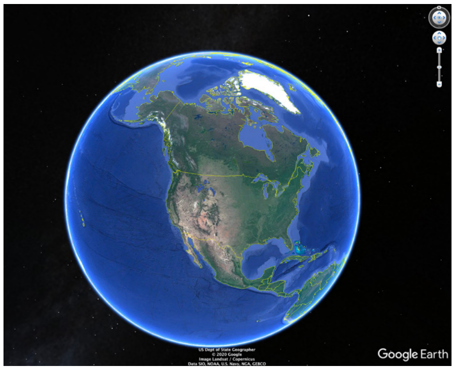


*Penstemon aridus*

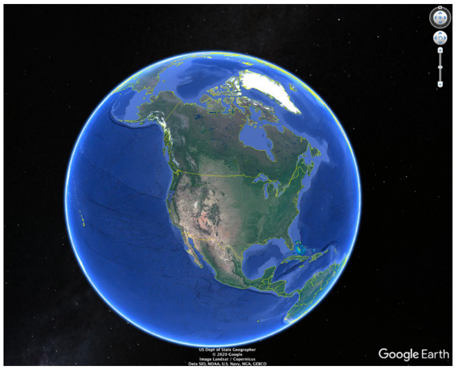


*Penstemon anguineus*

#### 1.6 mya to 1.5 mya

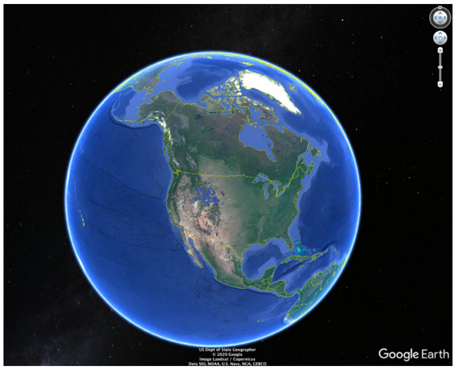


*Penstemon radicosus*

### 1.5 mya to 1.4 mya

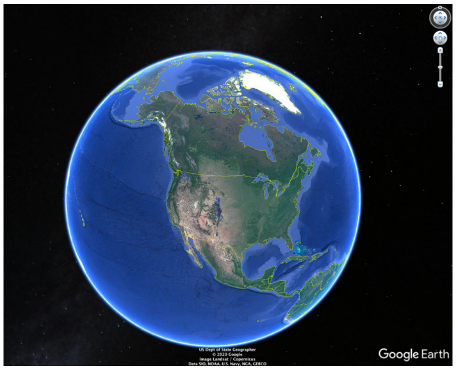

*Penstemon virens, P.pruinosus*

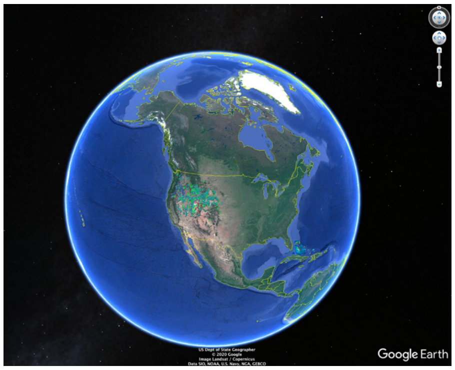

*Penstemon humilis*

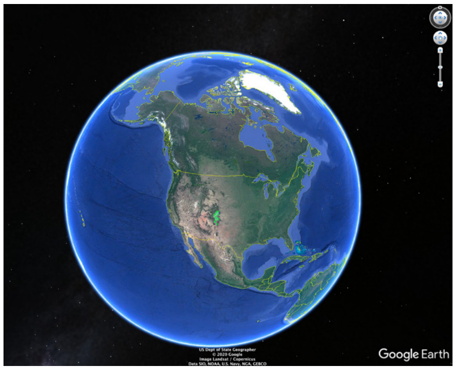

*Penstemon griffinii*

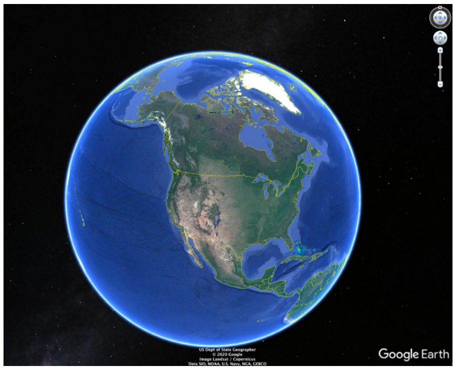

*Penstemon inflatus, P. degeneri*

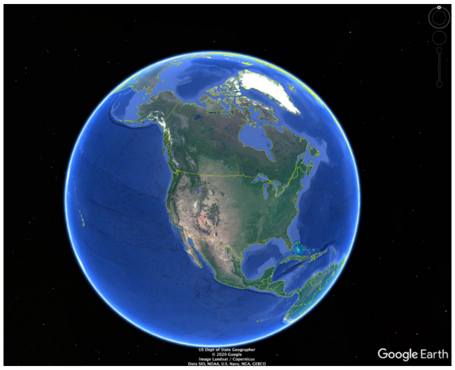

*Penstemon pumilis*

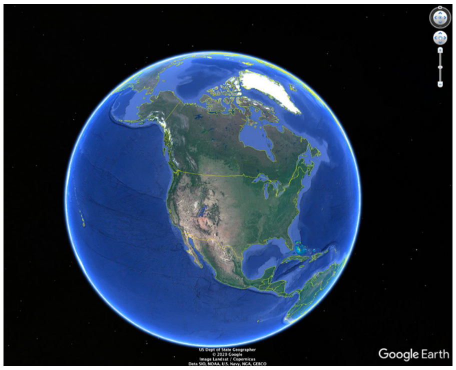

*Penstemon carnosus*

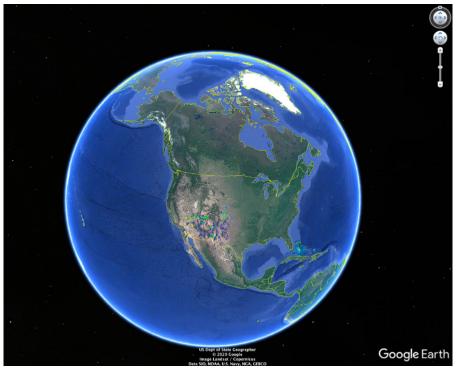

*Penstemon ambiguus, P. albomarginatus*

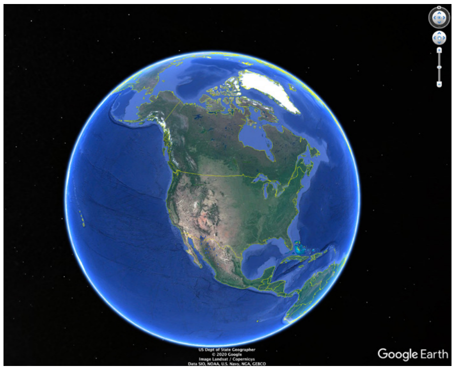

*Penstemon stenophyllus*

#### 1.4 mya to 1.3 mya

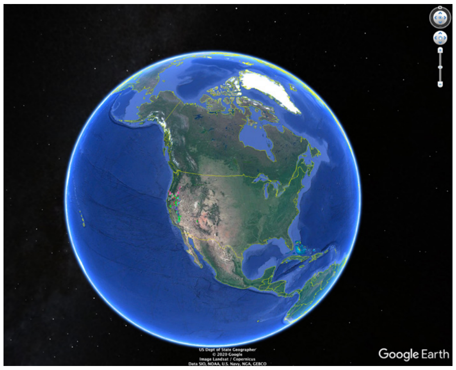


*Penstemon laetus*

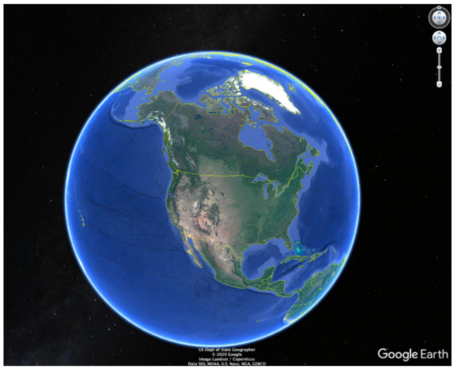


*Penstemon sepalulus, P. Parvulus*

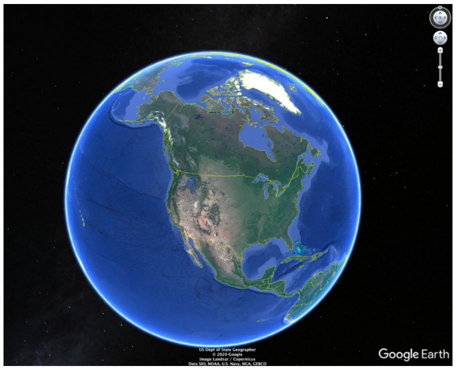


*Penstemon attenuatus* var. *attenuatus, P. elegantulus*

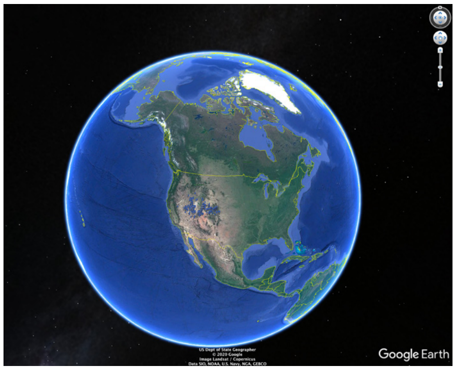


*Penstemon watsonii*

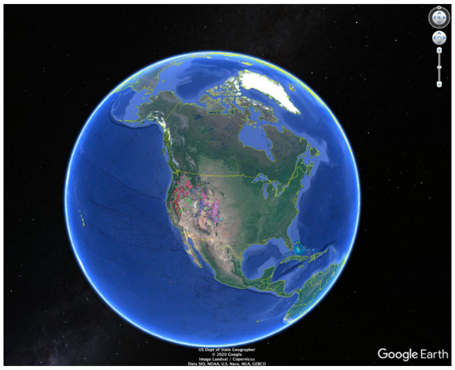


*Penstemon rydbergii*

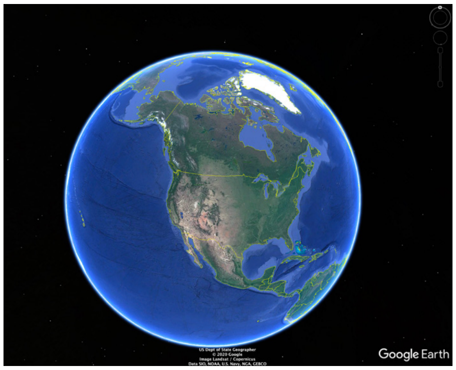


*Penstemon monoensis*

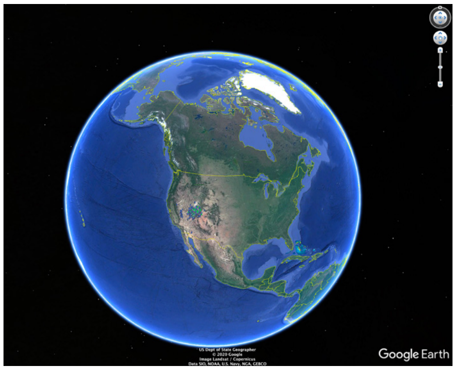


*Penstemon confusus, P. patens, P. ammopholis*

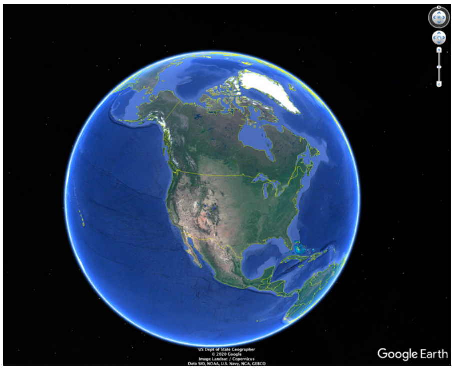


*Penstemon flowersii, P. harringtonii*

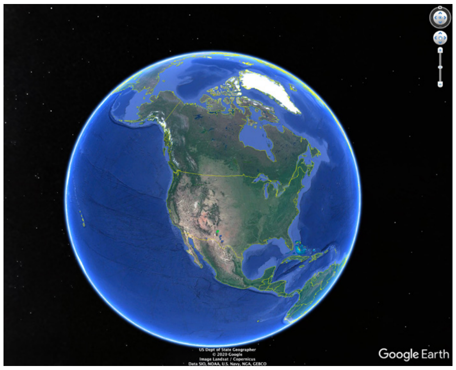


*Penstemon cardinalis*

#### 1.3 mya to 1.2 mya

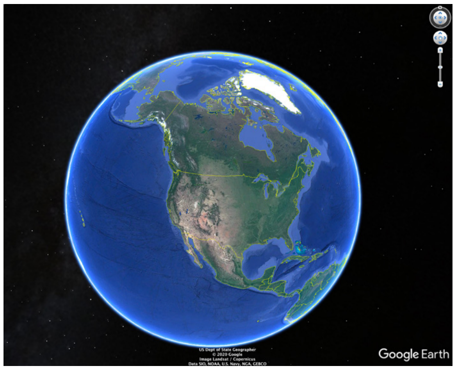


*Penstemon pappilatus, P. pudicus*

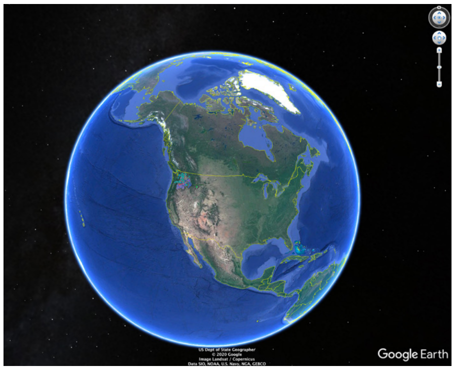


*Penstemon triphyllus, P. richardsonii*

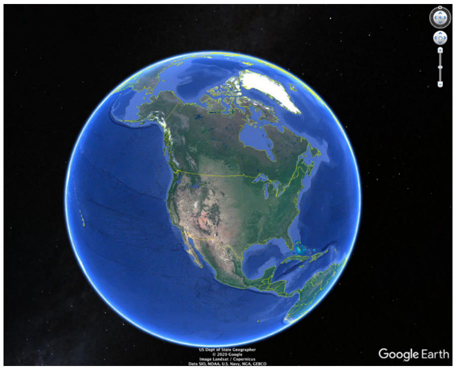


*Penstemon diphyllus, P. subserratus*

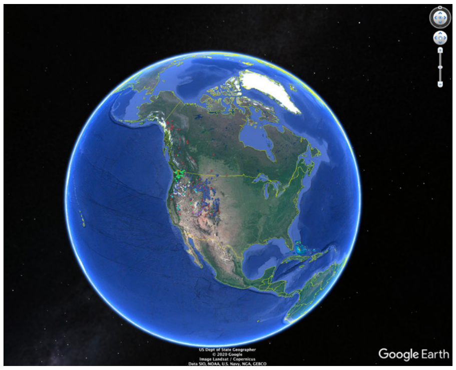

*Penstemon procerus*

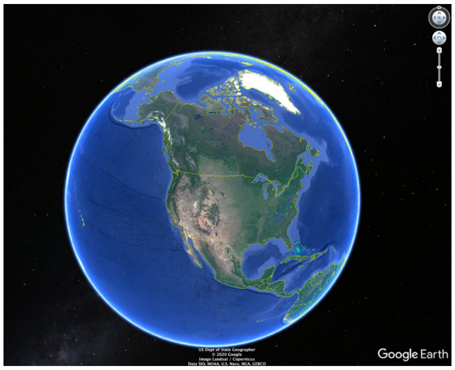

*Penstemon canescens*

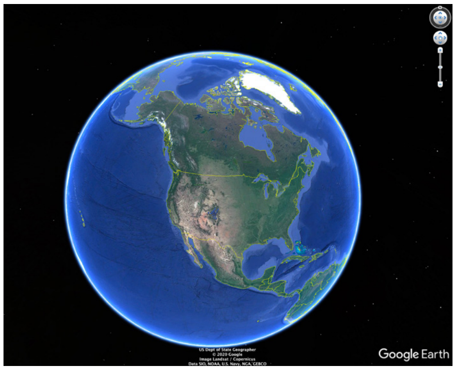

*Penstemon teucrioides, P. procumbens*

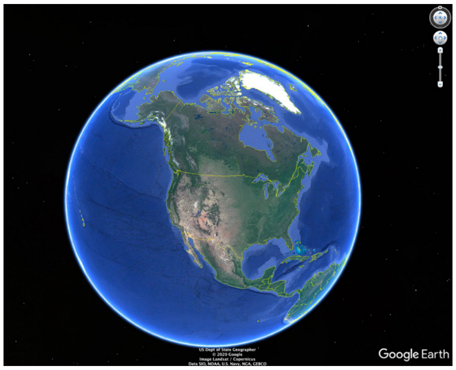

*Penstemon tusharensis*

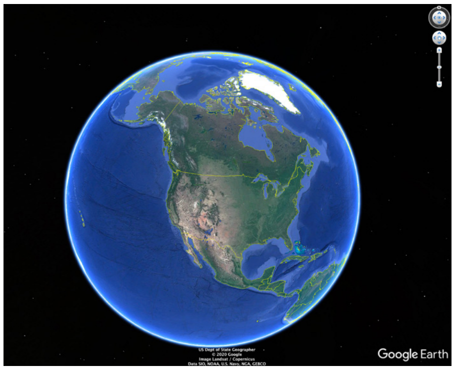

*Penstemon pinifolius*

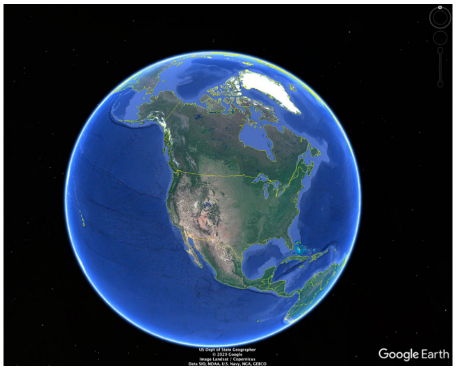

*Penstemon duchesnensis, P. acualis* var. *yampaensis*

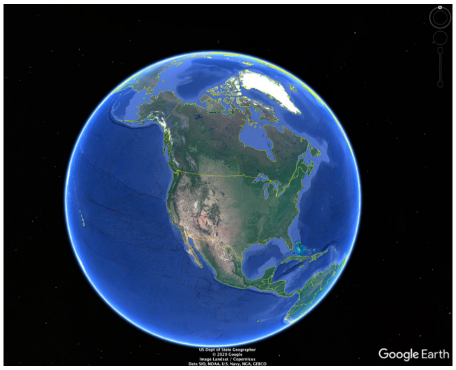

*Penstemon gormanii*

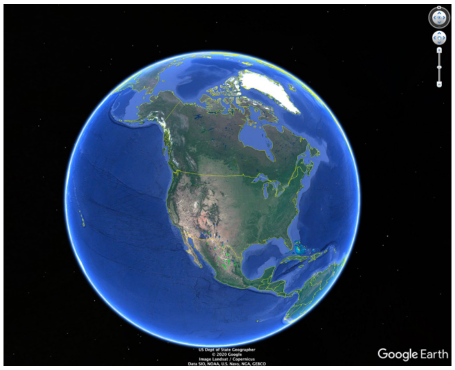

*Penstemon chamaeleon*

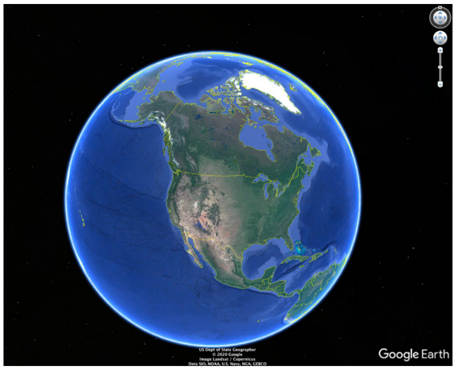

*Penstemon nudiflorus*

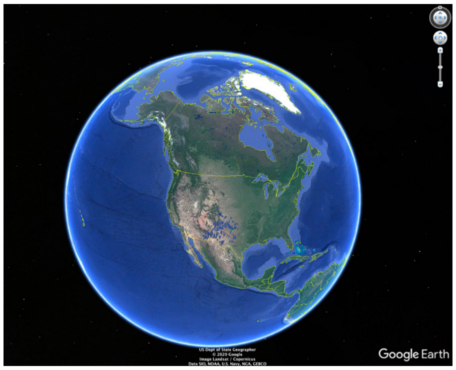

*Penstemon fendleri, P. versicolor*

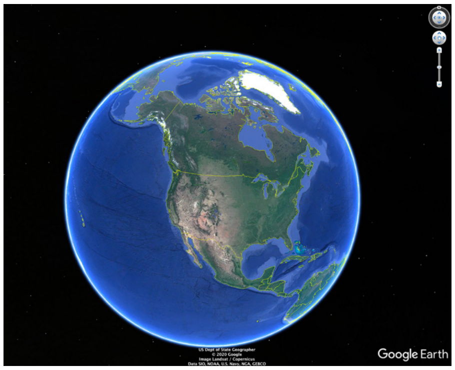

*Penstemon lavendulus*

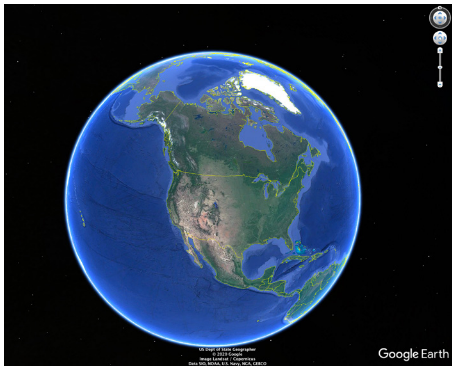

*Penstemon cyathophurus*

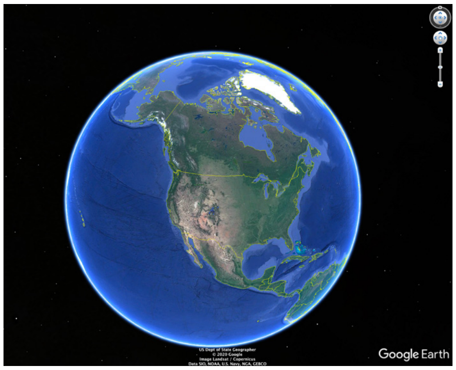


*Penstemon mensarum*

#### 1.2 mya to 1.1 mya

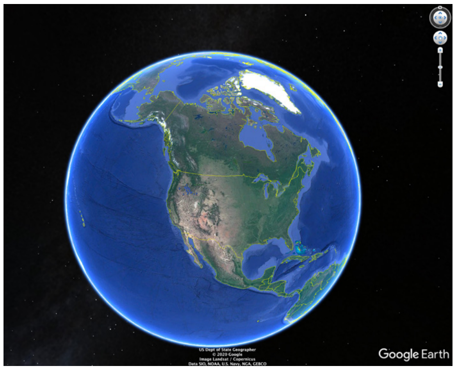


*Penstemon cusickii*

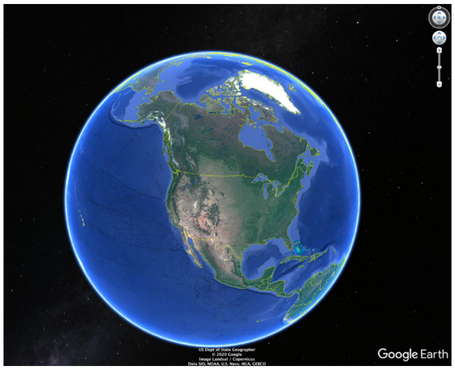


*Penstemon flavescens, P. spatulatus*

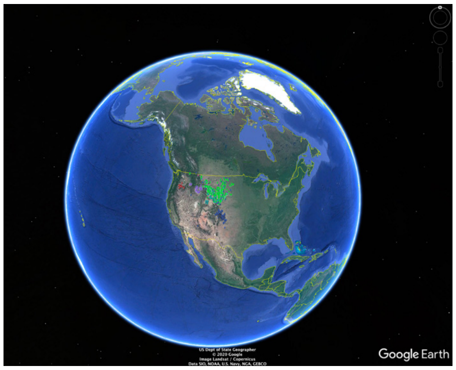


*Penstemon auriberbis, P. eriantherus*

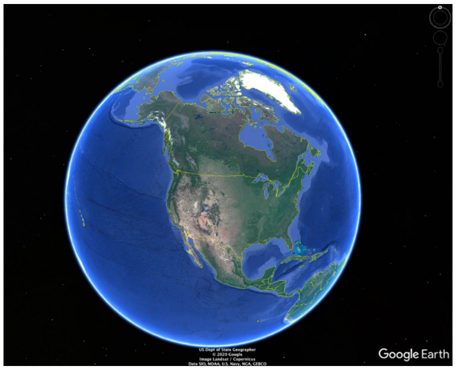


*Penstemon goodrichii, P.marcusii*

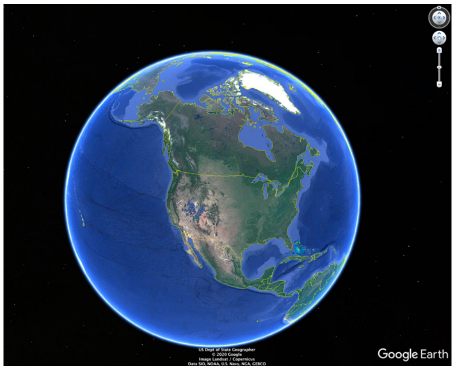


*Penstemon limmanifestus, P. bracteatus, P. pachyphyllus* var. *congestus*

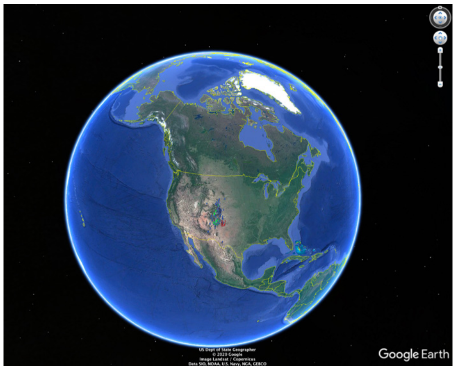


*Penstemon secundiflorus*

#### 1.1 mya to 1.0 mya

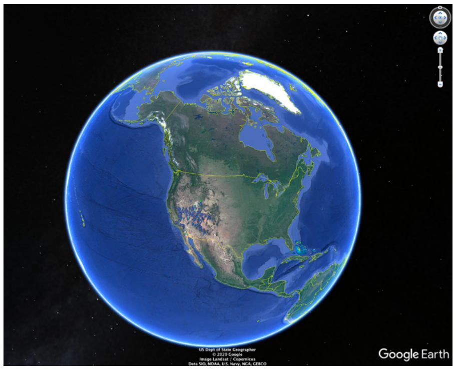


*Penstemon rostriflorus*

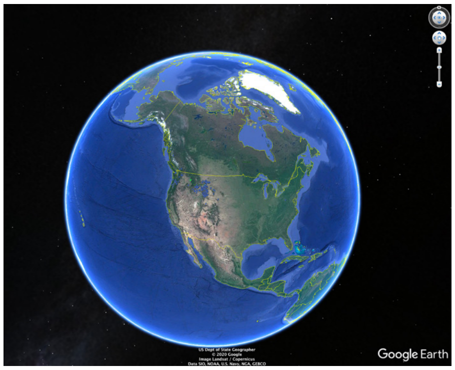


*Penstemon attenuatus* var. *pseudoprocerus, P. ovatus*

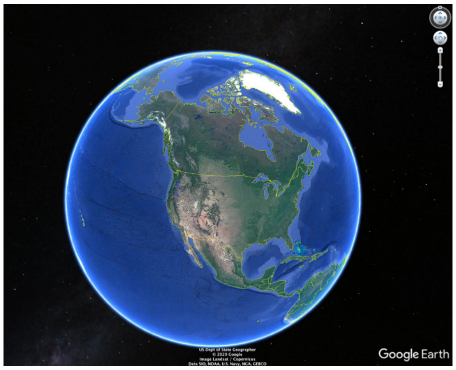


*Penstemon euglaucus*

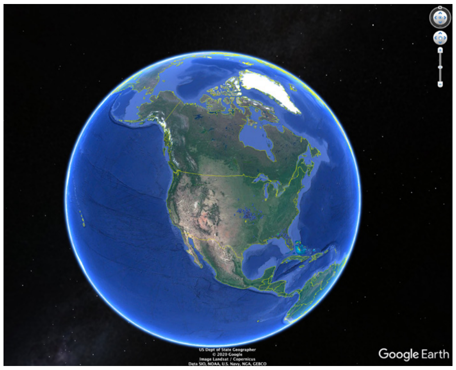

*Penstemon tubaeflorus, P. multiflorus*

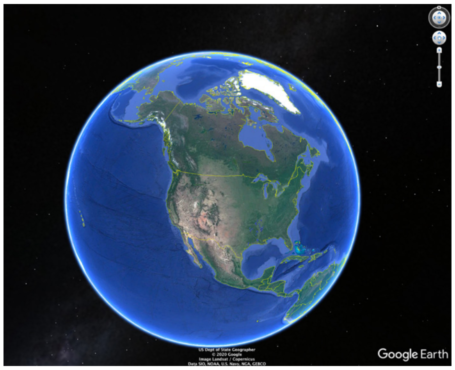

*Penstemon tenuis*

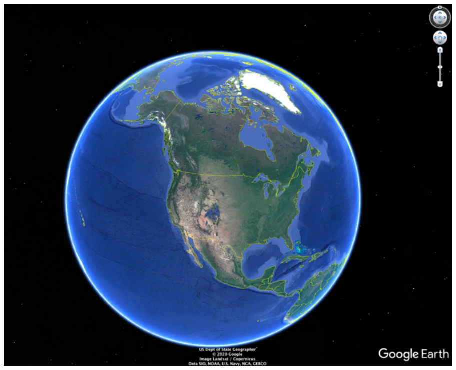

*Penstemon crandallii*

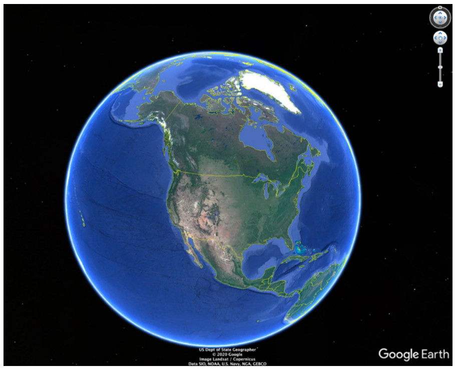

*Penstemon ramaleyi, P. glabrescens*

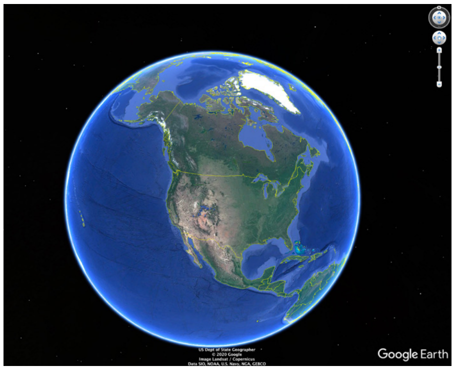

*Penstemon abietinus, P. perbrevis*

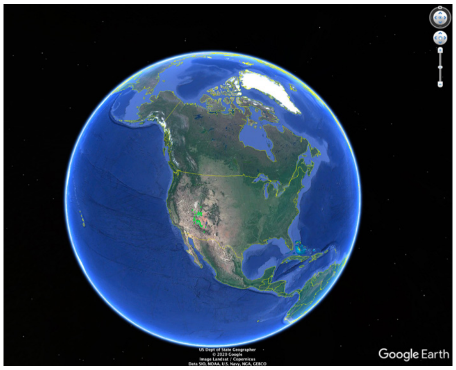

*Penstemon caespitosus* var. *desertipicti*

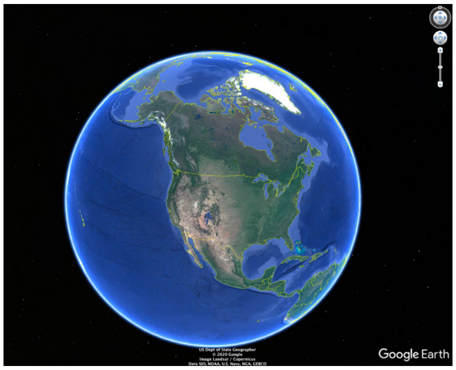

*Penstemon lentus*

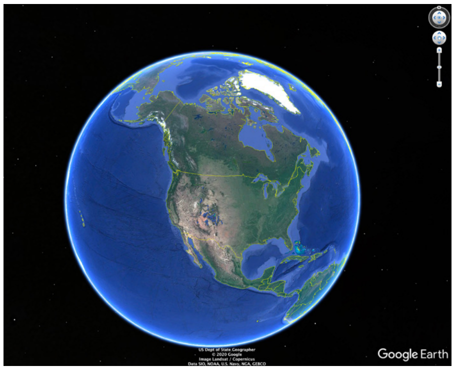

*Penstemon strictiformis, P. compactus*

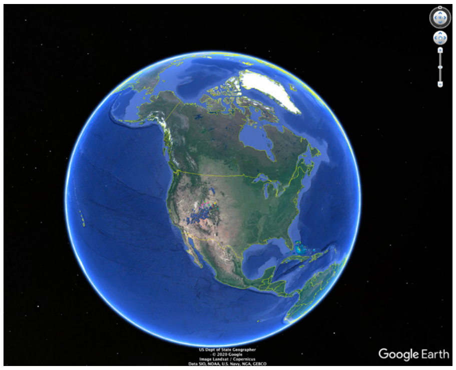

*Penstemon luculentus* clade

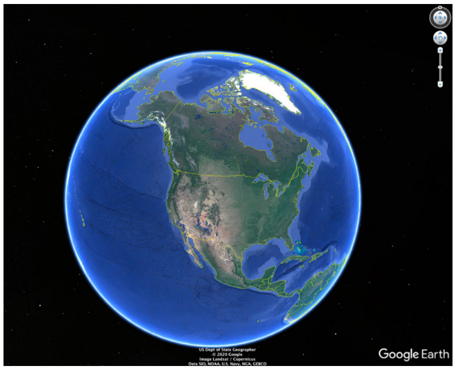

*Penstemon pseudoputus, P. labrosus, P. deaveri*

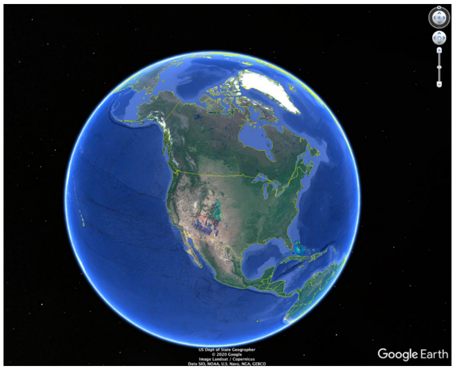

*Penstemon virgatus*

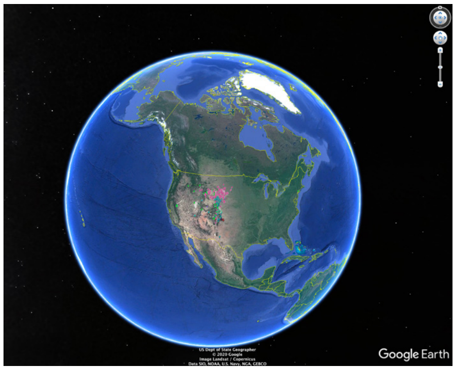

*Penstemon glaber*

#### 1.0 mya to 0.9 mya

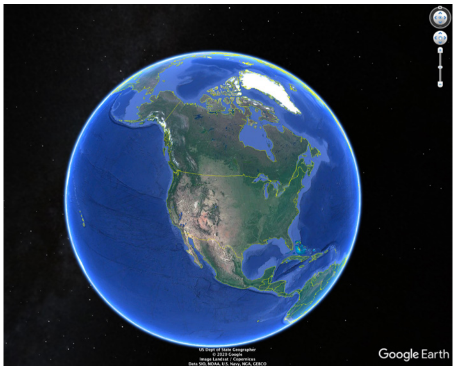

*Penstemon caesius, P. scapoides*

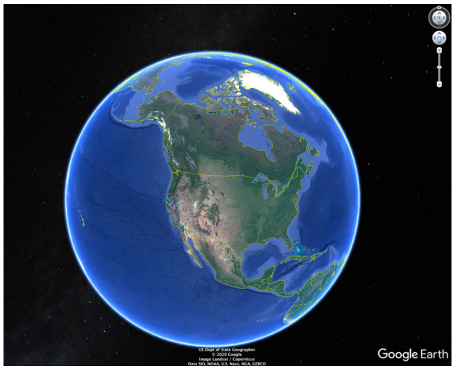

*Penstemon azureus clade*

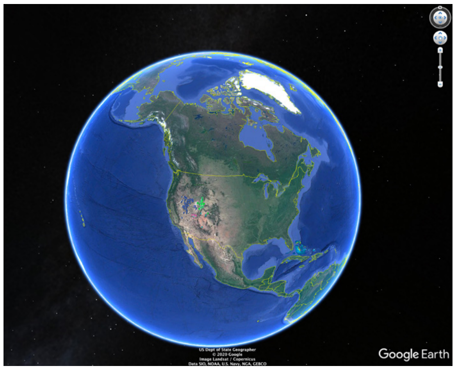

*Penstemon kingii, P. leonardii*

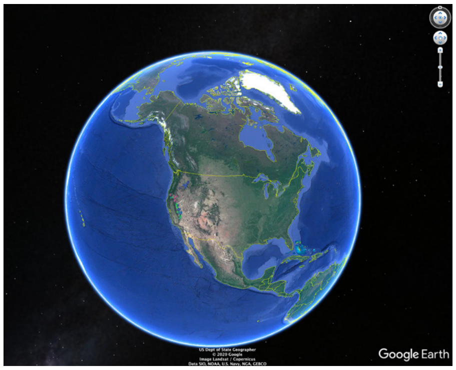

*Penstemon heterodoxus, P. attenuatus* var. *palustris*

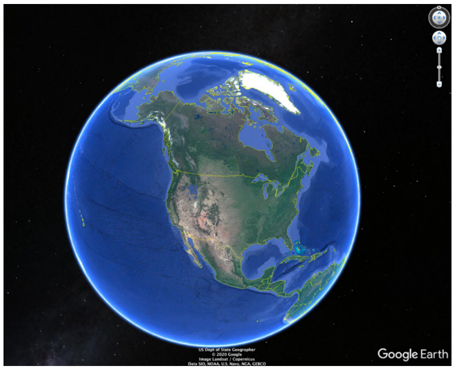

*Penstemon attenuatus* var. *militaris*

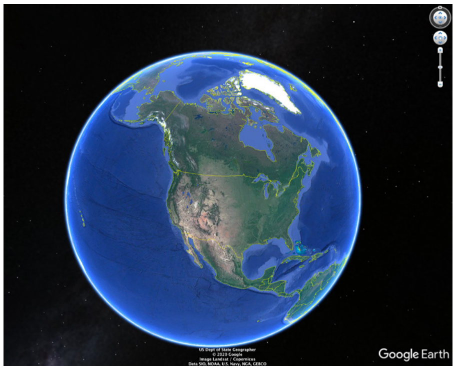

*Penstemon pratensis*

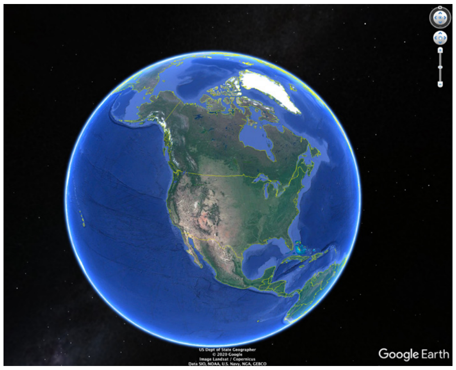

*Penstemon hesperisus, P. washingtonensis*

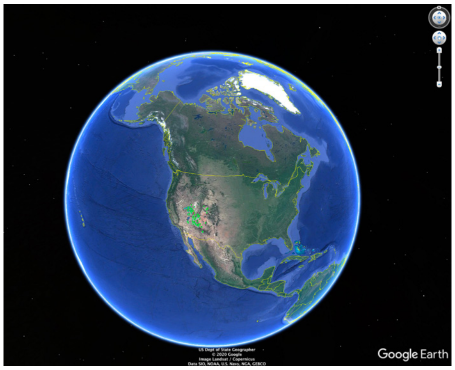

*Penstemon thompsoniae*

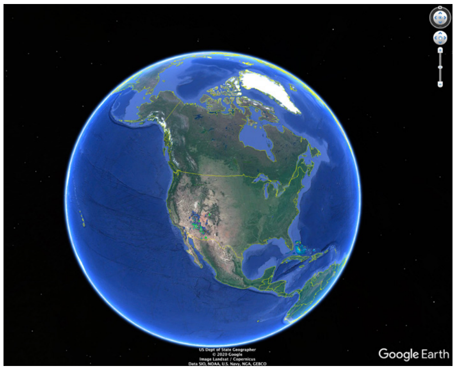

*Penstemon linarioides* var. *sileri, P. compactifolius, P. californicus*

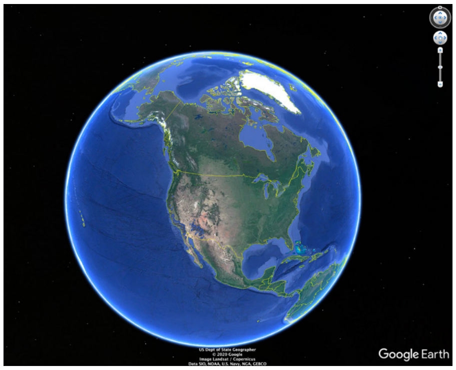

*Penstemon centranthifolius, P. cerrosensis, P. subulatus*

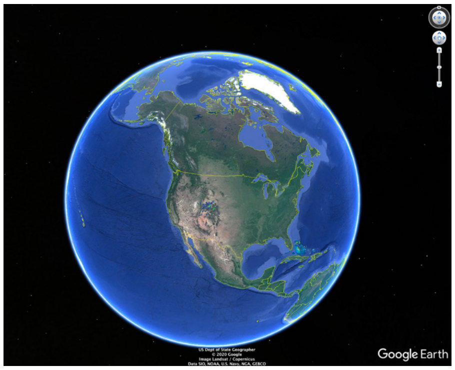

*Penstemon mucronatus, P. pachyphyllus* var. *pachyphyllus*

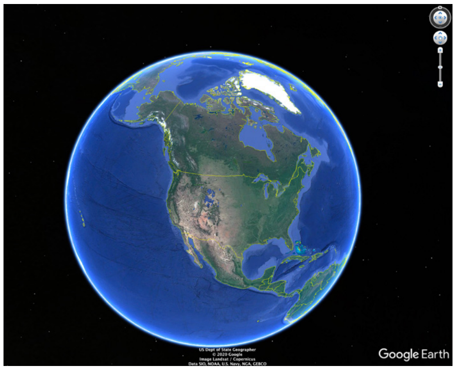

*Penstemon arenicola, P. angustifolius* var. *vernalensis*

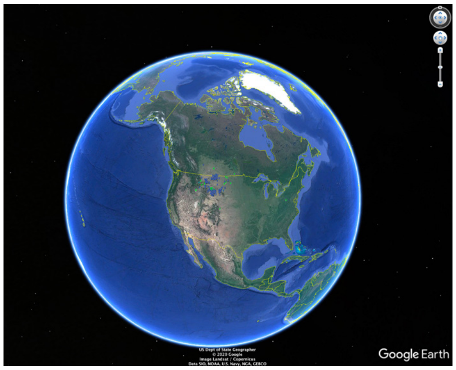

*Penstemon nitidus*

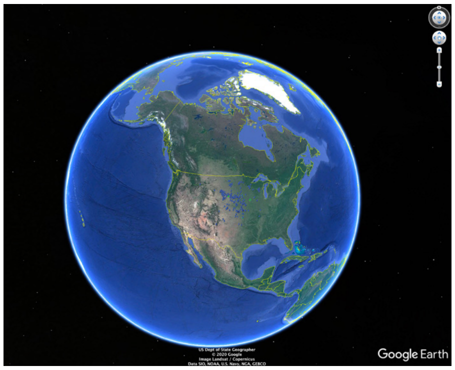

*Penstemon grandiflorus*

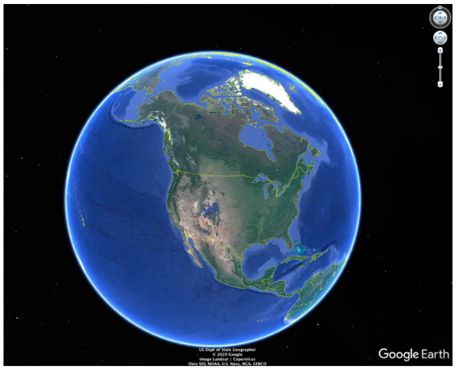

*Penstemon cyanocaulis, P. mensarum*

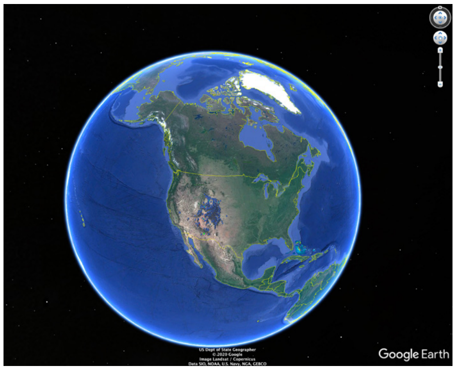

*Penstemon strictus, P. hallii, P. parvus*

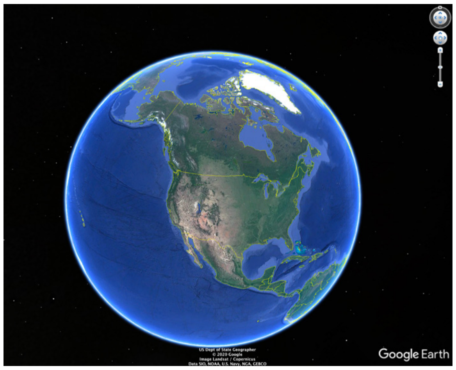

*Penstemon longiflorus*

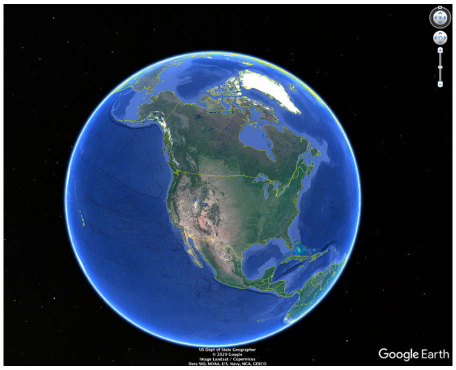

*Penstemon idahoensis, P. uintahensis*

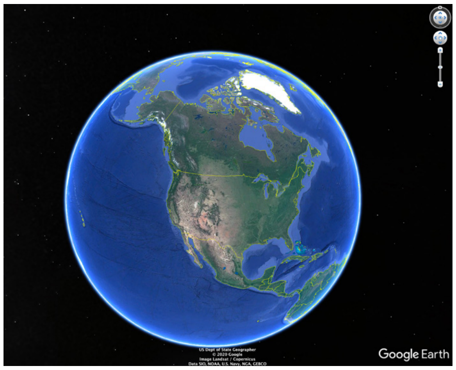

*Penstemon hidalgensis*

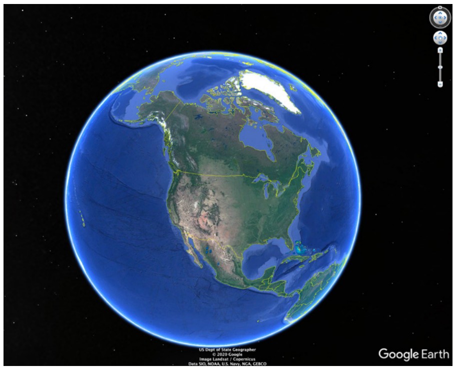

*Penstemon fasciculatus*

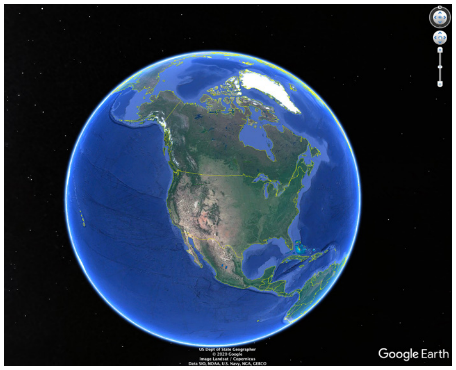

*Penstemon coriaceus*

#### 0.9 mya to 0.8 mya

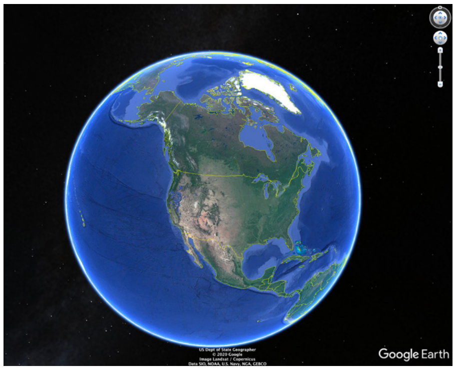

*Penstemon gracilentus* clade

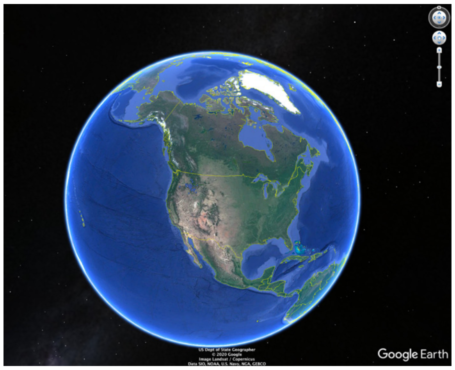

*Penstemon globosus*

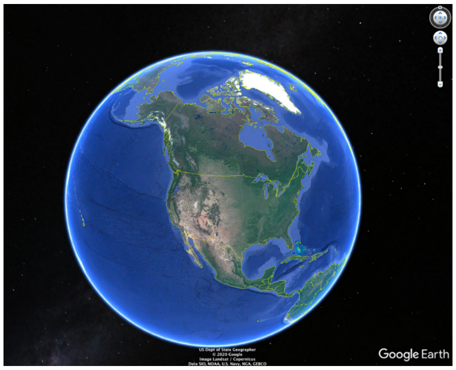

*Penstemon peckii, P. cinicola*

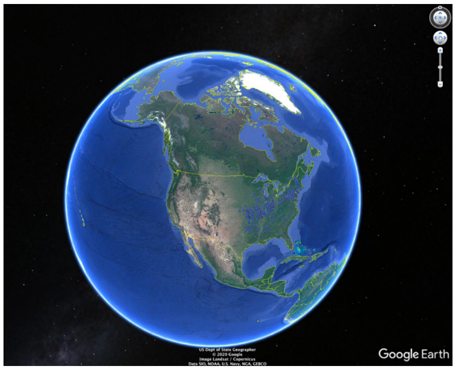

*Penstemon digitalis, P. calycosus*

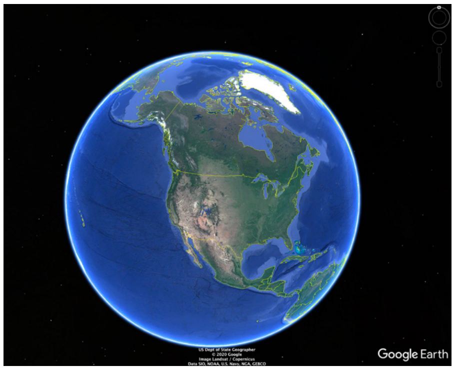

*Penstemon moffatii*

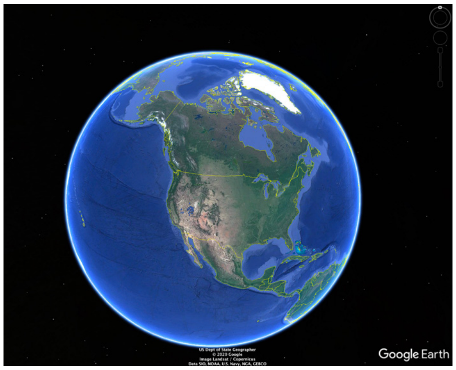

*Penstemon barnebyi*

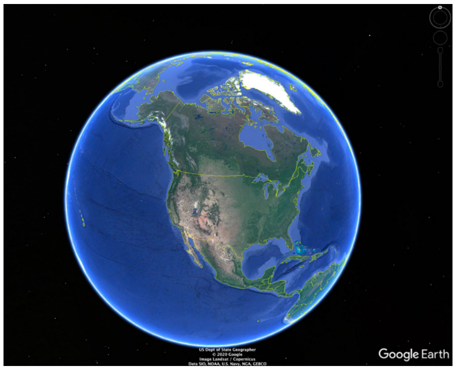

*Penstemon nanus, P. franklinii, P. pinorum, P. concinnus*

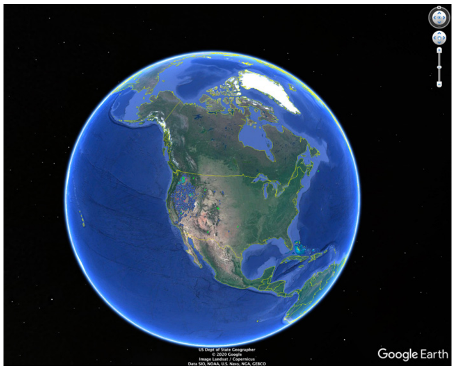

*Penstemon speciosus* clade

#### 0.8 mya to 0.7 mya

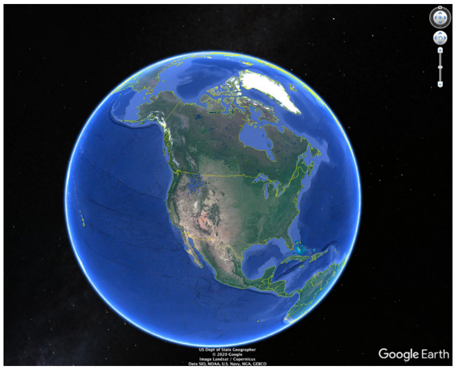

*Penstemon albertinus, P. wilcoxii*

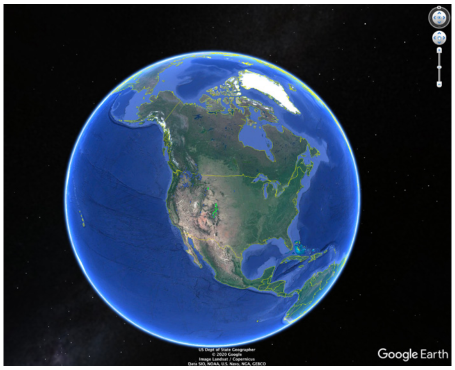

*Penstemon confertus*

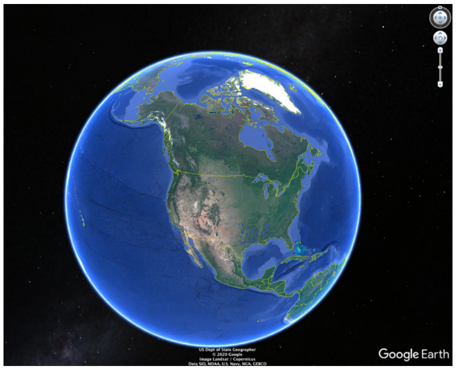

*Penstemon laxiflorus*

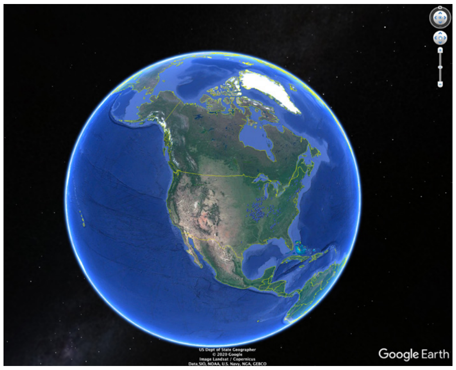

*Penstemon pallidus*

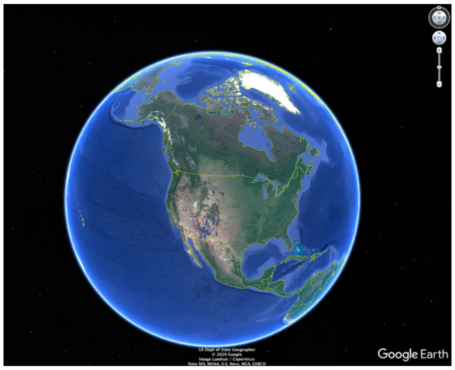

*Penstemon linarioides* vars. *coloradoensis* & *linarioides*

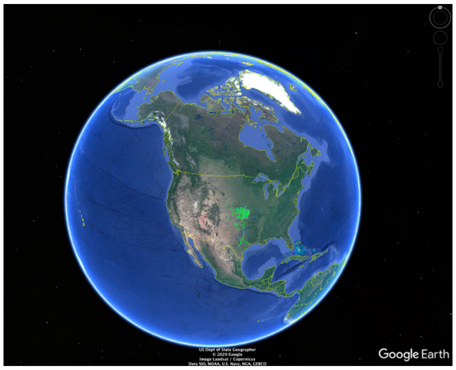

*Penstemon cobaea*

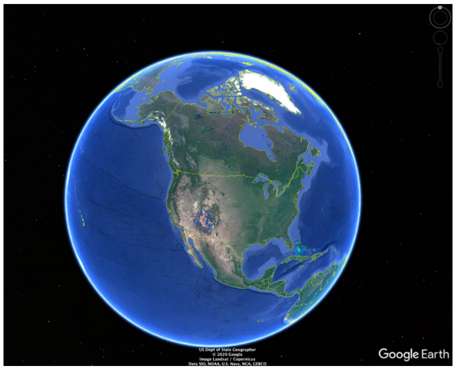

*Penstemon breviculus, P. ophianthus*

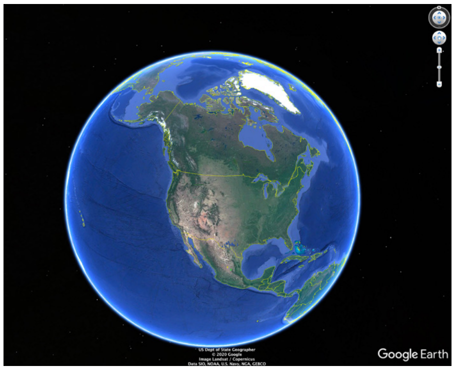

*Penstemon baccharifolius* clade

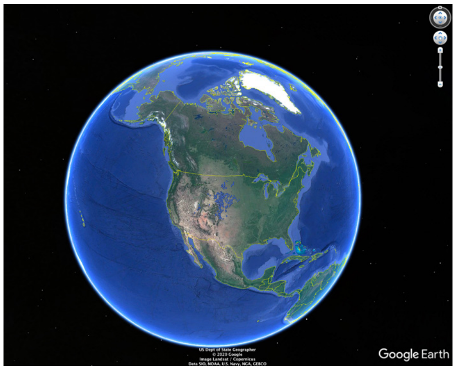

*Penstemon angustifolius* vars. *dulcis* & *angustifolius*

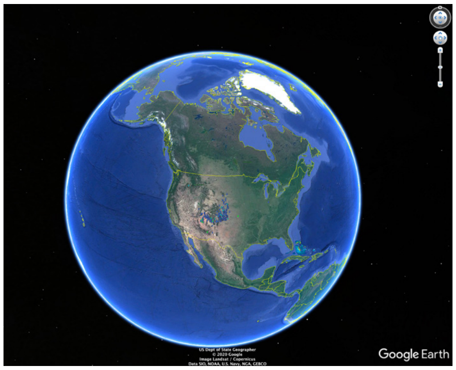

*Penstemon angustifolius* vars. *venosus* & *caudatus*

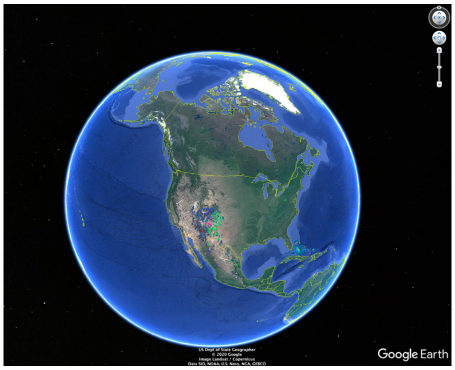

*Penstemon commarhenus* & rest of clade

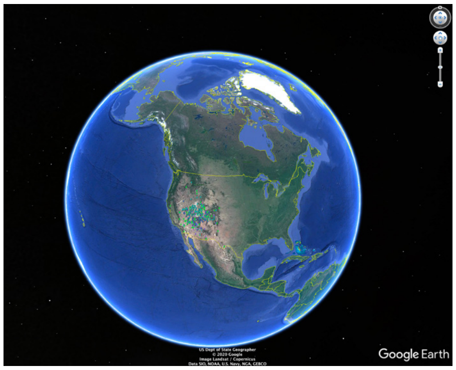

*Penstemon leiophyllus* & rest of clade

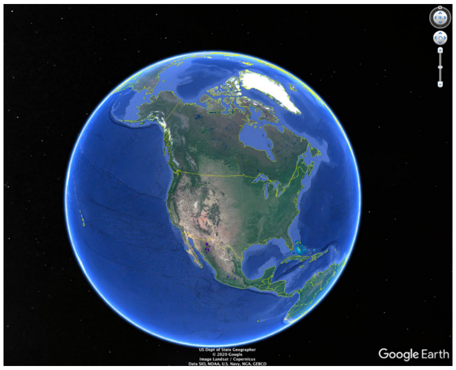

*Penstemon imberbis, P. wislizenii*

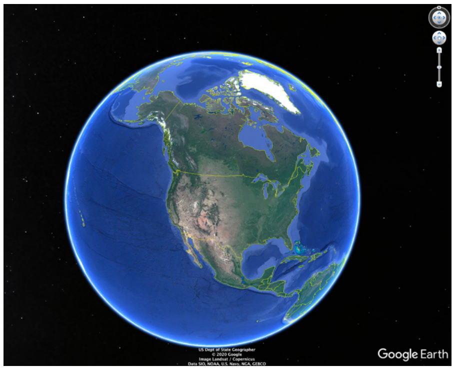

*Penstemon isophyllus*

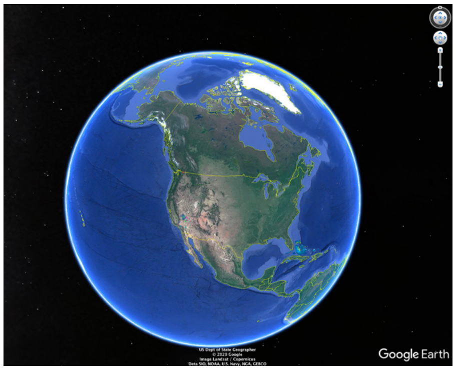

*Penstemon fruticiformis*

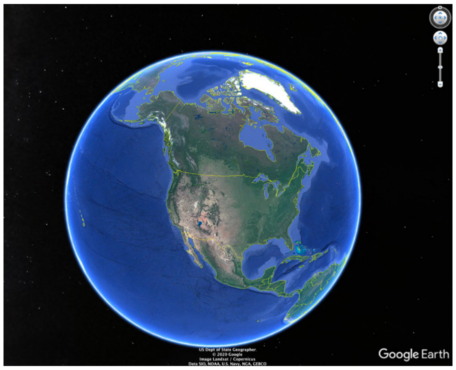

*Penstemon clutei*

### 0.7 mya to 0.6 mya

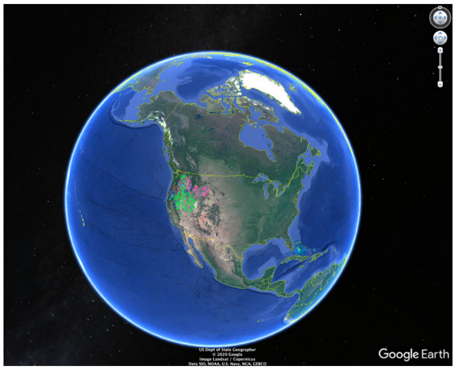

*Penstemon deustus clade*

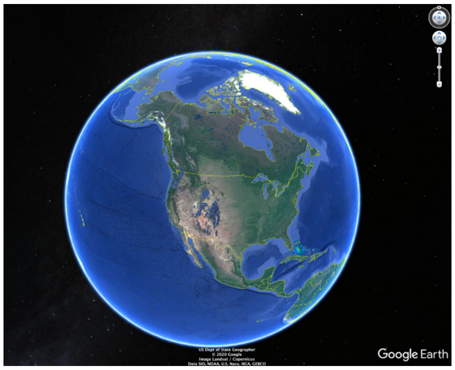

*Penstemon serrulatus, P. whippleanus*

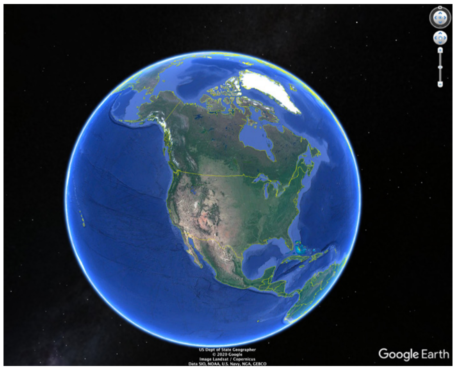

*Penstemon laxus*

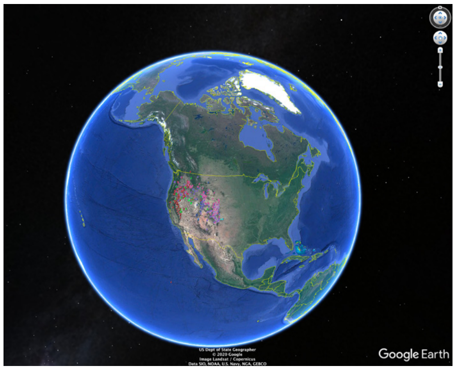

*Penstemon rydbergii*

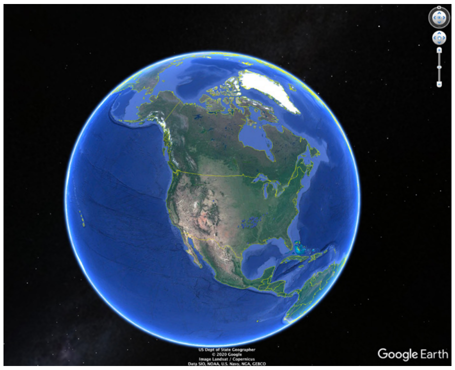

*Penstemon arkansanus, P. oklahomensis*

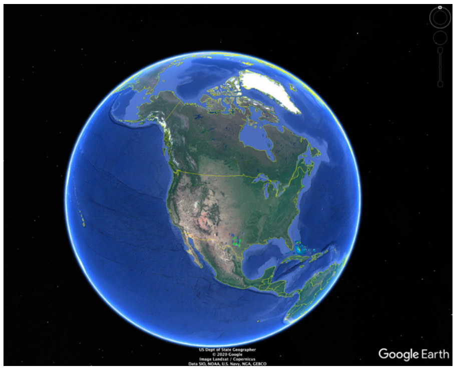

*Penstemon guadalupensis, P. triflorus*

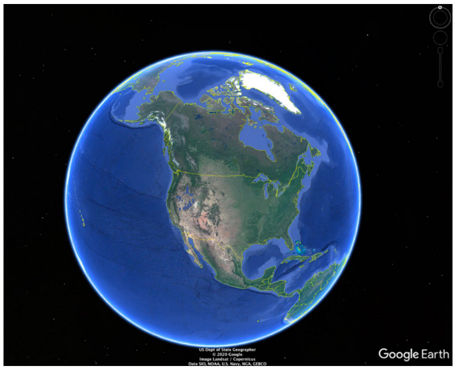

*Penstemon janishiae, P. miser*

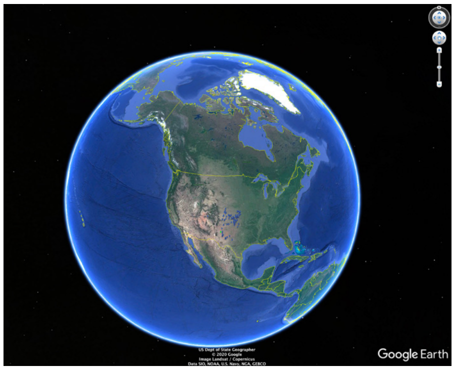

*Penstemon buckleyi*

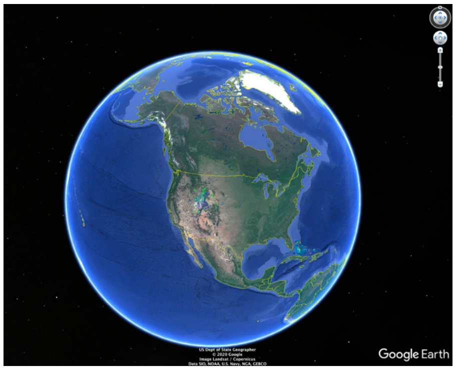

*Penstemon cyananthus*

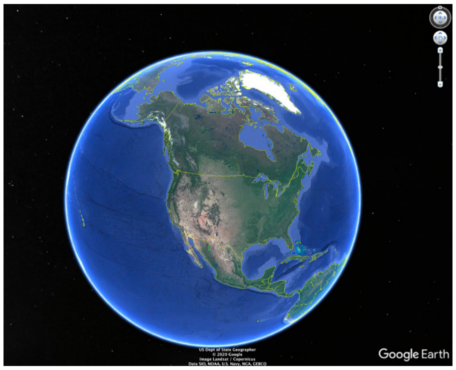

*Penstemon miniatus, P. plagapineus*

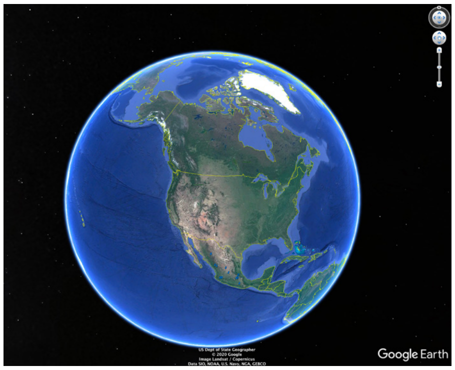

*Penstemon roseus*

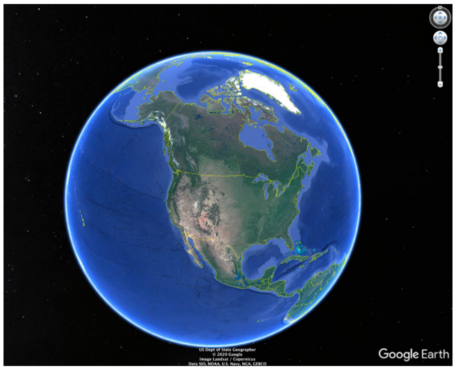

*Penstemon gentianioides, P. hartwegii*

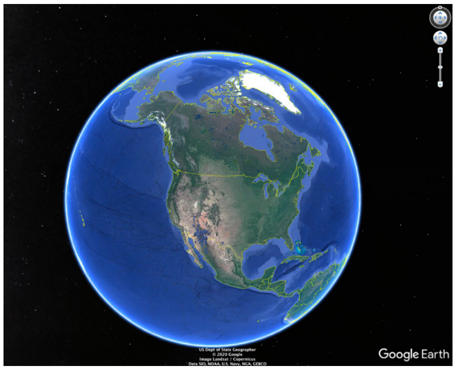

*Penstemon parryi, P. superbus*

#### 0.6 mya to 0.5 mya

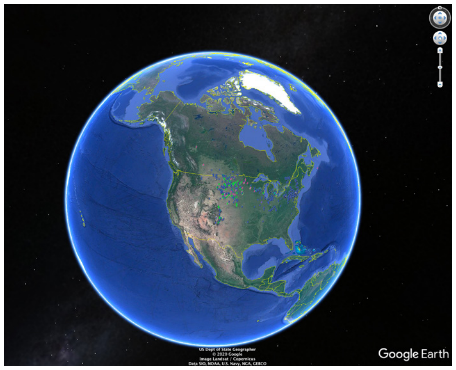

*Penstemon hirsutus, P. gracilis*

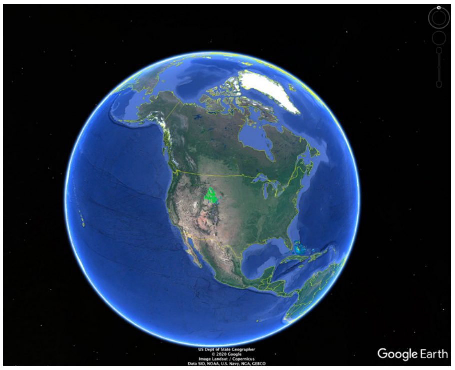

*Penstemon laricifolius*

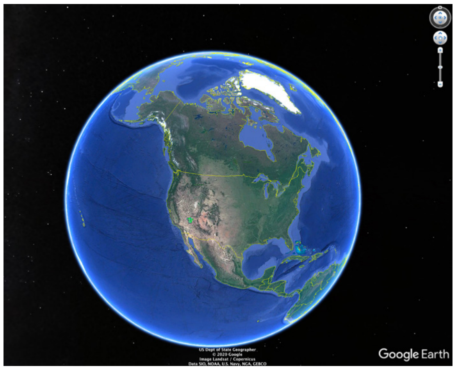

*Penstemon bicolor*

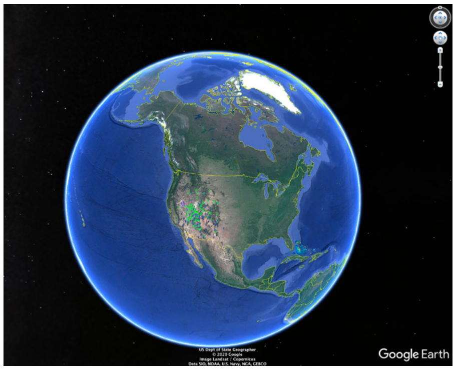

*Penstemon palmeri*

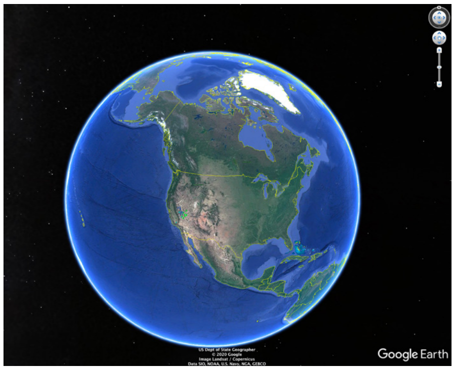

*Penstemon floridus, P. rubicundus*

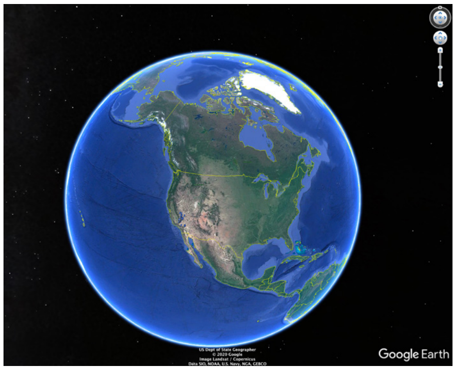

*Penstemon incertus* clade

#### 0.5 mya to 0.4 mya

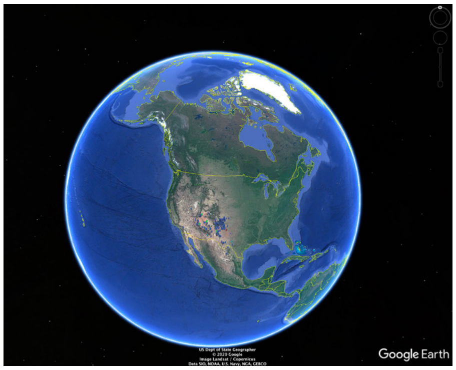

*Penstemon atwoodii, P. jamesii*

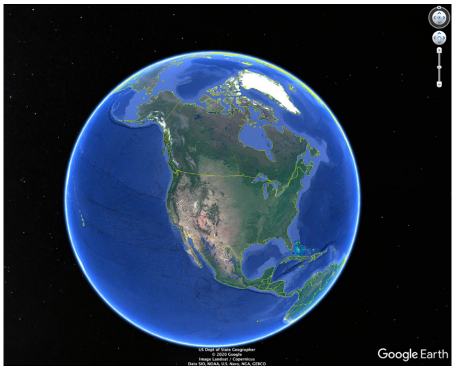

*Penstemon campanulatus*

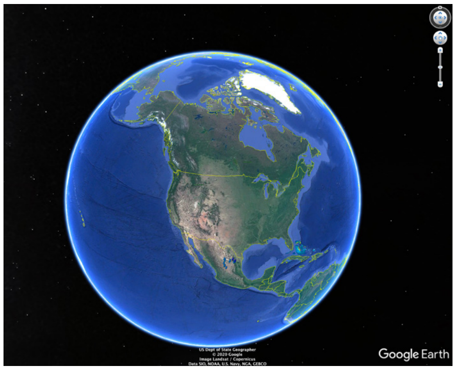

*Penstemon amphorellae* clade

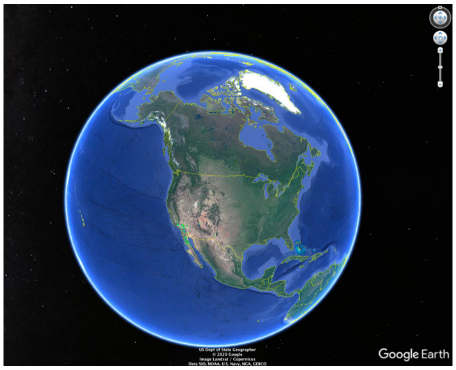

*Penstemon spectabilis*

#### 0.4 mya to 0.3 mya

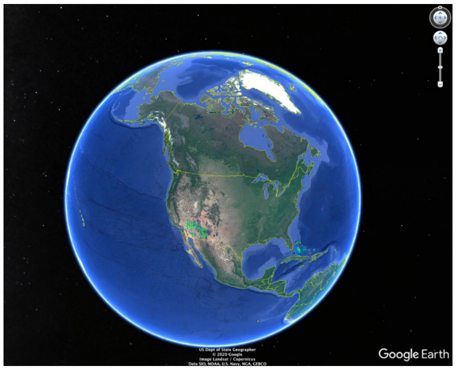

*Penstemon pseudospectabilis*

#### 0.3 mya to 0.2 mya

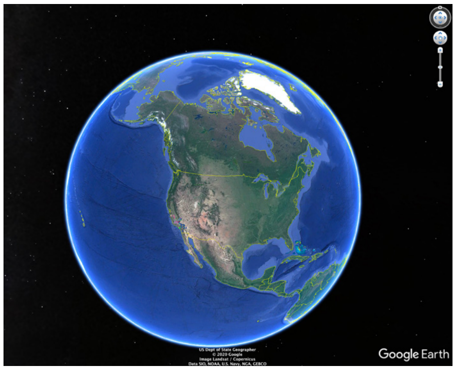

*Penstemon grinnellii*

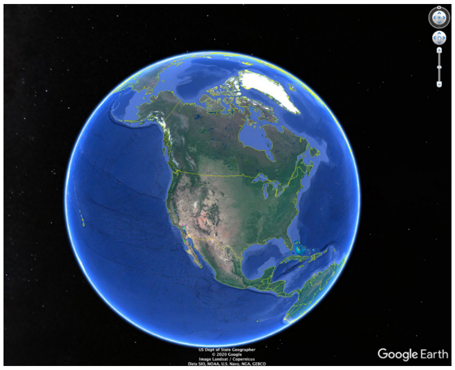

*Penstemon clevelandii*

### 0.2 mya to 0.1 mya

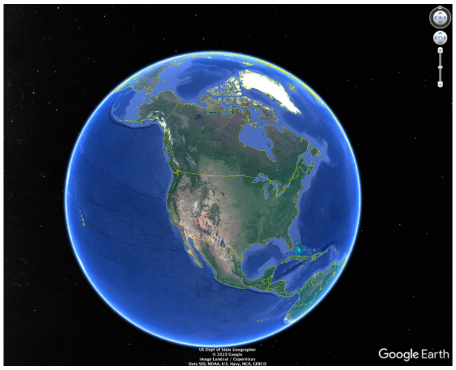


*Penstemon havardii, P. wrightii*
