## Supplementary material for "Phylogenetics of a Rapid, Continental Radiation: Diversification, Biogeography, and Circumscription of the Beardtongues (*Penstemon*; Plantaginaceae)": Table S1

Table S1. Primers used for targeted amplicon sequencing. * denotes primer pairs that were not used in the study.

| Locus | Forward Primer | Reverse Primer |
|---|---|---|
| COS4270 | ACACTGACGACATGGTTCTACAACCAAGCTCTTCACCTGGAA | TACGGTAGCAGAGACTTGGTCTAGACCAGCATAACAATTTTATTCCTAA |
| COS14240 | ACACTGACGACATGGTTCTACACCGACATTAGTCACGGTCCT | TACGGTAGCAGAGACTTGGTCTCGGCATTCCTTCAGATAAAC |
| COS23460 | ACACTGACGACATGGTTCTACATGGTTGTTCGTGTGAGGTTG | TACGGTAGCAGAGACTTGGTCTAAACTCATTATTTGCTCGATAGGG |
| COS24530 | ACACTGACGACATGGTTCTACATTAAAATGCAGGAGGGCTTG | TACGGTAGCAGAGACTTGGTCTCCAACCAAATCAGTTCTCTGC |
| COS50360 | ACACTGACGACATGGTTCTACACCATGGAATCAAACCTGGAC | TACGGTAGCAGAGACTTGGTCTAAGCCCAAATCGAAGAAGAA |
| COS57850 | ACACTGACGACATGGTTCTACAGAAGGAGCCTCAAAGCAGTG | TACGGTAGCAGAGACTTGGTCTGGATGTCCATCTAACCCGTTT |
| PPR876 | ACACTGACGACATGGTTCTACACAGCTTCTGGTAGATGGGCT | TACGGTAGCAGAGACTTGGTCTCTCTCCCCACAATCTTCGCC |
| PPR1651 | ACACTGACGACATGGTTCTACAACGCAGCTTCTGGTAGATGG | TACGGTAGCAGAGACTTGGTCTCACCACAATTACACACGCCC |
| PPR5729 | ACACTGACGACATGGTTCTACATTTGCTCACTGCTTGTGCTG | TACGGTAGCAGAGACTTGGTCTCCTTCATCCACCATGCCACA |
| PPR985 | ACACTGACGACATGGTTCTACATCCGTGGCATTTTTAGTGCG | TACGGTAGCAGAGACTTGGTCTCCGGAGAAAGCTCTTACGGTT |
| PPR1839 | ACACTGACGACATGGTTCTACATGCAACCGTAATGCTCGACT | ACACTGACGACATGGTTCTACATGCAACCGTAATGCTCGACT |
| 34130 | ACACTGACGACATGGTTCTACATCTAAGTTTGCGGATGTTGAGA | TACGGTAGCAGAGACTTGGTCTCATTCCCAGAACATACATGCAA |
| 59820 | ACACTGACGACATGGTTCTACAGCAGATTTAGTTTTACTCTCCTCCA | TACGGTAGCAGAGACTTGGTCTGGTCTTAAATACCATCTTCTGTGTCC |
| 80460 | ACACTGACGACATGGTTCTACACCGAAATTTTACCCAAAATCG | TACGGTAGCAGAGACTTGGTCTGCAATGTGGGATTTGTTCGT |
| 20370 | ACACTGACGACATGGTTCTACATTCAGAGCTCCCATTTTTGC | TACGGTAGCAGAGACTTGGTCTTTGACCTTCATCCAATAGAGCA |
| PPR369 | ACACTGACGACATGGTTCTACAGGAAAGGAAATCCATGCCCA | TACGGTAGCAGAGACTTGGTCTAGCCTTCAGTTACCATTCCG |
| PPR950 | ACACTGACGACATGGTTCTACATCTCCATCTTCGAGAACGCC | TACGGTAGCAGAGACTTGGTCTGCCGGATCAGGTGACGATAG |
| PPR1250 | ACACTGACGACATGGTTCTACAAAAGCCCTTCTTGCACGAGT | TACGGTAGCAGAGACTTGGTCTGGACAGCTTTGATTGCAGGG |
| PPR1561 | ACACTGACGACATGGTTCTACATCCCTTTTGCCTCATCGACC | TACGGTAGCAGAGACTTGGTCTGGTTCACACGGTGAATGTCG |
| 30360 | ACACTGACGACATGGTTCTACAAGGTTGCTAAAGGCCGATTC | TACGGTAGCAGAGACTTGGTCTGGGTCTTTATCTAAAAGGCGAGA |
| 35920 | ACACTGACGACATGGTTCTACAGGGGACAAAAATAGCAGAGC | TACGGTAGCAGAGACTTGGTCTTCACCGTGCTTGTTAAGTGC |
| 27260 | ACACTGACGACATGGTTCTACACTCCCCCGGAAAGTAACAAA | TACGGTAGCAGAGACTTGGTCTTTGTTTCATGTTGCGCCTTT |
| 2840 | ACACTGACGACATGGTTCTACATCTGGAAAAATTCCCTGGAC | TACGGTAGCAGAGACTTGGTCTGCGCTTTGCAAATTCTTGAG |
| 53950 | ACACTGACGACATGGTTCTACAAAACTGTGCTCTTCCTCCAA | TACGGTAGCAGAGACTTGGTCTGGGGAATGGTGACTCCTACA |

| Locus | Forward Primer | Reverse Primer |
|---|---|---|
| 62010 | ACACTGACGACATGGTTCTACAAGCAACGCCATAAACTGGAA | ACACTGACGACATGGTTCTACAAGCAACGCCATAAACTGGAA |
| 2350 | ACACTGACGACATGGTTCTACAGCTCCCATCTTTGTATATCTCG | TACGGTAGCAGAGACTTGGTCTCCCGTTCGTTCGATTGATAG |
| 18520 | ACACTGACGACATGGTTCTACATTCGAGAACGCCTCTAAACC | TACGGTAGCAGAGACTTGGTCTGTTATGCACAAAACGGGATG |
| 33495 | ACACTGACGACATGGTTCTACATCTCCATTTCTCAACCTCAGC | TACGGTAGCAGAGACTTGGTCTGCCCCTTCCTCTCCATACA |
| 38180 | ACACTGACGACATGGTTCTACAGGCATCAAAAGTGGATGATG | TACGGTAGCAGAGACTTGGTCTTCCCTCGTTGAGACATTCCT |
| 4810 | ACACTGACGACATGGTTCTACATACCAATTCCCCAGTTCTGC | TACGGTAGCAGAGACTTGGTCTATGGGAAGAGATGCTTACCTGA |
| 37450 | ACACTGACGACATGGTTCTACATGCTATCAAAACTTCGGCATC | TACGGTAGCAGAGACTTGGTCTGATCTCAAAAAGCACAACTCCA |
| 66330 | ACACTGACGACATGGTTCTACATGCAAATTCTTGAGCTGTCC | TACGGTAGCAGAGACTTGGTCTAAAATTCCCTGGACGCTTG |
| 48730 | ACACTGACGACATGGTTCTACACCATACGCGTAAATAAGAGAGC | TACGGTAGCAGAGACTTGGTCTTGATGGATATGGTAAAGCTAAACG |
| 3382 | ACACTGACGACATGGTTCTACATCTGAAAGCCTTGTACCAACC | TACGGTAGCAGAGACTTGGTCTGAGCCCTCTTGCCATTTCTA |
| 2829 | ACACTGACGACATGGTTCTACATGACCCGTTGACAACCCTAT | ACACTGACGACATGGTTCTACATGACCCGTTGACAACCCTAT |
| 604 | ACACTGACGACATGGTTCTACAATGGCCTCCGGTAATTCTCT | TACGGTAGCAGAGACTTGGTCTAGTGGCCTGAACTTTGCAGT |
| 598 | ACACTGACGACATGGTTCTACATCTGGGCTAACCTGAAATCG | TACGGTAGCAGAGACTTGGTCTTGGAAATGATCAAGAAATGAAGC |
| 233 | ACACTGACGACATGGTTCTACAACAACGCTGTGTGTTTGGTC | TACGGTAGCAGAGACTTGGTCTCCCACCAGCCCTTAACTACTC |
| 2978 | ACACTGACGACATGGTTCTACATCCATACAATGGTAAGATCACAAGA | TACGGTAGCAGAGACTTGGTCTGAAGGGTGTTCCGGGATTAT |
| 2919 | ACACTGACGACATGGTTCTACAAGAGTGTCATGGCCACCAAT | TACGGTAGCAGAGACTTGGTCTATGGACCGCATAGCTCAAAG |
| 2782 | ACACTGACGACATGGTTCTACAGAATTGAGGAGATTTGGGAATTT | TACGGTAGCAGAGACTTGGTCTCAGAATTGGGCCCTCCTAAG |
| 1397 | ACACTGACGACATGGTTCTACATCCAGTTTCGCTGAAATCACT | TACGGTAGCAGAGACTTGGTCTTAAAGGCCTTGGAGAAGCAA |
| 836 | ACACTGACGACATGGTTCTACATCCCAATTTATCCCAGAAAGC | TACGGTAGCAGAGACTTGGTCTAATCATGGGCGACCTATTTG |
| 21370* | ACACTGACGACATGGTTCTACACTGTTTTTCCAATTTTCCATCC | TACGGTAGCAGAGACTTGGTCTCAGGTTGTGGGCTACGATTT |
| 1331* | ACACTGACGACATGGTTCTACAGCACCAAGATATGCCATTGA | TACGGTAGCAGAGACTTGGTCTTCCGAGCTAAGGCTATACATTCA |
| rps12rpl20* | ACACTGACGACATGGTTCTACAATTAGAAANRCAAGACAGCCAAT | TACGGTAGCAGAGACTTGGTCTCGYYAYCGAGCTATATATCC |
| trnTL* | ACACTGACGACATGGTTCTACACATTACAAATGCGATGCTCT | TACGGTAGCAGAGACTTGGTCTTCTACCGATTTCGCCATATC |
| trnCD* | ACACTGACGACATGGTTCTACACCAGTTCAAATCTGGGTGTC | TACGGTAGCAGAGACTTGGTCTGGGATTGTAGTTCAATTGGT |
