## Supplementary material for "Phylogenetics of a Rapid, Continental Radiation: Diversification, Biogeography, and Circumscription of the Beardtongues (*Penstemon*; Plantaginaceae)": Table S2

Supplement Table S2. Biogeographic chronology based on time-calibrated tree.

| Time before present | Number of Glaciation Cycles | Species of *Penstemon* existing in time frame in chronological order (geographic area shown in Fig. ##) | Distribution Notes |
|---|---|---|---|
| 2.7 mya–2 mya | 4 major, 1 minor | *montanus* (6)<br>*lyallii* (6)<br>*ellipticus* (6)<br>*fruticosus* (3,6)<br>*davidsonii* (2,3)<br>*rupicola* (3)<br>*barrettiae* (3)<br>*cardwellii* (3)<br>*newberryi* (3)<br>*personatus* (3) | Eastern cordillera (6) from Colorado Plateau north to northwest cordillera, south throughout Cascade-Sierra (3) and Pacific Coast ranges (3), and lower elevations of Pacific Northwest (2). |
| 2 mya–1.8 mya | 1 major | *venustus* (2)<br>*dissectus* (10)<br>*calcareus* (11) | First appearance of *Penstemon* in the eastern United States (Coastal Plains (10): *P. dissectus*); *P. venustus* is in the Pacific Northwest (2); *P. calcareus* is in the southwest cordillera region (11). |
| 1.8 mya –1.7 mya | 2 minor | *smallii* (9)<br>*sudans* (3) | *P. sudans* is in the Cascade-Sierra range (3). First appearance of *Penstemon* in the Appalachian region (9). |
| 1.7 mya–1.6 mya | none | *harbourii* (6)<br>*retorsus* (6) | Southern area of eastern cordillera (6). |
| 1.6–1.5 mya | 2 minor | *glandulosus* (2)<br>*aridis* (6)<br>*anguineus* (3) | Eastern cordillera (6) and Cascade-Sierra ranges (3) |

| | | | |
|---|---|---|---|
| | | | and Pacific Northwest (2) |
| 1.5–1.4 mya | none | *radicosus* (5,**6**) | Eastern cordillera (6) and eastern intermountain region (5) |
| 1.4 mya–1.3 mya | none | *virens* (6)<br>*pruinosus* (2)<br>*humilis obtusifolius* (5)<br>*humilis breviolius* (5,**6**)<br>*humilis humilis* (2,5,6)<br>*griffinii* (6)<br>*inflatus* (6)<br>*degeneri* (6)<br>*pumilis* (6)<br>*carnosus* (5)<br>*albomarginatus* (5)<br>*ambiguus* (5,6.7,11)<br>*stenophyllus* (11,12) | Eastern cordillera (6) to Pacific Northwest (2), then south and east through intermountain region (5) into the western Great Plains (7).<br>Eastern cordillera (6) from southern region to northern region, then into the intermountain region (5) and southern Great Plains (7), and into the northwestern highlands of Mexico (11,12). |
| 1.3 mya–1.2 mya | none | *laetus laetus* (3)<br>*laetus sagittatus* (3)<br>*parvulus* (3)<br>*sepalulus* (5)<br>*attenuatus attenuatus* (3,**6**)<br>*elegantulus* (6)<br>*watsonii* (5,6)<br>*rydbergii rydbergii* (2,**6**)<br>*rydbergii oreocharis* (2,**3**)<br>*monoensis* (5)<br>*confusus* (5)<br>*patens* (5)<br>*ammophilus* (5)<br>*flowersii* (5)<br>*harringtonii* (6)<br>*cardinalis* (6) | Southern Cascade-Sierra (3), intermountain region (5), and eastern cordillera (6), then north in both ranges.<br>South intermountain region (5) and southeastern cordillera (6), north into Pacific northwest (2).<br>Intermountain region diversification.<br>Continuation of |

| | | | |
|---|---|---|---|
| | | | diversification in eastern cordillera. |
| 1.2 mya–1.1 mya | 2 major | *pappilatus* (3)<br>*pudicus* (4)<br>*triphyllus* (2)<br>*richardsonii* (2)<br>*diphyllus* (6)<br>*subserratus* (3)<br>*procerus procerus* (5,**6**)<br>*canescens* (9)<br>*teucrioides* (6)<br>*procumbens* (6)<br>*tusharensis* (5)<br>*pinifolius* (11)<br>*duchesnensis* (5)<br>*acaulis* (6)<br>*yampaensis* (6)<br>*gormanii* (1)<br>*dasyphyllus* (7,11)<br>*lanceolatus* (11,12)<br>*nudiflorus* (11)<br>*fendleri* (7,11)<br>*versicolor* (**6**,7)<br>*lavendulus* (6)<br>*cyathophorus* (6)<br>*mensarum* (6) | Cascade-Sierra (3,4) then up through the Pacific Northwest to the eastern cordillera (6). Oscillations east and west. Southeastern cordillera (6) to southwest (11), and then return to middle latitudes of eastern cordillera (6). In the east, distribution through the Appalachian region expands (9). Expansion from Cascade-Sierra (3) into Yukon Territory and Alaska (1). Expansion from southeastern cordillera (6) into southwest (11) and Mexico highlands (12). Expansion from southeastern cordillera (6) into southern Great Plains (7). |
| 1.1 mya –1.0 mya | 1 minor | *cusickii* (2)<br>*flavescens* (6)<br>*spatulatus* (2)<br>*auriberbis* (7)<br>*eriantherus eriantherus* (6,7)<br>*eriantherus cleburnei* (6)<br>*goodrichii* (5)<br>*immanifestus* (5) | Pacific Northwest (2) then east through eastern cordillera (6) and Great Plains (7). Eastern intermountain region (5) |

| | | | |
|---|---|---|---|
| | | *bracteatus* (5)<br>*pachyphyllus pachyphyllus* (5)<br>*pachyphyllus congestus* (5)<br>*secundiflorus* (**6**,7) | expanding west, and east into Great Plains (7). |
| 1.0 mya–0.9 mya | 1 minor | *rostriflorus* (5)<br>*platyphyllus* (5)<br>*patricus* (5)<br>*attenuatus pseudoprocerus* (6)<br>*ovatus* (3)<br>*attenuatus militaris* (6)<br>*euglaucus* (3)<br>*tubaeflorus* (**7**,8)<br>*multiflorus* (10)<br>*tenuis* (10)<br>*crandallii crandallii* (6)<br>*crandallii atratus* (5)<br>*ramaleyi* (6)<br>*glabrescens* (5)<br>*abietinus* (5)<br>*caespitosus caespitosus* (5,6)<br>*caespitosus perbrevis* (5)<br>*caespitosus desertipicti* (5,11)<br>*lentus* (5)<br>*strictiformis* (5,**6**)<br>*compactus* (5)<br>*luculentus* (6)<br>*gibbensii* (6)<br>*debilis* (6)<br>*caryi* (6)<br>*utahensis* (**5**,6)<br>*penlandii* (6)<br>*scariosus cyanomontanus* (5)<br>*scariosus albifluvis* (5)<br>*scariosus garrettii* (5)<br>*pseudoputus* (5,**11**)<br>*labrosus* (4)<br>*deaveri* (11)<br>*virgatus* (6,11)<br>*glaber* (6) | Southern Cascade-Sierra (3) and intermountain region (5), expanding north to eastern cordillera (6) and northern Cascade-Sierra (3). In the east: expansion from Appalachian region (9) into southern interior lowlands and eastern Great Plains (7,8). Southeastern cordillera (6) expanding into intermountain region (5), then southwest (11) and back into eastern cordillera (6). In the east, expansion through the coastal plain (10) |
| 0.9 mya–0.8 mya | 1 major | *caesius* (3)<br>*scapioides* (5)<br>*azureus* (3)<br>*heterophyllus* (3,4)<br>*neotericus* (3) | Southwest Cascade-Sierra (3) and spread through southwest |

| | | | |
|---|---|---|---|
| | | *kingii* (5)<br>*leonardii* (5)<br>*heterodoxus* (3)<br>*attenuatus palustris* (2)<br>*pratensis* (2,5)<br>*hesperius* (2)<br>*washingtonensis* (3)<br>*thompsoniae* (5)<br>*linarioides sileri* (5,**11**)<br>*californicus* (4)<br>*centranthifolius* (4)<br>*cerrosensis* (4)<br>*subulatus* (11)<br>*mucronatus* (6)<br>*pachyphyllus pachyphyllus* (5)<br>*arenicola* (6)<br>*angustifolius vernalensis* (5)<br>*nitidus* (6,**7**)<br>*grandiflorus* (**7**,8)<br>*cyanocaulis* (**5**,6)<br>*mensarum* (5)<br>*strictus* (6)<br>*hallii* (6)<br>*parvus* (5)<br>*longiflorus* (5)<br>*idahoensis* (5)<br>*uintahensis* (5)<br>*fasciculatus* (12)<br>*hidalgensis* (12)<br>*coriaceus* (12) | cordillera (4). Intermountain region diversification (5) and extension into southwest region (11) and southwest cordillera (4). Southeast cordillera (6) into southern intermountain region (5), southwestern cordillera (4) into northern Baja California (4) followed by trajectory north again. Eastern cordillera (6) into northern Great Plains (7), then east (8). Eastern cordillera (6) then west into intermountain region (5). Diversification in Mexico highlands (12). |
| 0.8 mya–0.7 mya | none | *laetus sagittatus* (3)<br>*filiformus* (3)<br>*purpusii* (4)<br>*roezlii* (3)<br>*gracilentus* (3)<br>*globosus* (6)<br>*peckii* (3)<br>*cinicola* (3)<br>*digitalis* (**8**,9)<br>*calycosus* (**8**,9)<br>*moffatii* (5)<br>*barnebyi* (5)<br>*nanus* (5)<br>*franklinii* (5) | Cascade-Sierra (3) to southwestern cordillera (4) into intermountain region (5), eastern cordillera (6) and into the Great Plains (7). In the east, expansion in interior lowland (8) and Appalachian region (9). |

| | | | |
|---|---|---|---|
| | | *pinorum* (5)<br>*concinnus* (5)<br>*speciosus* (2,5,7)<br>*payettensis* (6)<br>*acuminatus acuminatus* (2) | |
| 0.7 mya–0.6 mya | 1 major | *albertinus* (6)<br>*wilcoxii* (6)<br>*confertus* (2,6)<br>*laxiflorus* (10)<br>*pallidus* (8)<br>*linarioides coloradoensis* (6,**11**)<br>*linarioides linarioides* (6,**11**)<br>*cobaea* (7)<br>*breviculus* (5,11)<br>*ophianthus* (5,**11**)<br>*baccharifolius* (7)<br>*wendtiorum* (7)<br>*occiduus* (12)<br>*angustifolius dulcis* (5)<br>*angustifolius angustifolius* (6,**7**)<br>*angustifolius venosus* (**5**,11)<br>*angustifolius caudatus* (6,**7**)<br>*virgatus virgatus* (6,11)<br>*commarrhenus* (5,6)<br>*barbatus* (5,6,11,12)<br>*neomexicanus* (11)<br>*moriahensis* (5)<br>*virgatus asa-grayii* (6)<br>*leiophyllus leiophyllus* (5)<br>*eatonii undosus* (5,11)<br>*eatonii eatonii* (5,**6**)<br>*eatonii exertus* (11)<br>X *jonesii* (5)<br>*laevis* (5)<br>*pahutensis* (5)<br>*imberbis* (12)<br>*wislizeneii* (12)<br>*isophyllus* (12)<br>*fruticiformis* (4,5)<br>*clutei* (11) | Eastern cordillera (6), north through Pacific Northwest (2), south into intermountain region (5) and southwest (11). Expansion east into the Great Plains (7), southwest (11), and Mexico highlands (12). In the east, diversification in interior lowland (8) and coastal plain (10). |
| 0.6 mya–0.5 mya | none | *deustus deustus* (2)<br>*deustus suffrutescens* (3)<br>*gairdneri oreganus* (2) | Pacific Northwest (2), Cascade-Sierra (3), eastern |

| | | | |
|---|---|---|---|
| | | *gairdneri gairdneri* (2)<br>*seorsus* (2)<br>*serrulatus* (2)<br>*whippleanus* (6)<br>*laxus*(6)<br>*rydbergii oreocharis* (2,**3**)<br>*arkansanus* (8)<br>*oklahomensis* (7)<br>*guadalupensis* (7)<br>*triflorus integrifolius* (7)<br>*janishiae* (5)<br>*miser* (2,6)<br>*buckleyi* (7)<br>*cyananthus cyananthus* (5,6)<br>*cyananthus subglaber* (5,**6**)<br>*miniatus* (12)<br>*plagapineus* (12)<br>*roseus* (12)<br>*gentianioides* (12)<br>*hartwegii* (12)<br>*parryi* (12)<br>*superbus* (11) | cordillera (6). Diversification into Great Plains (7), interior lowland (8), Intermountain region (5), and Mexico Highlands (12). |
| 0.5 mya–0.4 mya | 1 major | *hirsutus* (8,**9**)<br>*gracilis* (7)<br>*laricifolius* (6)<br>*bicolor* (5)<br>*palmeri palmeri* (5)<br>*palmeri eglandulosus* (5,**6**)<br>*floridus* (5)<br>*rubicundus* (5)<br>*incertus* (5)<br>*angelicus* (4)<br>*vizcainensis* (4)<br>*reidmoranii* (4) | Great Plains (7), Eastern Cordillera (6), Intermountain Region (5), and Southwest Cordillera (5). In east, Interior lowlands and Appalachian region (8,9). |
| 0.4 mya–0.3 mya | 1 minor | *atwoodii* (5)<br>*jamesii* (7,**11**)<br>*campanulatus campanulatus* (12)<br>*campanulatus chihuahuensis* (11,12)<br>*leonensis* (12)<br>*vulcanellus* (12)<br>*tenuifolius* (12)<br>*potosinus* (12)<br>*amphorellae* (12)<br>*spectabilis spectabilis* (4)<br>*spectabilis subviscosus* (4) | Mostly southern expansions: Intermountain region (5), Great Plains (7) into Southwest (11). Southwestern cordillera (4), and Mexico highlands (12). |

| | | | |
|---|---|---|---|
| | | X *parishii* (4) | |
| 0.3 mya–0.2 mya | 1 major | *pseudospectabilis* (11) | Southwest (11) |
| 0.2 mya–0.1 mya | 1 major | *clevelandii clevelandii* (4)<br>*clevelandii connatus* (4)<br>*clevelandii mojavensis* (4)<br>*grinnellii grinnellii* (4)<br>*grinnellii scrophularioides* (4) | Southwest cordillera (4) |
| 0.1 mya–present | 1 major<br>1 minor | *havardii* (7)<br>*wrightii* (5,7) | Southern Great Plains (7) and Intermountain region (5). |
