## Supplementary material for "Phylogenetics of a Rapid, Continental Radiation: Diversification, Biogeography, and Circumscription of the Beardtongues (*Penstemon*; Plantaginaceae)": Table S3

Table S3. Revised Taxonomy of *Penstemon* based on phylogenetics. Subgenera and sections appear in same order as in Figure 1; species are listed in alphabetical order. Sections with subsections are indicated by *; subsections listed last. Taxa not in Fig. 1 are placed based on morphology and previous classification (Lindgren & Wilde 2003).

**Subgenus *Dasanthera***

*P. barrettiae*
*P. cardwellii*
*P. davidsonii*
*P. ellipticus*
*P. fruticosus*
*P. lyallii*
*P. montanus*
*P. newberryi*
*P. rupicola*

**Subgenus *Cryptostemon***

*P. personatus*

**Subgenus *Saccanthera***

*P. anguineus*
*P. azureus*
*P. caesius*
*P. cusickii*
*P. deustus*
*P. filiformis*
*P. floribundus*
*P. gairdneri*
*P. glandulosus*
*P. gracilentus*
*P. heterophyllus*
*P. kingii*
*P. laetus*
*P. leonardii*
*P. neotericus*
*P. papillatus*
*P. parvulus*
*P. patricus*
*P. platyphyllus*
*P. pudicus*
*P. purpursii*
*P. rhizomatosus*
*P. richardsonii*
*P. roezlii*
*P. rostriflorus*
*P. scapoides*
*P. seorsus*
*P. sepalulus*
*P. serrulatus*
*P. sudans*
*P. tiehmii*
*P. triphyllus*
*P. venustus*
*P. whippleanus*

**Subgenus *Penstemon***

<u>Section *Proceri**</u>

*P. albertinus*
*P. aridus*
*P. attenuatus*
*P. bleaklyi*
*P. cinicola*
*P. confertus*
*P. diphyllus*
*P. euglaucus*
*P. degeneri*
*P. elegantulus*
*P. flavescens*
*P. glaucinus*
*P. globosus*
*P. griffinii*
*P. hesperius*
*P. heterodoxus*
*P. humilis*
*P. inflatus*
*P. laxus*
*P. metcalfei*
*P. oliganthus*
*P. ovatus*
*P. peckii*
*P. pratensis*
*P. procerus*
*P. pruinosus*
*P. pseudoparvus*
*P. radicosus*
*P. rattanii*
*P. rydbergii*
*P. spatulatus*
*P. subserratus*
*P. tracyi*
*P. virens*
*P. washingtonensis*
*P. watsonii*
*P. wilcoxii*

<u>Section *Penstemon*</u>

*P. arkansanus*
*P. australis*
*P. brevisepalis*
*P. calycosus*
*P. canescens*
*P. deamii*
*P. digitalis*
*P. gracilis*
*P. hirsutus*
*P. kralii*
*P. laevigatus*
*P. laxiflorus*
*P. multiflorus*
*P. oklahomensis*
*P. pallidus*
*P. smallii*
*P. tenuiflorus*
*P. tenuis*
*P. tubaeflorus*

<u>Section *Caespitosi**</u>

*P. abietinus*
*P. caespitosus*
*P. californicus*
*P. crandallii*
*P. discolor*
*P. glabrescens*
*P. harbourii*
*P. linarioides*
*P. pinifolius*
*P. procumbens*
*P. ramaleyi*
*P. retrorsus*
*P. teucrioides*
*P. thompsoniae*
*P. tusharensis*

<u>Section *Dissecti*</u>

*P. dissectus*

Section *Cristati*

*P. acaulis*
*P. albidus*
*P. atwoodii*
*P. auriberbis*
*P. barnebyi*
*P. breviculus*
*P. calcareus*
*P. cerrosensis*
*P. cobaea*
*P. concinnus*
*P. distans*
*P. dolius*
*P. duchesnensis*
*P. eriantherus*
*P. franklinii*
*P. goodrichii*
*P. gormanii*
*P. grahamii*
*P. guadalupensis*
*P. jamesii*
*P. janishiae*
*P. laricifolius*
*P. marcusii*
*P. miser*
*P. moffatii*
*P. monoensis*
*P. nanus*
*P. ophianthus*
*P. pinorum*
*P. pumilis*
*P. triflorus*
*P. yampaensis*

Section *Chamaeleon*

*P. dasyphyllus*
*P. lanceolatus*
*P. ramosus*

Section *Baccharifolii*

*P. baccharifolius*
*P. occiduus*
*P. wendtiorum*

Section *Coerulei*

*P. ammophilus*
*P. angustifolius*
*P. arenicola*
*P. bracteatus*
*P. buckleyi*
*P. carnosus*
*P. centranthifolius*
*P. confusus*
*P. cyathoporus*
*P. fendleri*
*P. flowersii*
*P. grandiflorus*
*P. harringtonii*
*P. haydenii*
*P. immanifestus*
*P. lavendulus*
*P. lentus*
*P. mucronatus*
*P. nitidus*
*P. nudiflorus*
*P. patens*
*P. osterhoutii*
*P. pachyphyllus*
*P. secundiflorus*
*P. subulatus*
*P. versicolor*

Section *Ambiguii*

*P. albomarginatus*
*P. ambiguous*
*P. thurberi*

Section *Habroanthus*

*P. absarokensis*
*P. accuminatus*
*P. barbatus*
*P. cardinalis*
*P. caryi*
*P. commarrhenus*
*P. compactus*
*P. cyananthus*
*P. cyaneus*
*P. cyanocaulis*
*P. deaveri*
*P. debilis*
*P. eatonii*
*P. fremontii*
*P. gibbensii*
*P. glaber*
*P. hallii*
*P. henricksonii*
*P. idahoensis*
*P. labrosus*
*P. laevis*
*P. leiophyllus*
*P. lemhiensis*
*P. longiflorus*
*P. luculentus*
*P. mensarum*
*P. moriahensis*
*P. navajoa*
*P. neomexicanus*
*P. nudiflorus*
*P. pahutensis*
*P. parvus*
*P. payettensis*
*P. paysoniorum*
*P. penlandii*
*P. pennellianus*
*P. perpulcher*
*P. pseudoputus*
*P. putus*
*P. saxosorum*
*P. scariosus*
*P. speciosus*
*P. strictiformis*
*P. strictus*
*P. subglaber*
*P. tidestromii*
*P. uintahensis*
*P. utahensis*
*P. virgatus*
*P. wardii*

Section *Fasciculus*

*P. amphorellae*
*P. bolianus*
*P. campanulatus*
*P. coriaceus*
*P. fasciculatus*
*P. filisepalis*
*P. gentianoides*
*P. gentryi*
*P. hartwegii*
*P. hidalgensis*
*P. hintonii*
*P. imberbis*
*P. isophyllus*
*P. leonensis*
*P. minatus*
*P. mohinoranus*
*P. moronensis*
*P. perfoliatus*
*P. plagapineus*
*P. potosinus*

*P. roseus*
*P. salterius*
*P. skutchii*
*P. stenophyllus*
*P. tenuifolius*
*P. tepicensis*
*P. vulcanellus*
*P. wislizenii*

Section *Spectabiles*

*P. alamosensis*
*P. angelicus*
*P. bicolor*
*P. clevelandii*
*P. clutei*
*P. eximeus*
*P. floridus*
*P. fruticiformis*
*P. grinnellii*
*P. havardii*
*P. incertus*
*P. palmeri*
*P. parryi*
*P. petiolatus*
*P. pseudospectabilis*
*P. reidmoranii*
*P. rubicundus*
*P. rotundifolius*
*P. spectabilis*
*P. stephensii*
*P. superbus*
*P. vizcainensis*
*P. wrightii*

**Subsection designations**

**Subgenus *Penstemon***

Sect. *Proceri* subsect. *Proceri*

- *P. attenuatus*
- *P. cinicola*
- *P. confertus*
- *P. euglaucus*
- *P. flavescens*
- *P. glaucinus*
- *P. globosus*
- *P. hesperius*
- *P. heterodoxus*
- *P. laxus*
- *P. peckii*
- *P. pratensis*
- *P. procerus*
- *P. rydbergii*
- *P. spatulatus*
- *P. tracyi*
- *P. washingtonensis*

Sect. *Proceri* subsect. *Humiles*

- *P. albertinus*
- *P. aridus*
- *P. bleaklyi*
- *P. degeneri*
- *P. diphyllus*
- *P. elegantulus*
- *P. griffinii*
- *P. humilis*
- *P. inflatus*
- *P. metcalfei*
- *P. oliganthus*
- *P. ovatus*
- *P. pruinosus*
- *P. pseudoparvus*
- *P. radicosus*
- *P. rattanii*
- *P. subserratus*
- *P. virens*
- *P. watsonii*
- *P. wilcoxii*

Sect. *Caespitosi* subsect. *Caespitosi*

- *P. abietinus*
- *P. caespitosus*
- *P. crandallii*
- *P. glabrescens*
- *P. harbourii*
- *P. procumbens*
- *P. ramaleyi*
- *P. retrorsus*
- *P. teucrioides*
- *P. thompsoniae*
- *P. tusharensis*

Sect. *Caespitosi* subsect. *Linarioides*

- *P. californicus*
- *P. discolor*
- *P. pinifolius*
- *P. linarioides*
